# EZH2 inhibition promotes tumor immunogenicity in lung squamous cell carcinomas

**DOI:** 10.1101/2023.06.06.543919

**Authors:** Tanner J. DuCote, Xiulong Song, Kassandra J. Naughton, Fan Chen, Daniel R. Plaugher, Avery R. Childress, Abigail R. Edgin, Xufeng Qu, Jinze Liu, Jinpeng Liu, Fei Li, Kwok-Kin Wong, Christine F. Brainson

## Abstract

Two important factors that contribute to resistance to immune checkpoint inhibitors (ICIs) are an immune-suppressive microenvironment and limited antigen presentation by tumor cells. In this study, we examine if inhibition of the methyltransferase EZH2 can increase ICI response in lung squamous cell carcinomas (LSCCs). Our *in vitro* experiments using 2D human cancer cell lines as well as 3D murine and patient derived organoids treated with two inhibitors of the EZH2 plus interferon-γ (IFNγ) showed that EZH2 inhibition leads to expression of both major histocompatibility complex class I and II (MHCI/II) expression at both the mRNA and protein levels. ChIP-sequencing confirmed loss of EZH2-mediated histone marks and gain of activating histone marks at key loci. Further, we demonstrate strong tumor control in models of both autochthonous and syngeneic LSCC treated with anti-PD1 immunotherapy with EZH2 inhibition. Single-cell RNA sequencing and immune cell profiling demonstrated phenotypic changes towards more tumor suppressive phenotypes in EZH2 inhibitor treated tumors. These results indicate that this therapeutic modality could increase ICI responses in patients undergoing treatment for LSCC.

## INTRODUCTION

Lung squamous cell carcinoma (LSCC) is a subtype of non-small cell lung cancer (NSCLC) that historically has limited therapeutic options^1, 2^. The FDA recently approved first-line PD1/PD-L1 targeting immunotherapy for LSCC patients^3^. This therapy blocks the immune-evasion PD1/PD-L1 interaction, allowing for tumor-reactive T cells to expand and destroy the tumor. However, durable responses are seen in only ∼20% of LSCC advanced-stage patients^4^. Efforts to increase the response rates in individuals have focused on combination therapies. EZH2 is a histone methyltransferase that catalyzes histone H3 lysine 27 tri-methylation (H3K27me3), a mark associated with gene silencing^5^. The FDA-approved EZH2 inhibitor tazemetostat^6, 7^, as well as tool compounds including GSK126^8^, are specific EZH2 inhibitors that serve to decrease H3K27me3, de-repress genes, and may lead to improved immunotherapy responses though several mechanisms.

In order to study LSCC in immunocompetent hosts, several autochthonous genetically engineered mouse models have been established. One such model was generated through biallelic deletion of the tumor suppressors *Pten* and *Lkb1* (aka *Stk11*)^9^. Tumors from these mice were shown to have transcriptional similarity to human LSCC, had high expression of PD-L1 on the tumor-propagating cells, and had predominant populations of tumor-associated neutrophils^9^. It is widely believed that tumor associated neutrophils, in particular neutrophils that are sometimes described as granulocytic myeloid-derived suppressor cell (Gr-MDSC), can promote tumor growth by creating a lymphocyte-suppressive microenvironment^10^. Mechanisms through which neutrophils suppress T cells include high expression of arginase, and reactive oxygen species^10^. However, some neutrophils are thought to be tumor eliminating, and can create a pro-lymphocyte environment through production of TNF-α and CXCL10, and antigen presentation^10^. The neutrophil to lymphocyte ratio appears to strongly predict response to immunotherapy, suggesting that in the majority of NSCLCs, neutrophils are T cell suppressive^11^.

One essential mechanism for T cell activation is co-stimulation of antigens presented by <u>m</u>ajor <u>h</u>istocompatibility <u>c</u>omplexes (MHC) I and II. Many tumor cells have evolved to repress antigen presentation machinery in order to evade the immune system surveillance^12^. It has been reported that patient tumors with high expression of MHC II demonstrate greater response to anti-PD-1 checkpoint inhibitors in melanoma^13^. In other cancer types, it has been demonstrated that both MHC I and MHC II can be regulated by the chromatin modifying enzyme enhancer of zeste 2 (EZH2)^12, 14–17^.

Here, we utilized several murine and human models of lung squamous cell carcinoma (LSCC) to understand if and how EZH2 inhibition will boost immunotherapy responses. We found that inhibition of EZH2 catalytic activity with either GSK126 or EPZ6438 in the presence of IFNγ was able to de-repress numerous genes encoding antigen presentation constituents and the pro-T cell cytokines CXCL9/10/11 in both human and murine tumoroids. ChIP-sequencing in human patient-derived tumoroids (PDTs) further delineated the patterns of epigenetic changes in response to EZH2 inhibition and IFNγ treatment. In both autochthonous and syngeneic grafts, EZH2 inhibition alone or with immunotherapy led to excellent tumor control. Single cell RNA-sequencing (scRNAseq) and flow cytometry showed changes consistent with more immunogenic tumor cells and a more pro-T cell tumor microenvironment. Together, these data strongly support the addition of EZH2 inhibition to immunotherapy regimens that have now become first line treatment for many LSCC patients.

## RESULTS

### EZH2 inhibition allows up-regulation of MHC Class I and Class II in multiple models of lung squamous cell carcinoma

The Polycomb Repressive Complex 2 plays a central role in gene repression, including genes involved in immunogenicity^12, 14–17^. In order to understand the tumor-cell intrinsic effects of EZH2 inhibition, we treated four different non-small cell lung cancer cell lines with the EZH2 inhibitors GSK126 or EPZ6438 for 5 days, followed by two days of EZH2 inhibition with IFNγ. We reasoned that this would allow for the de-repression of numerous loci in the cells, and that IFNγ treatment could then activate interferon responsive genes (**Figure 1A**). Analysis of mRNA expression in these cultures showed that the MHC class I genes *B2M* and *HLA-A* were robustly upregulated by IFNγ in all cell lines, and that EZH2 inhibition led to further *HLA-A* up-regulation in three out of four cell lines (**Figure 1B**). The MHC class II genes *CIITA* and *HLA-DRA* showed a similar pattern, with significant stepwise increases in HLA-DRA expression with EZH2 inhibition, IFNγ treatment and combination (**Figure 1B**). We also examined gene expression of *CD274* (encoding PD-L1) and the putative LSCC stem cell marker NGFR (**Supp** **Figure 1A**). Consistent with our previous work^18^, we observed that *NGFR* was up-regulated by EZH2 inhibitor in A549 cells. PD-L1 results show that while IFNγ can reproducibly up-regulate this T cell suppressor, only in the HCC15 cell line does EZH2 inhibition drive additional expression. To examine if the up-regulation in genes led to increased cell surface protein expression, flow cytometry was used. An antibody against the MHC Class I proteins HLA-A,B,C showed a stepwise increase in expression levels, with the highest levels observed in IFNγ and EZH2 inhibitor co-treated cultures (**Figure 1C**). NGFR showed up-regulation at the protein level in 3 cell lines, but not in HCC95 that already expresses very high NGFR levels (**Supp** **Figure 1B**). PD-L1 expression was up-regulated significantly by IFNγ in A549 and HCC15 lines, and again HCC15 was the only cell line for which PD-L1 expression was further boosted by EZH2 inhibition. Strikingly, MHC Class II protein HLA-DR was dramatically increased in the cultures treated with a combination of IFNγ and EZH2 inhibitor, even in the H520 cells in which EZH2 inhibition drove a negligible increase in HLA-DRA mRNA (**Figure 1D**). Lastly, we confirmed changes in B2M and HLA-DR,DQ,DP and efficacy of EZH2 inhibition by H3K27me3 levels by western blotting (**Figure 1E**).

**Figure 1:**
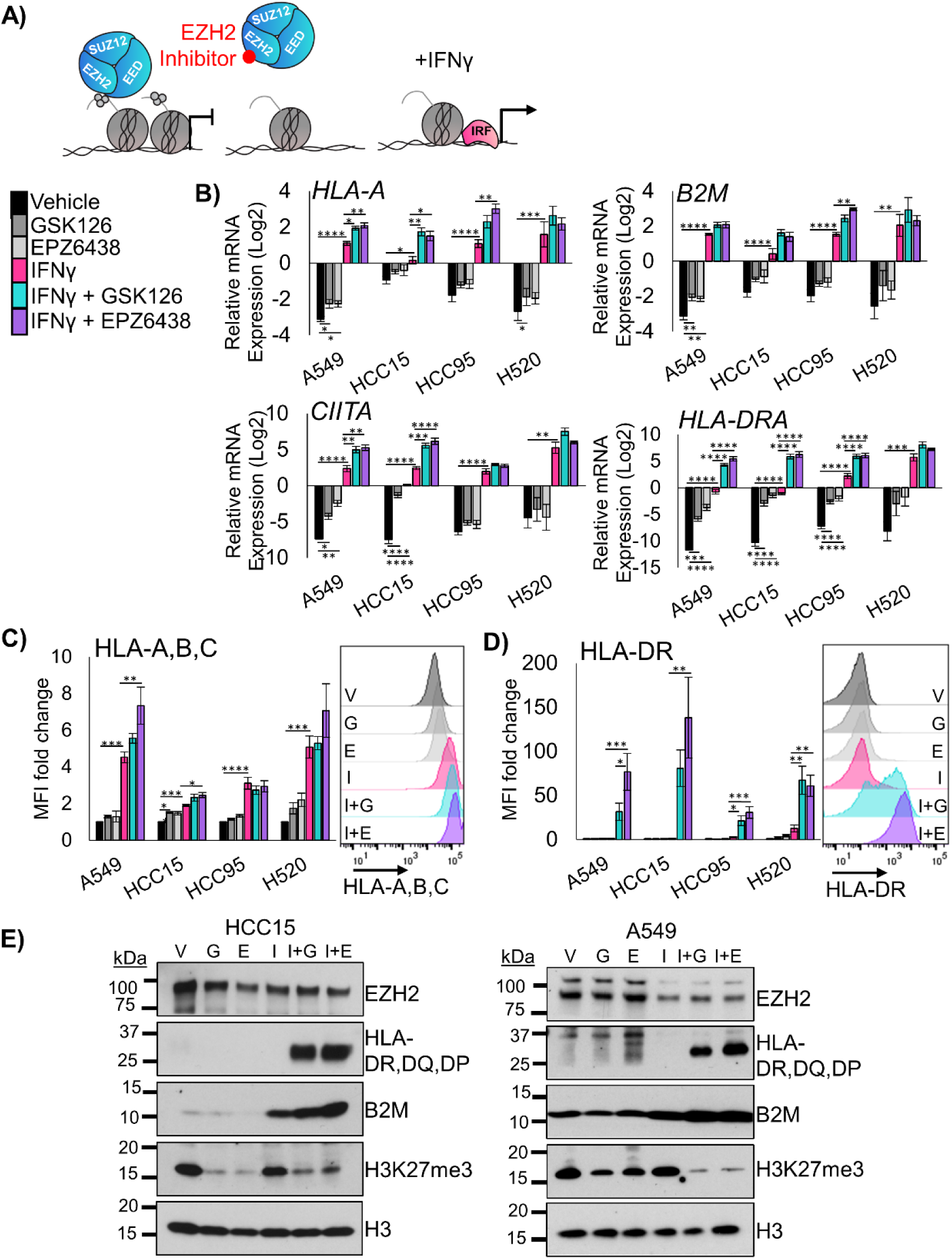
EZH2 inhibition allows up-regulation of MHC Class I and Class II in 2D human LSCC cell lines. **A)** Schematic for proposed mechanism: Inhibition of EZH2 methyltransferase activity by the drugs GSK126 or EPZ6438 will lead to de-repression of antigen presentation genes that can then be more effectively activated by interferon gamma. **B)** RT-qPCR in the indicated four human lung cancer cell lines treated for 7 days with vehicle or EZH2 inhibition with IFNγ added on day 5 for the genes *B2M*, *HLA-A*, *CIITA* and *HLA-DRA*, mean +/-SEM is graphed, n = 4 individual cultures, *indicates p<0.04, **p<0.008 ***p<0.0009, ****p<0.0001 by one-way ANOVA with pairwise comparisons and Holm-Šídák’s *post hoc* test. **C)** Flow cytometry analysis of indicated four human lung cancer cell lines treated for 7 days with vehicle or EZH2 inhibition with IFNγ added on day 5 for the cell surface proteins HLA-A,B,C and HLA-DR, mean +/-SEM is graphed, n = 4 individual cultures, * indicates p<0.04, ** p<0.006, ***p<0.0009, ****p<0.0001 by one-way ANOVA with pairwise comparisons and Holm-Šídák’s *post hoc* test. Representative histograms from HCC15 cell lines are shown, G=GSK126, E=EPZ6438, I=IFNγ, I+G= IFNγ+GSK126 and I+E= IFNγ+EPZ6438. **D)** Western blotting of A549 and HCC15 cell lines treated for 7 days with vehicle or EZH2 inhibition with IFNγ added on day 5 for the proteins B2M, HLA-DR,DQ,DP, EZH2, H3K27me3 and total histone H3. Data are representative of two individual cultures. See also Supplementary Figure 1.

Three-dimensional (3D) cultures allow for growth of tumor cells that cannot proliferate in 2D, and these cultures can retain the epigenetic state of *in vivo* tumors^19^. Therefore, we developed two patient-derived tumoroid (PDT) cultures from distinct LSCC patients. We treated these PDTs with the EZH2 inhibitors GSK126 and EPZ6438 for nine days, followed by two days of EZH2 inhibition with IFNγ. In tumoroids treated with EZH2 inhibitor and IFNγ, B2M and HLA-A mRNA were both increased (**Figure 2A**), as well as *CD274* and *NGFR* gene expression (**Supp** **Figure 2A**). Similarly to the 2D cultures, the most striking results were with *CIITA* and *HLA-DRA* (**Figure 2B**). By flow cytometry, both HLA-A,B,C and HLA-DR have significantly higher expression in tumoroids treated with both EZH2 inhibition and IFNγ (**Figure 2C+D**). Cell surface expression of NGFR and PD-L1 proteins were changed very minimally by the treatments (**Supp** **Figure 2B**). These results in 3D tumoroid cultures and the 2D cultures strongly implicate that de-repression of MHC class II is one of the primary outcomes of EZH2 inhibition in murine and human lung squamous cell carcinomas.

**Figure 2:**
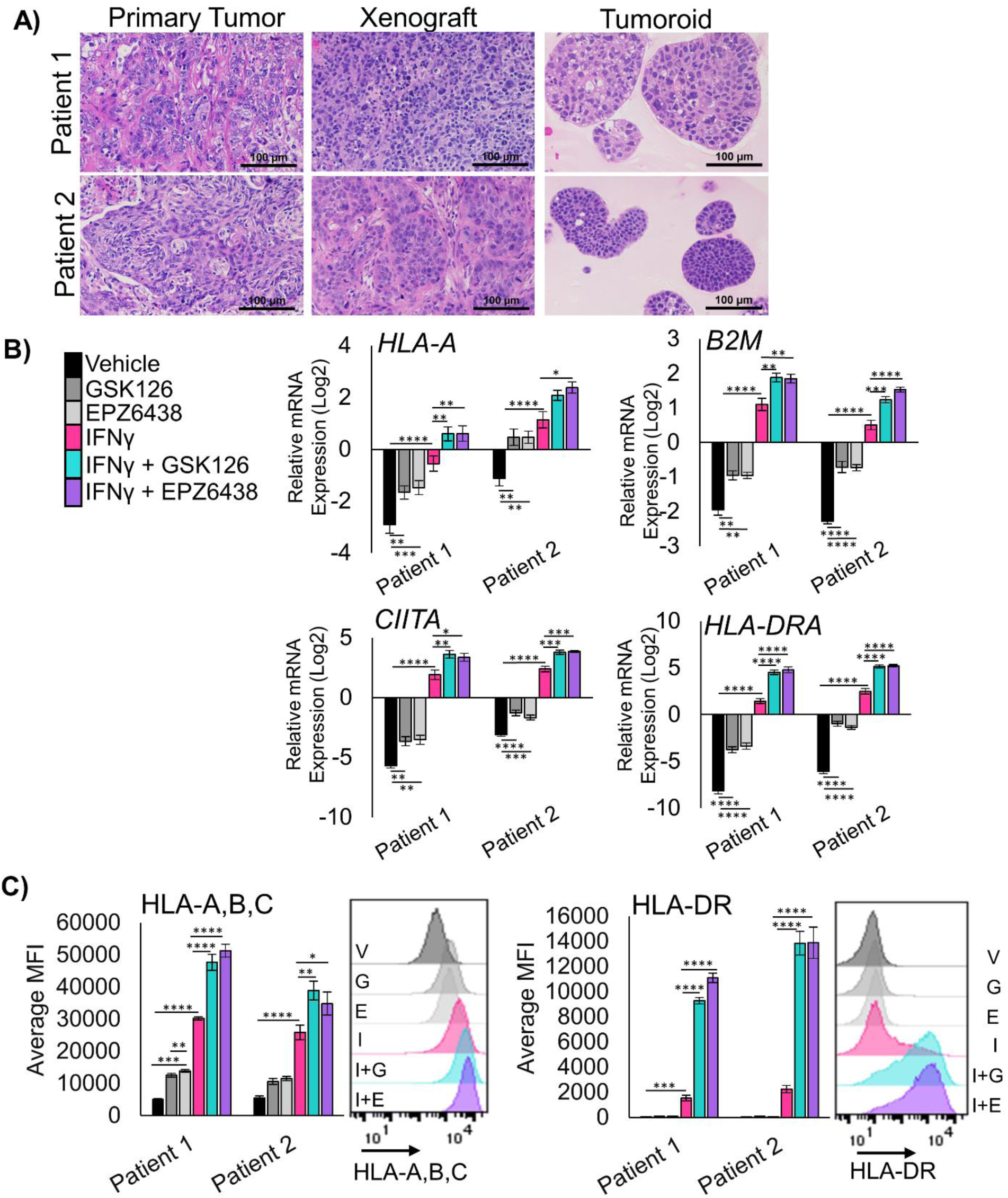
EZH2 inhibition allows up-regulation of MHC Class I and Class II in human LSCC patient-derived tumoroids. **A)** Hematoxylin and eosin staining of primary squamous cell carcinoma tissue, xenograft tissue from primary patient tissue, and tumoroids generated from xenografts, scale bars = 100µm. **B)** RT-qPCR in two unique patient-derived tumoroid cultures treated for 11 days and IFNγ added on day 9 for the genes *HLA-A*, *B2M*, *CIITA*, and *HLA-DRA*, mean +/-SEM is graphed, n = 4, * indicates p<0.03, **p<0.005, ***p<0.0008, ****p<0.0001 by one-way ANOVA with multiple comparisons and Holm-Šídák’s *post hoc* test. **C)** Flow cytometry analysis of both patient derived tumoroids treated for 11 days with IFNγ added in on day 9 for cell surface proteins HLA-A,B,C and HLA-DR, mean +/-SEM is graphed, n=4 biological replicates, * indicates p = 0.043, ** p<0.004, ***p<0.0009, ****p<0.0001 by one-way ANOVA with multiple comparisons and Holm-Šídák’s post *hoc test*. Representative histograms for patient 1 are shown, G=GSK126, E=EPZ6438, I=IFNγ, I+G=IFNγ+GSK126 and I+E=IFNγ+EPZ6438. See also Supplementary Figure 2.

### RNA- and ChIP-sequencing reveal regulation of both MHC and cytokine expression in tumor cells treated with EZH2 inhibitors

To effectively test immunotherapies *in vivo*, hosts with intact immune systems must be used. Genetically engineered mouse models meet this requirement by allowing for tumor formation in the autochthonous setting. We previously reported that conditional bi-allelic deletion of the genes *Pten* and *Lkb1* (aka *Stk11*) with inhaled adeno-Cre virus leads to lung squamous cell carcinoma^9^. From these squamous tumors, we developed tumoroid cultures (**Figure 3A**). Tumoroids treated with IFNγ were able to robustly upregulate all MHC class I and class II genes tested, with further up-regulation of the HLA-A ortholog *H2-K1* and *B2m* with EPZ6438 and IFNγ (**Supp** **Figure 3A**). Similar to human models, *Ngfr* was also up-regulated by EZH2 inhibition. Next, we performed flow cytometry, and observed significant increases in the MHC Class 2 protein I-A/I-E, in cultures treated with IFNγ and EZH2 inhibitor relative to IFNγ alone (**Figure 3B**). Furthermore, both H2-K^d^,D^d^ and PD-L1 expression were highest in tumoroids treated with both EZH2 inhibition and IFNγ.

**Figure 3:**
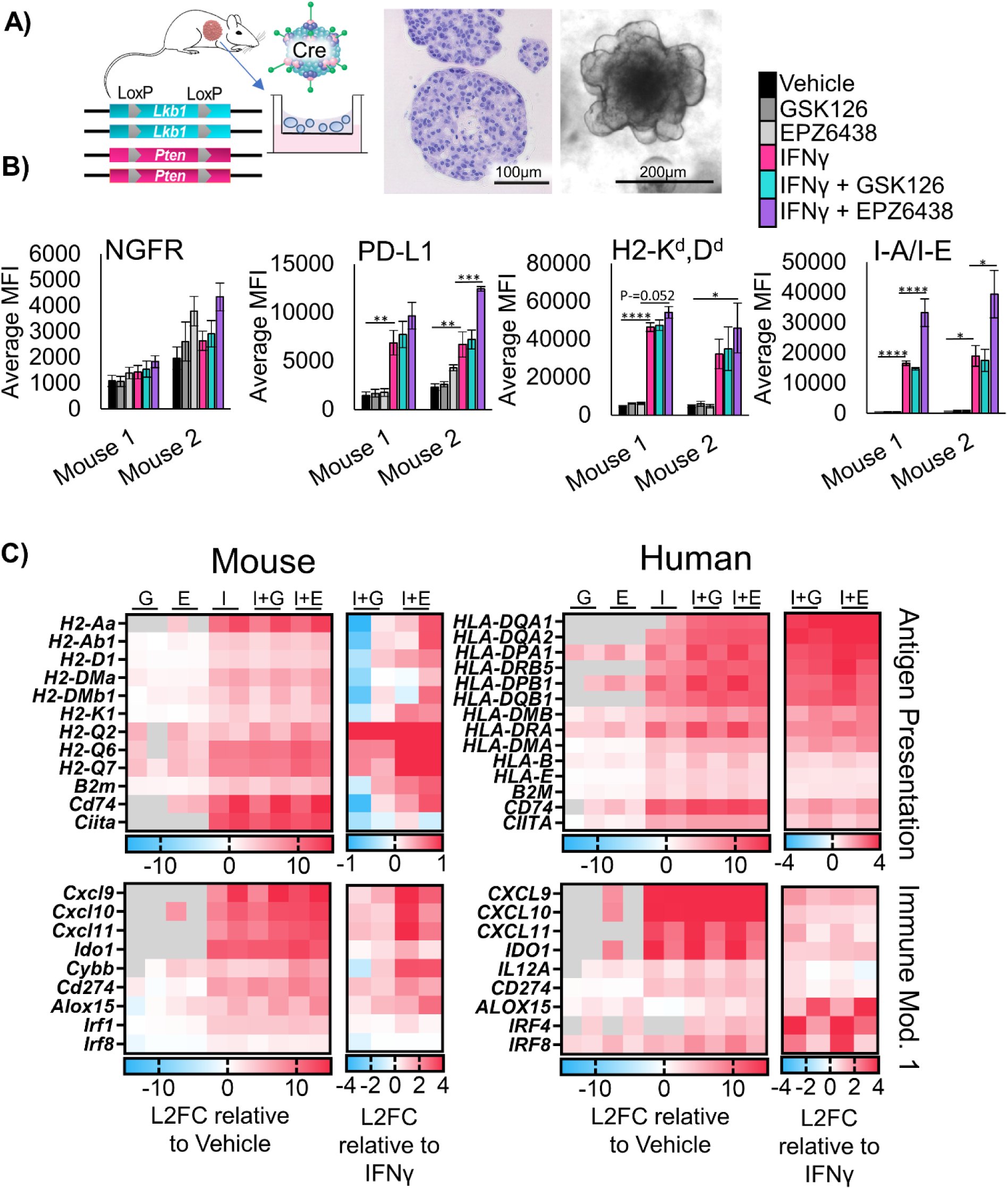
Murine LSCC organoids share de-repression of MHC and pro-T cell cytokines with human models. **A)** Schematic: Generation of murine tumoroids in air-liquid interface from tumor induced in *Lkb1/Pten* mice by adenoCre administration, showing H&E stain of tumoroids, scale bar = 100µm, and brightfield microscopy, scale bar = 200µm. **B)** Flow cytometry analysis of two separate murine tumoroid models treated for 11 days with IFNγ added on day 9 stained for cell surface expression of NGFR, PD-L1, H2Kd,Dd, and I-A/I-E, n = 5 individual experiments except mouse 2 I-A/I-E and PD-L1, n=4 individual experiments, * indicated p<0.031, **p<0.006, ***p=0.0002, ****p<0.0001 by one-way ANOVA with multiples comparisons and Holm-Šídák’s *post hoc* test **C)** Heat maps of Log2-fold change in expression level from patient derived and murine tumoroids treated for 11 days and IFNγ added in on day 9, G=GSK126, E=EPZ6438, I=IFNγ, I+G= IFNγ+GSK126 and I+E= IFNγ+EPZ6438. For each map, the first columns are sample 1, the second columns are sample 2. Expression relative to vehicle control and relative to IFNγ only are depicted. See also Supplementary Figure 3

To further confirm gene programs that are regulated by EZH2 inhibition in a conserved fashion in both mouse and human lung SCC tumoroids, we performed RNAseq on all four tumoroid models. We compared genes up-regulated by IFNγ and EZH2 inhibition relative to vehicle treatment, and genes up-regulated by combination of EZH2 inhibition and IFNγ relative be to IFNγ (**Figure 3C**). In addition to genes involved in antigen presentation, we observed a conservation of up-regulation of the pro-T cell cytokines *CXCL9/10/11* when EZH2 inhibition and IFNγ treatments were combined. In human cells, there was also a down-regulation of the pro-neutrophil cytokines *CXCL1/2/3*, and the cytokines *IL1A* and *IL1B* in response to treatment with EZH2 inhibition and IFNγ. We next performed gene set enrichment analysis (GSEA)^20^ and observed a decrease in MYC targets, E2F targets, and DNA repair gene programs in response to EZH2 inhibition. Additionally, we saw an increase in pathways involved in the inflammatory response and IFN-response in response to EZH2 inhibition, and both effects were maintained when IFNγ was also added (**Supp** **Figure 3B, Supp Table 1**).

To confirm the direct targets of EZH2 inhibition, we performed ChIPseq on the Patient 1 PDT model treated with vehicle, IFNγ, EPZ6438 or a combination of IFNγ and EPZ6438. We assessed enrichment of chromatin bound to H3K27me3, H3K27ac and H3K4me3 histone marks. We observed that IFNγ treatment increased the number of peaks bound by H3K27me3 by 47% and that H3K27me3 peaks were nearly completely ablated by treatment with EPZ6438 (**Figure 4A, Supp** **Figure 4A+B**). Several patterns of epigenetic gene regulation emerged from this analysis. One pattern was observed at MHC Class II genes, and involved a loss of H3K27me3 with EZH2 inhibitor treatment, but only with IFNγ and EZH2 inhibitor together did the loci produce transcript (**Figure 4B**, **Supp** **Figure 4C**). Another pattern was that IFNγ alone increased H3K27me3, H3K27ac, H3K4me3, and gene expression, and EZH2 inhibition with IFNγ boosted gene expression further by reduction of H3K27me3. This happened at loci including the *CXCL9/10/11* cluster (**Figure 4C**). A smaller group of genes, including *ALOX15*, showed up-regulated transcription when EZH2 was inhibited regardless of IFNγ treatment (**Figure 4D**). *ALOX15* is involved in resolving inflammatory states, and therefore, may be able to reduce a pro-tumor inflammatory environment^21^. Lastly, there were genes including *IL1B* that were expressed in the vehicle control cells, and transcriptionally turned off by EZH2 inhibition, despite loss of H3K27me3 at the loci (**Figure 4E**). Given that IL1β is a known driver of myeloid cell recruitment and immunosuppression^22^, a decrease in its expression could lead to more effective T cell responses.

**Figure 4:**
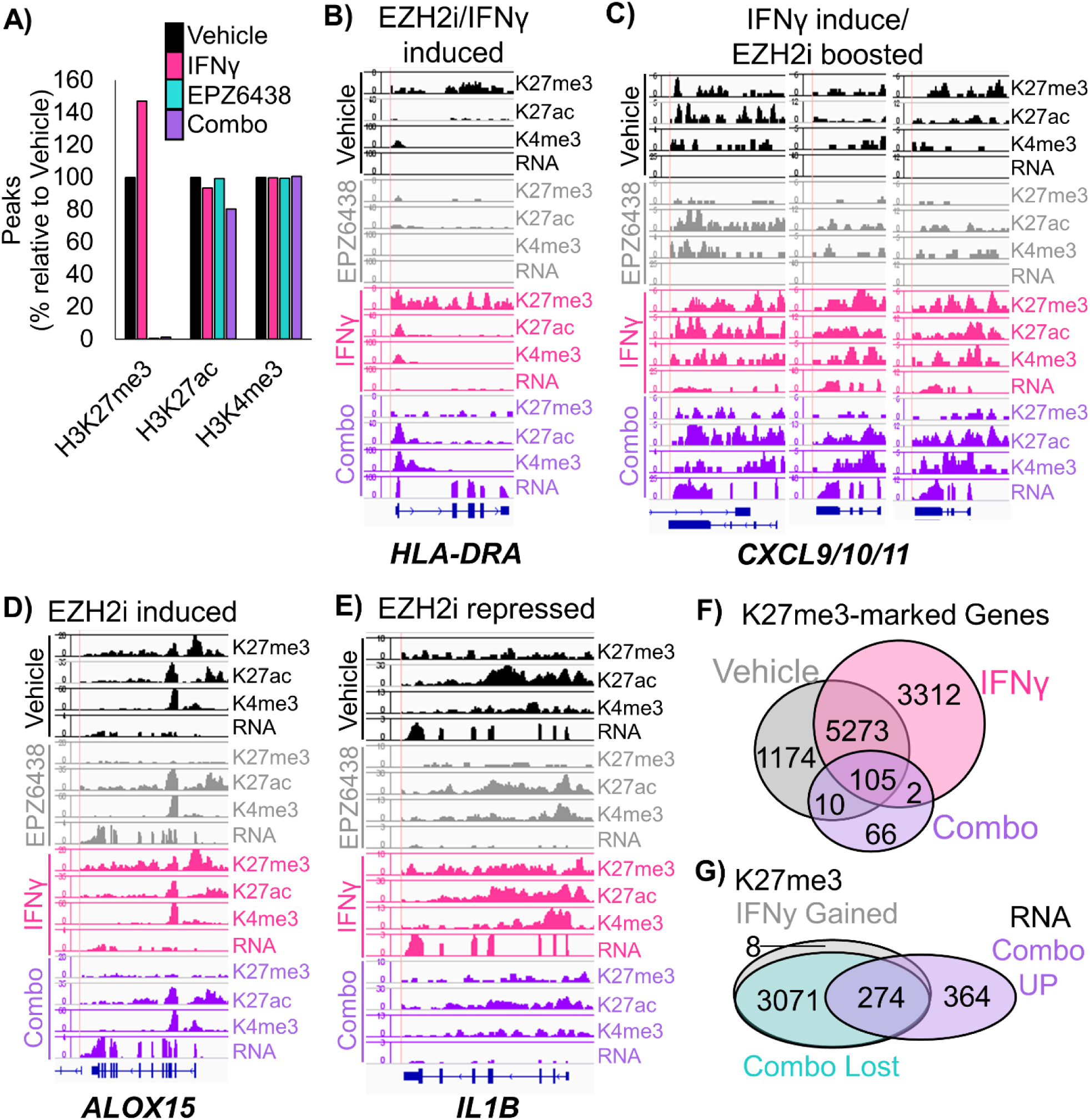
ChIP-sequencing of human patient-derived organoids confirms direct regulation of MHC and pro-T cell cytokines by EZH2. **A)** Peaks called at FDR 1E-7 for ChIP-seq using the chromatin marks H3K27me3, H3K27ac, and H3K4me3 in patient-derived tumoroids from the indicated treatment groups. **B-D)** Wiggle plots for H3K27me3, H3K27ac, and H3K4me3 histone mark enrichments, and matched RNAseq tracks in patient-derived tumoroids from the indicated treatment groups for the genes: **B)** *HLA-DRA* **C)** *CXCL9/10/11* **D)** *ALOX15* **E)** *IL1B* **F)** H3K27me3 peaks were called for each treatment group and GREAT was used to identify associated genes, which were then depicted by Venn diagram. **G)** H3K27me3 peaks that were gained or increased more than 2-fold with IFNγ treatment, and lost with EPZ6438 treatment were linked to associated genes by GREAT. The Venn diagram shows the overlap of these genes with those significantly up-regulated in combination treated vs IFNγ treated tumoroids. See also Supplementary Figure 4.

We were intrigued that ChIPseq analysis revealed a global increase in H3K27me3 in tumoroids treated with IFNγ. In the literature, we could only find one example of this potentiation of PRC2 activity by IFNγ in human macrophages^23^. We first examined peaks and associated genes that gained and lost H3K27me3 in our PDT model. We observed that there were 3729 H3K27me3 peaks corresponding to 3312 genes that were present only in the IFNγ treated cells. There were also 766 H3K27me3 peaks, corresponding to 766 genes, unique to vehicle control cells. These results suggest that H3K27me3 peaks are being both gained and repositioned in the chromatin in response to IFNγ treatment. Next, we assessed both peaks that were gained and those that were increased more than 2-fold, and found 3773 H3K27me3-enriched peaks corresponding to 3351 genes in IFNγ treatment compared to vehicle (**Figure 4F**). Of these genes, the majority (99%) did not change in gene expression in IFNγ compared to vehicle cells, while 33 genes were up-regulated and 3 genes were downregulated in RNA expression. Intriguingly, 989 of these genes (30%) also gained H3K27ac marks with IFNγ treatment, suggesting the PRC2 activity was directed to these genes to help repress the IFNγ response. When EZH2 inhibition was combined with IFNγ, of the 3773 peaks that gained H3K27me3 with INFγ treatment, all but 8 peaks were lost (**Figure 4G**). In the combination treatment, of the 638 genes that were significantly up-regulated compared to IFNγ treatment in these same tumoroids, 43% (274) of the genes had gained H3K27me3 peaks with IFNγ treatment and lost those peaks with combo treatment, with 32% (206) of the genes also gaining H3K27ac peaks. The major pathways for the 274 genes regulated in this fashion included Inflammatory responses, Cell adhesion and signaling, TP53 and apoptosis, Nervous system and Polycomb targets (**Supp Table 2**). Together these data indicate that IFNγ treatment increases Polycomb-mediated gene repression, which can be relieved by treatment of cells with EZH2 inhibitor and drive cells to become more immunogenic.

### Treatment of lung squamous cell carcinoma tumor-bearing mice with EZH2 inhibitor and anti-PD1 results in strong tumor control

To study the effects of EZH2 inhibition in combination with immunotherapy *in vivo*, we induced tumors to grow in the lungs in *Lkb1/Pten* mice by adeno-Cre inhalation and randomized mice onto four treatment arms. We treated mice for 4 weeks with magnetic resonance imaging of the thoracic cavity at baseline, 2 weeks, and 4 weeks to quantify lung tumor burden (**Figure 5A**). Consistent with *LKB1* mutation predicting poor response to immunotherapy^15–17^, anti-PD1 treatment alone had only a small impact on tumor growth (**Figure 5B**). In contrast, treatment of mice with the EZH2 inhibitor GSK126 showed excellent tumor control with some tumor regression, and treatment with EZH2 inhibitor and anti-PD1 lead to significant tumor regression in all mice tested. Although these results were exciting, the experiments did not use the newly FDA-approved EZH2 inhibitor tazemetostat. In order to test tazemetostat in a faster and more cost-effective system, we established a syngeneic graft model by injecting *Lkb1/Pten* tumoroids into the parental mouse strain. By nuclear phenotyper, we observed that the tumor from this graft model closely resembled those in the autochthonous model, including the predominant infiltration of neutrophils (**Figure 5C**). After 14 days of treatment, tumors treated with EPZ-6438 alone had not increased in volume, and combination treated tumors grew initially, but began to regress at day 9, and ended with lower tumor volumes than EZH2 inhibition alone (**Figure 5D**).

**Figure 5:**
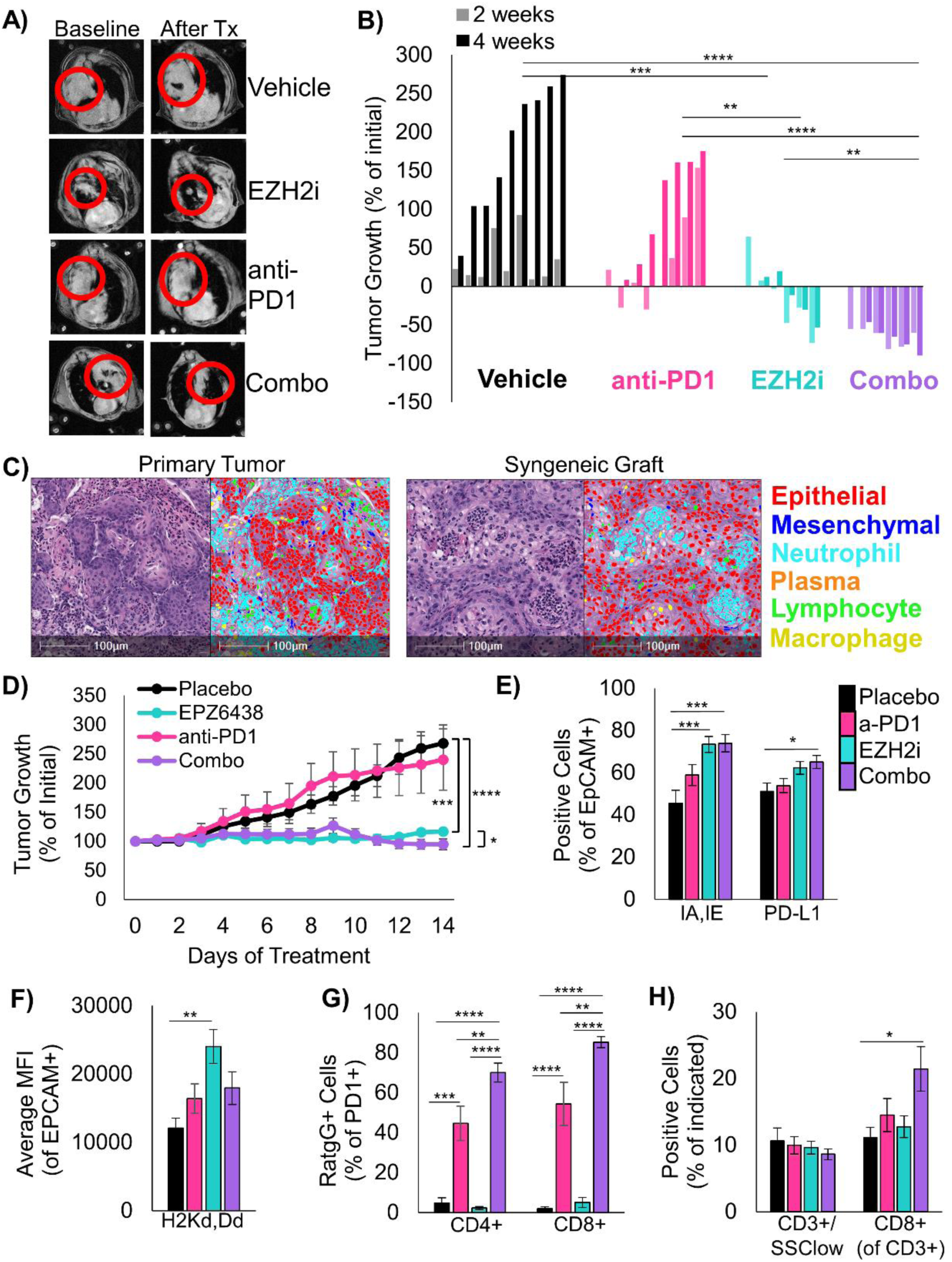
EZH2 inhibition combined with immunotherapy is extremely effective in mouse models of LSCC. **A)** Representative MRI scans of autochthonous mice from each treatment arm at baseline and after treatment. **B)** Waterfall plot showing change in tumor volume for each mouse on all treatment arms, **** indicates p<0.0001, *** indicates p<0.001, ** indicates p<0.01, * indicates p<0.05 by one-way ANOVA with multiple comparisons and Holm-Šídák’s *post hoc* test on log2-transformed values. **C)** H&E and HALO nuclear phenotyper images showing the cells within an autochthonous *Lkb1/Pten* tumor and a syngeneic graft seeded from *Lkb1/Pten* tumoroids. **D)** Percentage tumor growth from the syngeneic mouse model during 14 days of indicated treatments. *** indicates p=0.0004, **** p<0.0001 by one-way ANOVA with multiple comparisons and Holm-Šídák’s *post-hoc* test, * indicates p=0.042 by two-tailed t test on log2-transformed values, Mice/tumors n are placebo=4/8, EPZ6438=5/8, anti-PD1=6/8, combo=5/9, mean +/-s.e.m. is plotted. **E)** Flow cytometry analysis of dissociated tumors from the syngeneic grafts from the indicated treatment arms at day 14. Percentage of EpCAM+ cells expressing IA/IE or PD-L1 are graphed, mean +/-s.e.m. is plotted, placebo n=7, EZH2 inhibitor n=7, anti-PD1 n=8, combo n=7 with 2 experimental replicates each, * indicates p=0.035, *** p=0.0008 by one-way ANOVA with multiple comparisons and Holm-Šídák’s *post-hoc* test. **F)** From the same tumor grafts, MFI for HLA-A in the EpCAM+ cells is graphed, mean +/-s.e.m. is plotted, n=7, EZH2 inhibitor n=7, anti-PD1 n=8, combo n=7 with 2 experimental replicates each, ** indicates p=0.0015 by one-way ANOVA with multiple comparisons and Holm-Šídák’s *post-hoc* test. **G)** From tumor grafts, PD1+/CD3+/CD4+ cells and PD1+/CD3+/CD8+ were gated and percentage of cells bound to Rat-IgG2A antibody are graphed, mean +/-s.e.m. is plotted, n=6, EZH2 inhibitor n=6, anti-PD1 n=7, combo n=7, ** indicates p<0.006, *** p=0.0001, **** p<0.0001 by one-way ANOVA with multiple comparisons and Holm-Šídák’s *post-hoc* test. **H)** From the grafts, percentage of CD3+/SSClow cells within the CD45+ fraction and percentage of CD8+ cells withing the CD3+ fraction were graphed, please see Supp Figure 5C for representative gates, n=8, EZH2 inhibitor n=8, anti-PD1 n=9, combo n=7 with 2 experimental replicates each, * indicates p=0.0197, **** p<0.0001 by one-way ANOVA with multiple comparisons and Holm-Šídák’s *post-hoc* test. See also Supplementary Figure 5.

The marked efficacy of EZH2 inhibition alone in both the autochthonous model and graft models were somewhat surprising, and lead us to compare to EPZ6438 response *in vitro*. By normalized cell counts, there were significant reductions in cultures treated with EPZ6438 and IFNγ compared to vehicle control only in Mouse 2 and Patient 2 (**Supp** **Figure 5A**). Furthermore, for the Mouse 1 tumoroids used for the *in vivo* grafts, *in vitro* data suggested that EPZ6438 itself has no effect on ATP production (**Supp** **Figure 5B**), which implies that the tumor control seen *in vivo* in response to EPZ6438 alone is due almost entirely to immune response. In order to better understand the cell types present, grafts from each treatment arm were dissociated and analyzed by flow cytometry for a panel of immune cell markers (**Supp** **Figure 5C**). Consistent with our *in vitro* data, tumors in mice treated with both EZH2 inhibitor and anti-PD1 had a significant increase in MHC Class II I-A/I-E positive cells, with a smaller less significant increase in PD-L1 positive cells (**Figure 5E**). Tumor cells also showed higher level of H2-K1, particularly in the EZH2 single agent treated tumors (**Figure 5F**). With this flow cytometry panel, we were also able to validate the binding of the anti-PD1 antibody to PD1 positive CD4 and CD8 positive cells. Interestingly, T cells in the EZH2 inhibitor with anti-PD1 treated mice had higher percentages of PD1+ cells bound to antibody than anti-PD1 only treated mice (**Figure 5G**). Although total T cell proportions were not different, there was a significant increase in CD8+ T cells in combination treated mice compared to placebo (**Figure 5H**). There was also a trend towards fewer CD11B+/Ly6G+ or Ly6G+/F4/80-tumor-associated neutrophils, but these differences did not reach significance. One population that was significantly increased in the combination treated mice relative to all other cohorts was a Ly6G+/F4/80+ myeloid cell, and this could indicate a conversion of neutrophils into macrophages by EZH2 inhibition which is a phenomenon described in both human and mouse^24, 25^(**Supp** **Figure 5D**).

### Single cell RNA-sequencing confirm mechanisms through which EZH2 inhibition drives increased tumor immunogenicity

Finally, to assess the transcriptional heterogeneity of cell types in the tumors and how transcriptional programs were changed by treatment, we performed single cell RNA sequencing (scRNAseq). We analyzed fresh autochthonous lung tumors after four weeks of therapy, and also analyzed lungs of mice that were on placebo or EZH2 inhibition that had no tumors as controls. From the tumors and total lungs, we identified 16 unique cell populations that we annotated based on conserved markers (**Figure 6A, Supp Table 3**). Analysis of cell proportions in each treatment group revealed significant decreases in tumor cells and some macrophage and neutrophil populations, and significant increases in T cells, cycling cells, normal lung, and some neutrophil populations (**Figure 6B**). Next, we analyzed the differentially expressed genes in three major groups, the tumor cells (malignant epithelial cells), the macrophage and dendritic cells, and the neutrophils. By GSEA, we observed a dramatic decrease in protein synthesis pathways in tumors cells from combination treatment compared to either single treatment, and increases in DNA pathways, which could reflect cell cycle arrest or DNA damage. Pathways involved in oxidative phosphorylation were upregulated in all three cell populations, and pathways involved in interferon signaling were up-regulated in the tumor cells and the macrophage/dendritic cells with combination treatment (**Figure 6C, Supp** **Figure 6A, Supp Tables 4+5**). To understand the individual genes changed by treatment, we also assessed differential expression of each cell type in the treated groups relative to placebo (**Figure 6D**). Tumor cells that were treated with EZH2 inhibitor alone or in combination with anti-PD1 had increased *H2-K1*, *B2m*, and *Ifngr1* expression, and decreased expression of neutrophil-recruiting chemokines such as *Cxcl3*, *Cxcl5*, and *Ppbp* (aka *Cxcl7*). Moreover, macrophage and dendritic cells had increased MHC class II and IFN-response genes, while neutrophils showed upregulation in IFN-response genes in the combo group relative to both EZH2 inhibition and anti-PD1 treatments alone.

**Figure 6:**
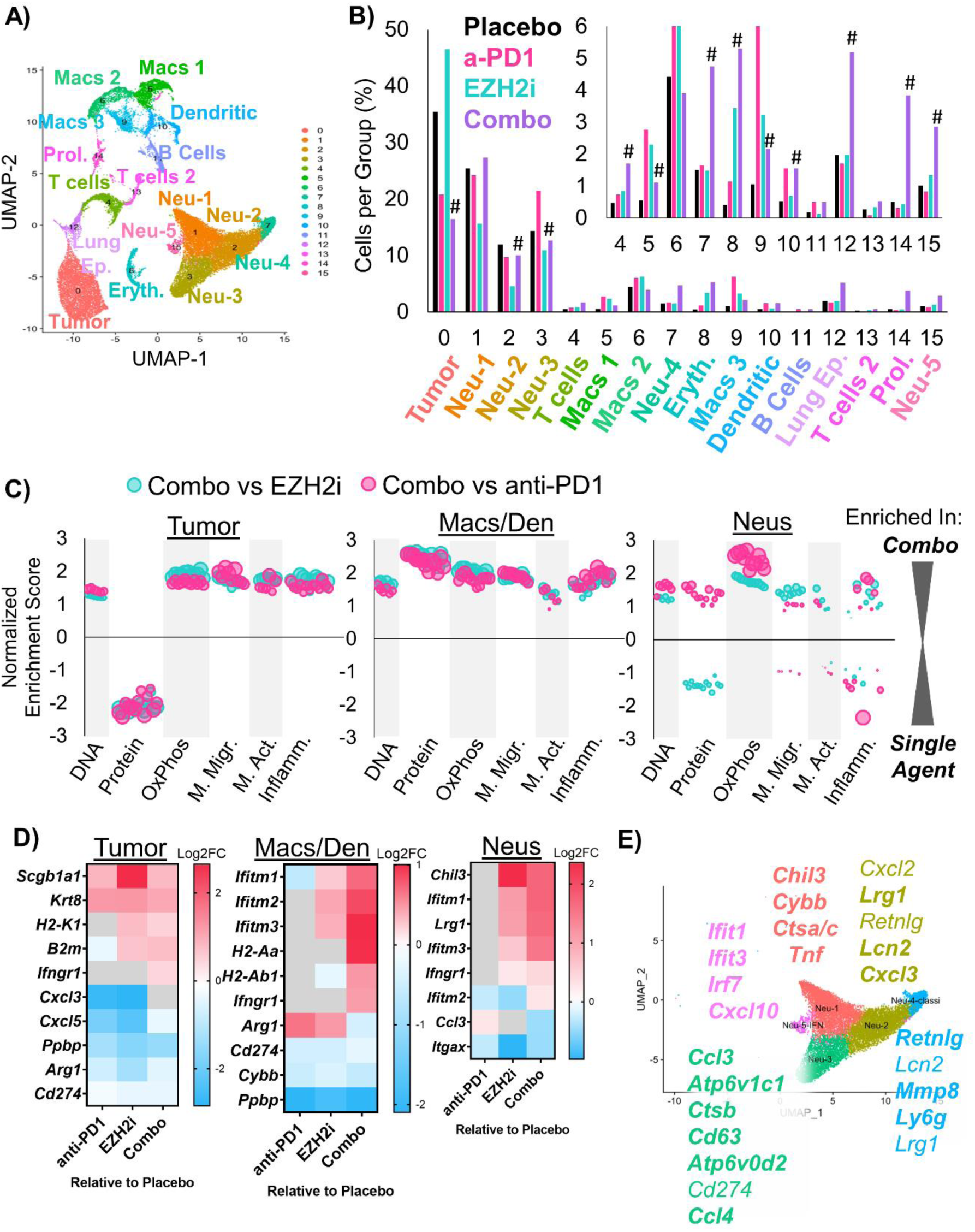
Single Cell RNA sequencing highlights neutrophil heterogeneity shifts in response to EZH2 inhibition combined with immunotherapy. **A)** Annotated Uniform Manifold Approximation and Projection (UMAP) plot showing the 16 different populations within lung tumors of the *Lkb1/Pten* model of LSCC after treatment with placebo, GSK126, anti-PD1, or combined GSK126 with anti-PD1. **B)** Percentage of cells per treatment group graphed for all populations, # indicates adjusted p<0.0012 by proportion z-test. **C)** Gene Set Enrichment Analysis depicting gene sets that are enriched or depleted in Tumor, Macrophage/Dendritic Cells, or Neutrophils in mice treated with EZH2 inhibitor and anti-PD1 contrasted with either treatment alone. Normalized Enrichment Scores are plotted and bubble sizes estimate false discovery rate. See also Supp. Table 2. D) Heat maps showing differentially expressed genes among tumor, macrophages and dendritic cells, and neutrophils between GSK126, anti-PD1, and combination treated mice compared to placebo on the log2 fold change. **E)** UMAP of 5 neutrophil populations showing selected genes that are highly expressed in each cluster. See also Supplementary Figure 6.

It was surprising that the neutrophil populations remained predominant in tumors treated with EZH2 inhibition and immunotherapy, given the large amount of data suggesting that neutrophils prevent proper immunotherapy response. Therefore, we next interrogated the differential gene expression profiles of the five identified neutrophil populations (**Figure 6D, Supp Table 6**). Based on the literature^10^, we believe that there are three populations of neutrophils in our data set that appear to promote tumor elimination. These three populations, Neu1, Neu4, and Neu5, express genes such as *Tnf*, *Cxcl10*, and multiple IFN-response genes. Interestingly, we observed increases in these populations in the group treated with the combined therapy. Moreover, we saw significant decreases in Neu2 and Neu3 populations, which we believe to be immunosuppressive. These two populations expressed genes *Cxcl3*, *Ccl3*, *Ccl4, Atp6v1c1*, *Atp6v0d2*, which have been associated with a tumor promoting phenotype.

However, Neu3 also has an MHC class II antigen presentation gene (*H2-Eb1*), suggesting it may be a ‘hybrid’ tumor promoting and tumor eliminating phenotype. Demonstrating a conservation among models, genes bolded in our figure correspond to genes identified by other groups in neutrophil populations from lung cancers by scRNAseq^26–28^.

In order to understand if neutrophils were also altered in the bone marrow when an EZH2 inhibitor was administered, we performed scRNAseq on the bone marrow of the same mice used for the tumor analysis (**Supp** **Figure 6A**)^29^. Using markers from a recent scRNAseq report on neutrophil development, we observed that Neu1 is the most mature, with Neu6+7 containing myelocytes and pro-myelocytes (**Supp** **Figure 6B**). Our Neu3 population was enriched for expression of *Mmp8* and *Retnlg*, which corresponded to band cells in the previous manuscript. Using this information, we marked Neu1+2 as mature and hypersegmented, Neu3 as banded, BM-Neu4 as meta-myelocyte, Neu5 as myelocyte, and Neu6-8 as pro-myelocytes. By proportion analysis, significantly more neutrophils were mature, and significantly fewer neutrophils were in the meta-myelocyte stage in mice treated with EZH2 inhibitor alone or in combination with anti-PD1 compared to placebo-treated mice (**Supp** **Figure 6C**). To validate this result, we also examined nuclear morphology of the populations by cytospin and again observed more mature and fewer myelocyte cells in EZH2 inhibitor-treated mice compared to placebo-treated mice (**Supp** **Figure 6D**). Similarly, bone marrow cells isolated from the syngeneic graft-bearing mice treated with EZH2 inhibitor were more apoptotic, suggesting a more mature phenotype (**Supp** **Figure 6E**). Together these data show a systemic shift in neutrophil identity away from an immature myeloid suppressor cell phenotype towards a more mature tumor eliminating phenotype, and suggests that neutrophils can be compatible with immunotherapy in the right contexts.

## DISCUSSION

Here, we demonstrate that inhibition of EZH2 in LSCC can boost immunotherapy in several ways (**Figure 7**). In both 2D and 3D *in vitro LSCC* models, treatment with EZH2 inhibition and IFNγ led to increases in MHC I/II and pro-inflammatory cytokine expression. In 3D patient-derived tumoroids, ChIPseq confirmed a switch from repressive to active chromatin at these genes in response to treatment. Next, we employed both autochthonous and syngeneic models of LSCC driven by biallelic deletion of the tumor suppressors *Lkb1* and *Pten*. We observed significant tumor control in both the anti-PD1 with EZH2 inhibitor combination, as well as EZH2 inhibition alone. Using scRNAseq and immune cell profiling, we identified increases in MHC I/II expression and a shift towards tumor-eliminating neutrophils within tumors. Studies have found that *IFNG*, *CXCL9*, and *CD274* expression in tumor specimens correlate to stronger immunotherapy responses^30^. In this study, we found that the pro-T cell cytokines *CXCL9/10/11* are strongly induced by a combination of IFNγ and EZH2 inhibition in both human and murine tumoroids, and that this gene cluster is a direct PRC2 target in human lung cancer cells. Our data mirror those seen in urothelial cells^14^. In addition, *Arg1* was significantly downregulated in autochthonous tumors treated with EZH2 inhibition. Lastly, we found that neutrophil populations shifted towards IFN-responsive and TNFα expressing populations. These data point to multiple overlapping mechanisms through which T cells can be recruited to tumors to turn ’cold’ tumors ’hot’.

**Figure 7:**
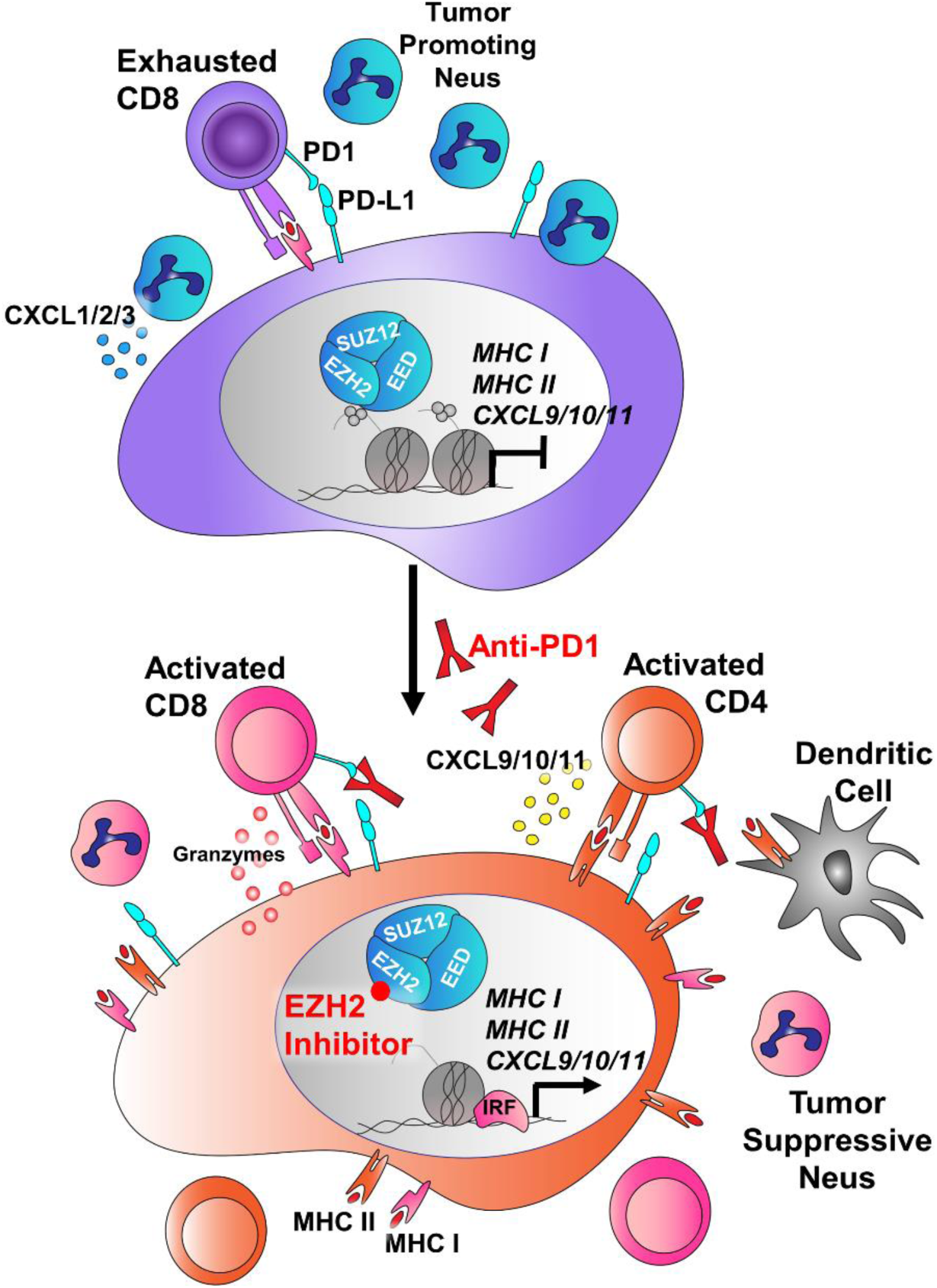
Schematic of tumor cell-intrinsic and microenvironmental consequences of EZH2 inhibition that boost immunotherapy response in LSCC. In LSCC, tumors can evade the immune system through expression of PD-L1, inhibiting T cell activation. In addition, these tumors secrete high levels of *CXCL1/2/3* (mouse orthologs *Cxcl3/5/7*) that attract T-cell suppressive neutrophils, and can express high levels of Arginase, that further drive T cell suppression. In response to EZH2 inhibition, the tumors up-regulation MHC Class I and Class II antigen presentation machinery, and switch from expression of *CXCL1/2/3* to expression of the T cell promoting cytokines *CXCL9/10/11* and the inflammatory resolution molecule *ALOX15*. *IL1B* and *Arg1* are downregulated, and the neutrophils surrounding the tumor take on more tumor-repressive phenotypes. When anti-PD1 antibody is added, the net result is tumor regression through immune targeting of tumor cells.

In addition to strongly influencing the tumor immune microenvironment, EZH2 inhibition drove dramatic increases in expression of both MHC class I and class II. Several studies have also observed that EZH2 plays a major role in repression of MHC class I in head and neck, small cell lung cancer and melanoma^15–17^, and MHC class II in urothelial and AML cancers^14, 31^. Furthermore, MHC class II expression has proven to correlate well with increased ICI response rates in melanoma, and depletion of MHC class II in a NSCLC can covert tumors from ICI sensitive to resistant^13, 32^. While anti-PD1 ICI is thought to activate CD8+ T cells to target the tumor through MHC class I interactions, CD4+ T cells can kill MHC class II+ cells though FAS/FASL interactions. Again, these data suggest not one but multiple mechanisms through which EZH2 inhibition can alter both tumor cells and immune cells for a net effect of increased tumor immunogenicity. This multi-mechanism aspect may help to delay or prevent acquired resistance phenotypes that occur readily when only one molecular pathway is altered. Furthermore, our data suggest an efficacy of EZH2 inhibition as a single therapy in certain instances, and this observation could be used to test EZH2 inhibition as a single agent clinically in LSCC patients who cannot tolerate immunotherapy.

Importantly, the model we used is deficient for LKB1, which is correlated with immunotherapy resistance^33–35^. However, the majority of data are in non-squamous histology tumors and may rely upon KRAS mutational status^36^. Our data suggest that when a tumor is squamous in epigenetic state, that EZH2 inhibition combined with immunotherapy will be an effective treatment approach regardless of tumor genotype. What is less clear is whether tumors that are adenocarcinoma in histology will respond as effectively, and therefore more research is needed. MHC class I and class II expression are intrinsic properties of specific alveolar and bronchiolar lung cells^37, 38^. Therefore, these molecules may already be expressed in adenocarcinomas, depending on the tumor cell-of-origin. We also focused on models with significant neutrophil infiltration. Our recent study of human lung cancers showed that macrophages and plasma cells can also predominate in tumors^39^, and it will be important to test EZH2 inhibition in the context of diverse tumor-immune microenvironments moving forward.

One aspect of immunotherapy response that was not addressed in the current study is the role of tumor mutation burden and neo-antigens. In general, genetically engineered mouse models are thought to have relatively low TMB^40^. However, studies have suggested that EZH2 inhibition can increase genotoxic stress in other models^41^, and this idea should be explored in lung cancer. The effect of EZH2 inhibition in models with varied TMBs would also be worthwhile. Despite this limitation, this work demonstrates several mechanisms through which EZH2 inhibition can boost ICI responses in lung cancer. The EZH2 inhibitor tazemetostat was recently FDA approved, and is already in clinical trials with ICIs for urothelial tumors (NCT03854474). Importantly, these data suggest that EZH2 inhibition plus ICI, or EZH2 inhibition alone could be viable options specifically for lung SCC, and these data serve as strong premise for clinical investigation. Lastly, the model systems we have characterized will be useful tools to explore further mechanisms of ICI response and resistance.

## METHODS

### Animal work

The studies described in this manuscript employ genetically engineered mouse models of squamous lung cancer through the biallelic deletion of the genes *Lkb1 and Pten.* Both alleles are flanked by loxP sites and are deleted by Cre-recombinase administration. Experimental animals are inoculated with 2.9-5 x 10^7^ PFU adeno-CMV-Cre (University of Iowa) through intranasal instillation and are monitored for tumor burden via magnetic resonance imaging (MRI) 40 weeks post infection. Once tumors were detected, animals were randomized and placed onto one of four treatment arms. GSK126 was formulated by adding solid GSK126 (MedChem Express/Xcess Bio) to Captisol®, chopped very finely, then added to sterile saline pH 4.6, and the solution was sonicated. Anti-PD1 clone RMP1-14 (BioXCell #BP0146 or #BE0146) and IgG2a isotype control (BioXCell #BP0089) were diluted with InVivoPure pH 7.0 dilution buffer (BioXCell #IP0070) or InVivoPure pH 6.5 dilution buffer (BioXCell # IP0065) to a concentration of 1.25 μg/μL. Animals were administered GSK126 at 300mg/kg i.p. twice per week, 5-7mg/kg anti-mouse PD1 i.p. immunotherapy three times per week, or a combination of both GSK126 and anti-PD1. Placebos used for treatment arms were Captisol® in sterile saline and rat IgG2a isotype antibody. Treatment regimen was 4-6 weeks and animals were monitored for tumor burden every two weeks via MRI. To prepare formulation of EPZ6438 (MedChem Express #HY-13803/CS-1758), EPZ6438 was added to a solution of 0.1% Tween 80 and 0.5% sodium carboxymethylcelluose and the solution was sonicated. EPZ6438 was administered 250mg/kg by gavage twice daily for 14 days. All experiments were approved by the Dana-Farber Cancer Institute or University of Kentucky Institutional Animal Care and Use Committees (IACUCs).

### Cell lines

Cell lines were maintained according to University of Kentucky biosafety guidelines. All human cell lines were cultured in RPMI 1640 (Gibco, #11875-093), supplemented with 8% fetal bovine serum (VWR), penicillin/streptomycin (Gibco, #15140-122), and 4mM GlutaMAX™ (Gibco, #35050-061) at 37°C and 5% CO_2_. All cell lines were tested regularly for mycoplasma with MycoAlert™ PLUS Mycoplasma Detection Kit (Lonza) and prophylactic treatment with Plasmocin (InvivoGen, #ant-mpt) was used routinely. Cell lines were verified by STR analysis with CellCheck9 by IDEXX laboratories before beginning of experiments and used within 10 passages of authentication.

### Flow cytometry analysis and sorting

For all 2D flow cytometry experiments, cells were trypsinized from culture plates and incubated with antibodies at 1:100 dilution in PBS + 10% FBS (PF10) for 15 minutes at room temperature. The cells were then resuspended in 300μL of PF10 + 4’,6-diamidino-2-phenylindole (DAPI) (1:250). Antibodies for human cell analysis: EpCAM-FITC (BD Biosciences, #347197), PD-L1-PE (eBiosciences, #12-5983-42), CD49f-Alexa Fluor® 647 (BD Biosciences, #562473), NGFR-PECy7 (BioLegend, #345110), CD49f-FITC (Invitrogen, #11-0495-82), HLA-DR-APCCy7 (BioLegend, #307618), and HLA-A,B,C-APC (BioLegend, #311410). Antibodies used for mouse studies: I-A/I-E-PerCP-Cy5.5 (BioLegend #107626), H-2K^d^/H-2D^d^-Alexa Fluor® 647 (BioLegend #114712), NGFR (Cell Signaling #8238S), anti-Rabbit-FITC (Millipore #F2765), PD-L1-PE (BioLegend #124308), Sca1-APCCy7 (BioLegend #108126), EpCAM-PECy7 (BioLegend #118216), CD49f-FITC (Invitrogen #11-0495-82). All antibodies were bound at room temperature for 10 minutes at a dilution of 1:100, with the exception of Sca1 that is used at 1:50. Syngeneic grafts were dissected, minced, dissociated with collagenase/dispase (SIGMA 10269638001, 6mg/mL) for 45 minutes at 37°C and filtered through 40µm cell strainers. Antibodies used for staining were: anti-PD1-BV421 (BioLegend #135218), anti-CD8-BV711 (BioLegend #100747), anti-CD4-BV786 (BioLegend #100552), anti-CD45-FITC (BioLegend #103108), anti-CD3-BB700 (BioLegend #100328), anti-Rat-IgG2A-PECF594 (BioLegend #405432), Ly6G-BV786 (BioLegend #127645), F4/80-PECF594 (BioLegend #123146), CD11b-PECy7 (BioLegend # 101216) with the live-dead stain Zombie UV™ (BioLegend 423107). AnnexinV staining was performed with the Biolegend AnnexinV-FITC, 7AAD kit (#655942) on bone marrow samples (described below) according to the manufacturer’s instructions.

### BM histopaque gradient, Cytospin and Staining

Following euthanasian, mouse legs were removed at the acetabulum and placed into magnesium/calcium free PBS (Cytiva, #SH30256.02). The bones were cleaned and flushed into a 1.5ml tube using magnesium/calcium free PBS. The remaining cells were pelleted via pulse centrifugation then were subject to red blood cell lysis in 250µl of red blood cell lysis. Following, red blood cell lysis the cells were washed with 1ml of calcium/magnesium free PBS then pelleted with pulse spin centrifugation. Cells were then plated for apoptosis at 200,000 cells in 12 well plates in DMEM/F12 media (ThermoFisher, 11330032) with an additional 4 mM Glutamax (ThermoFisher, 35050061), 5 µg/mL ITS (insulin/transferrin/selenium, SIGMA, I3146) and 8-9% fetal bovine serum (VWR, 97068-085). Remaining cells were resuspended in 1 ml of calcium/magnesium free PBS. For Histopaque gradient, in a 5mL flow cytometry tube, 1.5ml of Histopaque 1119 (SIGMA #RNBK6705) was added at room temperature and 1.5ml of Histopaque 1077 (SIGMA #RNBL3022) at room temperature was carefully pipette on top. Samples were then added on top of the histopaque layers slowly to persevere the interface between the layers. The tubes were then spun at 25°C at 1000x g for 25 minutes with no brake. Following centrifugation, the cloudy interface was collected into a 1.5mL tube and washed with 1ml of magnesium/calcium free PBS. The cells were resuspended in 1mL of magnesium/calcium free PBS and 150uL was taken to perform cytopsin. For cytopsin, 150µL cell suspension was pipetted into funnel and slides were spun at 550 rpms for 1 minute. The areas where cells had adhered were then circled with a wax pen and 150µL of paraformaldehyde or formalin was added for 15 minutes. The fixing agent was tapped off and then 150µL of PBS with 0.1% Triton-X was added for 15 minutes. The slides were stained using hematoxylin for 1 minute. The slides then proceed from 70% ethanol for 1 minute, 95% ethanol for 1 minute, 100% ethanol for 3 minutes, xylene for 3 minutes, and finally xylene for 3 minutes. The slides were then allowed to dry and the cover slip was added using cytoseal (Thermo #527665). Slides were imaged at 60x and 500 nuclei from each mouse were called for differentiation state in a blinded fashion.

### Western Blot

Cells were lysed with RIPA buffer (50 mM Tris, 150 mM NaCl, 0.5% Deoxycholate, 1% NP-40, 0.1% SDS, 1 mM DTT, and 1% protease/phosphatase inhibitor) and supernatant was cleared by centrifugation. Protein concentration was determined using the Pierce BCA assay kit (Thermo, 23227). Between 40mg and 120mg of each protein sample was boiled in Laemmli buffer with 10% β-Mercaptoethanol and equal percentages of each sample were run on 4-15% polyacrylamide gels (BioRad, 4561086). Resolved proteins were wet-transferred to nitrocellulose membranes (Amersham), which were then blocked in a 5% BSA (VWR, 9048-46-8) solution made in 1x TBST buffer (20 mM Tris base, 0.15 M NaCl, 0.1% Tween, adjusted to pH 7.6). Membranes were then incubated with antibody solutions prepared in 5% BSA overnight. Antibodies used were (H3K27me3 Cell Signaling C26B11 1:500, EZH2 Cell Signaling 5246S 1:200, B2M Millipore MABF1968 1:2000, HLA-DR,DQ,DP AbCAM ab7856 1:500, Total Histone H3 AbCAM ab1791 1:5000). After washing, secondary antibodies were added (Novus anti-Rabbit-HRP and anti-Mouse-HRP), incubated, and washed. Bands were visualized with West Plus Pico ECL (Thermo) and exposed to Hyperfilm™ ECL™ film (Amersham). Protein molecular weights were determined with Precision Plus Protein™ Kaleidoscope™ Prestained Protein Standards (BioRad).

### MRI of genetically engineered mouse models

After adeno-Cre instillation, tumors were monitored via magnetic resonance imaging (MRI) and when tumor burden was measurable, mice were placed on one of four arms of a treatment regimen. Animals were anaesthetized via inhalation of isoflurane and kept warm on heated waterbed, vitals were monitored via cardiac and respiratory cycle (SA instruments), and recorded every 10 minutes while the animal was under anesthesia. The SA instruments pneumatic respiratory monitor was used to remove breathing artifacts by gating on the respiratory cycle. The Bruker ClinScan system used to scan the animals had 12 cm of actively shielded gradients, maximum strength 630 mT/m, and a slew rate of 6300 T/m/s. This instrument is a 7T system with 2×2 array coil and 2D gradient echoT1-weighted sequences. The parameters used for imaging are as follows: 18 slices, TR = 170 ms, TE = 2.4 ms, α=38°, Navg=3, FOV 26 x 26 mm^2^, 1mm thickness, matrix size 256 x 256, for a voxel size of 0.102 x 0.102 x 1.0 mm. In 2021, the system was upgraded to a Bruker Biospec system. For this upgraded machine, the Bruker IntraGate software was used to remove respiratory and cardiac motion with parameters: 18 slices, TR = 8.96ms ms, TE = 3.4 ms, α=10°, oversampling = 28, FOV 26 x 26 mm2, 1mm thickness, matrix size 192 x 192, 10 minutes. Models were then built on Slicer 3D software to calculate tumor volume.

### Tumor Cell 3D culture

Murine tumoroids were seeded in DMEM/F12 media (ThermoFisher, 11330032) with an additional 4mM Glutamax (ThermoFisher, 35050061), 5µg/mL ITS (insulin/transferrin/selenium, SIGMA, I3146) and 8-9% fetal bovine serum (VWR, 97068-085), 12.5 mg/ml bovine pituitary extract (Invitrogen, 13028-014), 0.1 mg/ml cholera toxin (SIGMA, C-8052), 25 ng/ml mEGF (Invitrogen, 53003018) and 25 ng/ml rmFGF2 (R&D Systems, 3139-FB/CF). Tumoroids were seeded and maintained in growth factor reduced and phenol red-free Matrigel (Corning, 47743-722) in transwells with 0.4 µm pore size (Corning). Human tumoroids were seeded in DMEM/F12 media (ThermoFisher, 11330032) with an additional 4mM Glutamax (ThermoFisher, 35050061), 20ng/mL FGF7 (VWR 10771-958), 50ng/mL FGF10 (VWR 10772-106), 40ng/mL Noggin (VWR 10772-456), 500nM A83-01 (R&D systems 2939), 5uM Y-27632 (Abmole Y-27632), 500nM SB202190 (SIGMA S7067), B27 Supplement (Gibco 1750-44), 1.25mM N-acetylcysteine (SIGMA A9165), 5mM Nicotinamide (SIGMA N0636), and penicillin/streptomycin (Invtirogen 15140-122)^42^. Tumoroid cultures were established in the presence of plasmocin (Invivogen Ant-mpt-1). Tumoroids were seeded and allowed to become established before starting treatment. Tumoroids were placed on six different treatment arms: DMSO as vehicle control, GSK126 (5μM), EPZ6438 (5μM), IFN-γ (20ng/mL), or combination of GSK126 (5μM) and IFN-γ (20ng/mL) or EPZ6438 (5μM) and IFN-γ (20ng/mL). Tumoroids were fed every two days for 11 days total, adding in IFN-γ on day nine.

### ChIPseq

To perform ChIP analysis, we followed the ChIP Cell Fixation protocol provided by Active Motif. Cells were fixed by adding 1/10^th^ volume of freshly prepared solution of 11% Formaldehyde (SIGMA #F-8775), 0.1M NaCl (Fisher Scientific #S271-10), 1mM EDTA pH 8.0 (Invitrogen #AM9261), 50mM HEPES pH 7.9 (SIGMA #H0887), diluted in H_2_O to the dissociated cell in media, and incubated for 15 minutes at room temperature. The samples were then quenched by added a 1/20^th^ volume of 2.5M Glycine (SIGMA #G-7403). Next, the samples were washed with cold 0.5% IGEPAL® CA-630 (SIGMA #I8896) in PBS pH 7.4 and centrifuged at 800g for 10 minutes at 4°C. The supernatant was removed and the cell pellets were washed in cold 0.5% IGEPAL®-PBS and centrifuged for another 10 minutes at 4°C. Cells were then resuspended with 100mM PMSF in ethanol, and centrifuged once again. Supernatant was removed and pellets were snap frozen with liquid nitrogen and stored at -80°C. These samples were sent to Active Motif for ChIP-sequencing using the antibodies H3K27me3 (Active Motif 39155, 4µL antibody per 40µg chromatin), H3K4me3 (Active Motif 39159, 4µL antibody per 40µg chromatin) or H3K27ac (Active Motif 39133 4µL antibody per 40µg chromatin). Quality control and read alignment was performed by ActiveMotif. Briefly, the 75-bp single-end sequence reads were mapped to the human reference genome hg38 using the bwa samse with default settings. Reads that had >2 mismatches and multimapping reads were removed followed by PCR deduplication. The resulting bam files were normalized to account for the differences in the sequencing depth. Samples within each antibody group were reduced by random sampling to the number of unique alignments present in the smallest sample. Since the 5’-ends of the aligned reads represent the ends of the ChIP/IP-fragments, the tags were extended in silico using Active Motif software at their 3’-ends to a length of 200bp. To identify the density of fragments along the genome, the genome was divided into 32-nt bins and the number of fragments in each bin was determined. The MACS2 version v2.1.0 peak finding algorithm was used to identify regions of ChIP-seq enrichment over background, with pvalue threshold of enrichment 1E-07 for all datasets. Genomic regions known to have low sequencing confidence were removed using blacklisted regions defined by the ENCODE project. The selected peak intervals were annotated to the nearest transcription start sites (TSS) using the KnownGene hg38 TSS annotation. To compare peak metrics, overlapping intervals were grouped into merged regions, defined by the start coordinate of the most upstream interval and the end coordinate of the most downstream interval. In locations where only one sample has an interval, this interval defines the merged region. Peak distribution patterns were obtained using seqplots across all merged intervals from −5 kb to +5 kb to include distal promoters and regulatory regions. Heatmaps were generated for visualization of tag distributions, which are mapped across target regions. The average values for all target regions in heatmaps were calculated and plotted in histograms. Peaks unique to each genotype or conserved in multiple genotypes were annotated by GREAT^43^ to associate each genomic region with all genes whose regulatory domain it overlaps. The resulting gene lists were used to identify significantly enriched gene signatures from GSEA curated signature gene sets.

### Histology and HALO**®**

Mice were euthanized and lungs were inflated with 10% buffered formalin overnight, then stored in 70% ethanol. Tissues were embedded in paraffin and sectioned at the Markey Cancer Center Biospecimens Procurement and Translational Pathology Shared Resource Facility (BPTP SRF). H&E-stained slides were scanned at 40x with an Aperio slide scanner and images were used for further analysis. To validate that the sub-cutaneous grafts contained similar cellular proportions to lung tumors, we used our previously described nuclear phenotyper^19, 39^ shown in **Figure 5C**.

### Single cell RNA sequencing

Lungs were harvested from mice and dissociated by finely mincing with scissors and triturating with a 5mL serological pipette filled with Ca2+/Mg2+-free PBS. In order to enrich for immune cell populations, they were then stained with EpCAM-PECy7 (BioLegend #118216) and CD31-APC (Biolegend Cat# 102510), bound to beads and run through Miltenyi LS columns (#130-042-401) on a magnet. The flow through was collected and cells were captured for reverse transcription by a 10x Genomics Chromium controller. Bone marrow cells were isolated on a histopaque gradient as described above. Reverse transcription was performed using the 10X Genomics Single Cell 3’ v3 Kit. Libraries were prepared and sequenced, and the sequencer-produced Chromium single-cell data and then the Cell Ranger toolkit version 3.1 (10X Genomics) was used to de-multiplex samples from raw sequencing reads to gene-count matrices with alignment to the mm10 genome (v93). In order to perform the downstream analysis, such as cell type identification and differential gene expression analysis, Seurat (V3) R package^44^ was employed to aggregate the gene-counts matrices from all samples and provide the analytical insight. A five-step process was performed using Seurat package: (1) For quality control and data pre-processing, we discarded genes expressed in fewer than three cells and discarded the low-quality cells that had less than 100 genes expressed or percentage of mitochondrial transcripts > 7. The count matrices were then log-transformed. (2) For sample merging and feature selection, we combined the log-transformed matrices of each sample and applied the variance-stabilizing transformation (vst) method to remove cell-to-cell variation. The top 1500 genes were selected for sample integration. (3) For dimension reduction and clustering, we applied “PCA” to reduce the dimensionality of the merged data to 50 principal components, and then performed a shared nearest neighbor (SNN) modularity optimization-based algorithm to identify clusters of cells. We utilized the UMAP ^45^ technique to visualize the clustering results as shown in **Figure 6A**. (4) Cell cluster identities were called by examining highly expressed genes in each cell cluster (**Supp Table 3**). (5) For differential gene expression analysis for each cell population, we used a DEG (Differential Expressed Gene) identification method in Seurat, namely “MAST”^46^, to identify the up/down-regulated sets of genes between treatment groups in the three major populations, neutrophils, macrophage/dendritic cells, and tumor cells. In the **Figure 6B**, the distribution of cell clusters across genotypes was assessed by z-test and p values were adjusted for multiple hypothesis testing.

### RNA Isolation

To perform RNA isolation Absolutely RNA Miniprep kits were used (Agilent #400805). Cell pellets were re-suspended in lysis buffer plus 0.7% β-mercaptoethanol and stored at -80°C. An equal volume of 70% nuclease free ethanol was added to each sample and the solution was added to a column. Columns were washed once with low-salt wash buffer once and DNase digestion was performed for 15 minutes at 37°C. The column was then washed once with high-salt buffer, followed by two more low-salt washes. The RNA was eluted with pre-warmed elution buffer for 2 minutes and stored at -80°C.

### RT-qPCR and Sequencing

Concentration of RNA samples was quantified by NanoDrop 8000™ spectrophotometer (Thermo Fisher #ND-8000-GL). To perform cDNA synthesis, random hexamers (50ng/μL) and dNTPs (10mM) were mixed in a 1:1 ratio and 2μL was placed in each PCR tube. Next, 1000ng of RNA was added and volume was brought up to 12μL, placed in C1000 Touch™ thermal cycler (BioRad), and run for 5 minutes at 65°C. A master mix of reagents was made for each sample (5x reverse transcription buffer, 50mM MgCl_2_ (Invitrogen #AM9530), 0.1M dithiothreitol, 40U/μL RNaseOUT™ (Invitrogen #100000840), 200U/μL SuperScript™ III (Invitrogen #56575). The cDNA protocol was performed on thermal cycler as follows: 25°C for 10 minutes, 50°C for 50 minutes, 70°C for 15 minutes. After the 70°C step, 1μL of RNase H (Ambion #AM2293) was added to each tube and protocol continued for 20 minutes at 37°C. After genertion, cDNA is diluted 1:5 before performing qPCR and stored at -80°C. To perform qPCR master mixes were made for each gene of interest using TaqMan™ Fast Advanced Master Mix (Invitrogen #4444964) and TaqMan™ qPCR Assays, then run on QuantStudio 3 (applied biosystems #A28567). Data were analyzed by calculating *Gene of Interest*(Ct_reference_-Ct_exprimental_)-*Gapdh*(Ct_reference_-Ct_experimental_) and the data were graphed on the log_2_ scale. Library preparation and sequencing were performed by the Beijing Genomics Institute (BGI Group) using DNBseq to a depth of 24 million 100bp paried-end reads. Sequencing reads were trimmed and filtered using Trimmomatic (V0.39)^47^ to remove adapters and low-quality reads. Reads from human samples were mapped to Ensembl GRCh38 transcript annotation (release 98), and mouse samples to Ensembl GRCm38 (mm10) transcript annotation (release 82), using RSEM^48^. Gene expression data normalization and differential expression analysis were performed using the R package edgeR^49^.

### Statistics and reproducibility

Statistical analyses were carried out using GraphPad Prism or Microsoft Excel. Unless otherwise stated, all numerical data are presented as mean ± standard error of the mean. For grouped analyses, one-way ANOVA with Holm-Šídák’s multiple comparisons correction was used. A p value (or adjusted p value) less than 0.05 was considered statistically significant.

## ACKNOWLEDGEMENTS

The authors thank Dave Powell in the Markey Cancer Center Small Animal Imaging Facility (MCCSAIF) for extensive help with the MRI scanning, and Drs. Doug Harrison and Jim Begley at the University of Kentucky A&S Imaging Center for preparation of the scRNAseq samples and initial analysis. This work was supported in part by NCI K22 CA201036, Kentucky Lung Cancer Research Program, V Foundation Scholar Award, American Cancer Society Grants IRG-85-001-25 and 133123-RSG-19-081-01-TBG, NCI R01 CA237643, the American Institute for Cancer Research, and American Association for Cancer Research-Bayer Innovation and Discovery Grant (CFB), NIGMS P20 GM121327-03 (CFB), NCI T32 CA165990 (DRP), NIEHS T32 5T32ES007266 (TJD and ALB), and NHLBI F31 HL151111(ALB). This research was also supported by the Biostatistics & Bioinformatics, Cancer Research Informatics, Oncogenomics, Biospecimen Procurement & Translational Pathology, and Flow Cytometry & Immune Monitoring Shared Resource Facilities of the University of Kentucky Markey Cancer Center (P30CA177558).

## DATA AVAILABILITY STATEMENT

All data presented in this manuscript are available from the corresponding author upon reasonable request. The sequencing data are available at NCBI GEO under the super-series GSE233665.

The RNA-sequencing data are available under accession number: GSE233468 The ChIP-sequencing data are available under accession number: GSE233469 The scRNA-sequencing data are available under accession number: GSE233665

## AUTHOR CONTRIBUTIONS

Conceptualization: CFB, TJD, Jinpeng L, KKW; Data curation: CFB, TJD, DRP, XQ, Jinpeng L; Formal analysis: CFB, TJD, XQ, Jinpeng L; Funding acquisition: CFB, KKW; Data acquisition and processing: CFB, TJD, KJN, FC, ARC, ARE, XQ, FL; Bioinformatics and biostatistics: XQ, Jinze L, Jinpeng L; Writing-original draft: TJD, CFB; Writing - review & editing: TJD, DRP, CFB.

