## Supplemental Tables and Figures for "EZH2 inhibition promotes tumor immunogenicity in lung squamous cell carcinomas"

Tanner J. DuCote et al.,

**Supplementary Tables 1-6**

**Supplementary Figures 1-6**

**Supplemental Table 1: GSEA on RNAseq of Tumoroids, related to Figure 3**
**NES=Normalized Enrichment Score, FDR=False Discovery Rate**

| MSigDB Signature Name | Patient 1 | | Patient 2 | | Patient 1 | | Patient 2 | |
|---|---|---|---|---|---|---|---|---|
| | *GSK126 vs Veh* | | *GSK126 vs Veh* | | *EPZ6438 vs Veh* | | *EPZ6438 vs Veh* | |
| | NES | FDR q-val | NES | FDR q-val | NES | FDR q-val | NES | FDR q-val |
| HALLMARK_MYC_TARGETS_V1 | -2.274 | 0.000 | -2.505 | 0.000 | | 1.000 | -2.127 | 0.000 |
| HALLMARK_MYC_TARGETS_V2 | -1.482 | 0.079 | -1.940 | 0.014 | -1.711 | 0.017 | -1.743 | 0.026 |
| HALLMARK_DNA_REPAIR | -1.356 | 0.085 | -1.413 | 0.097 | -0.833 | 0.644 | -1.207 | 0.217 |
| HALLMARK_UNFOLDED_PROTEIN_RESPONSE | -1.360 | 0.096 | -1.273 | 0.160 | -1.542 | 0.039 | -1.142 | 0.268 |
| HALLMARK_E2F_TARGETS | -0.922 | 0.688 | -1.357 | 0.117 | | 1.000 | -0.632 | 0.966 |
| HALLMARK_REACTIVE_OXYGEN_SPECIES_PATHWAY | -1.868 | 0.008 | -1.594 | 0.058 | -1.818 | 0.007 | -1.625 | 0.035 |
| HALLMARK_OXIDATIVE_PHOSPHORYLATION | -2.657 | 0.000 | -2.152 | 0.000 | -2.161 | 0.004 | -2.125 | 0.000 |
| HALLMARK_INFLAMMATORY_RESPONSE | 1.501 | 0.025 | 0.821 | 0.927 | 1.242 | 0.183 | -1.659 | 0.036 |
| HALLMARK_INTERFERON_GAMMA_RESPONSE | 1.273 | 0.164 | 0.999 | 0.793 | 1.479 | 0.036 | 0.639 | 1.000 |
| | *GSK+IFN vs IFN* | | *GSK+IFN vs IFN* | | *EPZ+IFN vs IFN* | | *EPZ+IFN vs IFN* | |
| | NES | FDR q-val | NES | FDR q-val | NES | FDR q-val | NES | FDR q-val |
| HALLMARK_MYC_TARGETS_V1 | -1.753 | 0.024 | -2.284 | 0.000 | -1.801 | 0.026 | -2.111 | 0.000 |
| HALLMARK_MYC_TARGETS_V2 | -1.527 | 0.062 | -2.244 | 0.000 | -1.637 | 0.051 | -2.139 | 0.000 |
| HALLMARK_DNA_REPAIR | -1.064 | 0.420 | -1.540 | 0.069 | -1.121 | 0.361 | -1.419 | 0.117 |
| HALLMARK_UNFOLDED_PROTEIN_RESPONSE | 0.892 | 0.977 | -0.829 | 0.814 | 0.716 | 1.000 | -0.648 | 0.965 |
| HALLMARK_E2F_TARGETS | -1.001 | 0.478 | -1.530 | 0.054 | -0.480 | 1.000 | -0.730 | 0.984 |
| HALLMARK_REACTIVE_OXYGEN_SPECIES_PATHWAY | -1.763 | 0.032 | -1.321 | 0.149 | -1.806 | 0.038 | -1.398 | 0.096 |
| HALLMARK_OXIDATIVE_PHOSPHORYLATION | -2.091 | 0.002 | -1.393 | 0.107 | -2.064 | 0.000 | -1.557 | 0.067 |
| HALLMARK_INFLAMMATORY_RESPONSE | 1.503 | 0.036 | 0.882 | 0.934 | 1.178 | 0.421 | -1.413 | 0.100 |
| HALLMARK_INTERFERON_GAMMA_RESPONSE | 1.722 | 0.002 | 1.066 | 0.698 | 1.385 | 0.118 | 0.912 | 1.000 |
| | **Mouse 1** | | **Mouse 2** | | **Mouse 1** | | **Mouse 2** | |
| | *GSK126 vs Veh* | | *GSK126 vs Veh* | | *EPZ6438 vs Veh* | | *EPZ6438 vs Veh* | |
| | NES | FDR q-val | NES | FDR q-val | NES | FDR q-val | NES | FDR q-val |
| HALLMARK_MYC_TARGETS_V1 | -1.948 | 0.001 | -1.404 | 0.086 | -1.701 | 0.089 | -1.465 | 0.092 |
| HALLMARK_MYC_TARGETS_V2 | -1.635 | 0.017 | -1.463 | 0.064 | -1.543 | 0.110 | -0.778 | 1.000 |
| HALLMARK_DNA_REPAIR | -1.572 | 0.025 | -0.815 | 0.981 | -1.542 | 0.073 | -1.315 | 0.167 |
| HALLMARK_UNFOLDED_PROTEIN_RESPONSE | -1.348 | 0.112 | -1.000 | 0.690 | -1.313 | 0.178 | -0.693 | 0.931 |
| HALLMARK_E2F_TARGETS | -2.393 | 0.000 | -2.560 | 0.000 | -0.597 | 0.980 | -2.231 | 0.003 |
| HALLMARK_REACTIVE_OXYGEN_SPECIES_PATHWAY | -1.054 | 0.424 | 0.979 | 0.591 | -1.317 | 0.218 | 0.695 | 0.960 |
| HALLMARK_OXIDATIVE_PHOSPHORYLATION | 0.534 | 0.998 | 0.499 | 0.999 | -1.198 | 0.258 | -0.728 | 1.000 |
| HALLMARK_INFLAMMATORY_RESPONSE | 2.178 | 0.000 | 1.233 | 0.227 | 1.811 | 0.000 | 1.662 | 0.006 |
| HALLMARK_INTERFERON_GAMMA_RESPONSE | 1.772 | 0.014 | 1.614 | 0.032 | 1.775 | 0.001 | 1.811 | 0.000 |
| | *GSK+IFN vs IFN* | | *GSK+IFN vs IFN* | | *EPZ+IFN vs IFN* | | *EPZ+IFN vs IFN* | |
| | NES | FDR q-val | NES | FDR q-val | NES | FDR q-val | NES | FDR q-val |
| HALLMARK_MYC_TARGETS_V1 | -1.618 | 0.050 | -2.248 | 0.000 | -2.003 | 0.008 | -2.443 | 0.000 |
| HALLMARK_MYC_TARGETS_V2 | -2.020 | 0.007 | -2.006 | 0.000 | -1.930 | 0.007 | -1.969 | 0.005 |
| HALLMARK_DNA_REPAIR | -0.938 | 0.889 | -1.046 | 0.469 | -1.350 | 0.144 | -1.591 | 0.030 |
| HALLMARK_UNFOLDED_PROTEIN_RESPONSE | -1.559 | 0.044 | -1.936 | 0.001 | -1.732 | 0.017 | -1.955 | 0.004 |
| HALLMARK_E2F_TARGETS | -1.185 | 0.458 | -2.775 | 0.000 | -1.024 | 0.392 | -2.832 | 0.000 |
| HALLMARK_REACTIVE_OXYGEN_SPECIES_PATHWAY | -0.911 | 0.884 | 0.768 | 0.945 | -1.309 | 0.143 | -0.828 | 0.799 |
| HALLMARK_OXIDATIVE_PHOSPHORYLATION | 0.510 | 0.998 | -0.779 | 1.000 | -1.114 | 0.294 | -1.681 | 0.023 |
| HALLMARK_INFLAMMATORY_RESPONSE | 2.196 | 0.000 | 1.512 | 0.108 | 1.894 | 0.000 | 1.895 | 0.000 |
| HALLMARK_INTERFERON_GAMMA_RESPONSE | 1.888 | 0.001 | 1.426 | 0.121 | 1.663 | 0.007 | 1.937 | 0.000 |

**Supplemental Table 2: GSEA on Genes That were Up-regulated and Lost H3K27me3 Peaks in Combination Treatment vs IFNy alone, related to Figure 4**

| Group | MSigDB Signature Name | GeneRatio | BgRatio | P Value | Adj. P value | FDR q value |
|---|---|---|---|---|---|---|
| Inflammatory Responses | MANNE_COVID19_COMBINED_COHORT_VS_HEALTHY_DONOR_PLATELETS_DN | 16/272 | 228/21697 | 3.32E-08 | 4.54878E-06 | 3.8288E-06 |
| | MANNE_COVID19_ICU_VS_HEALTHY_DONOR_PLATELETS_DN | 13/272 | 162/21697 | 1.4E-07 | 1.74026E-05 | 1.46481E-05 |
| | REACTOME_INTERFERON_GAMMA_SIGNALING | 8/272 | 93/21697 | 2.23E-05 | 0.001175233 | 0.000989219 |
| | REACTOME_PD_1_SIGNALING | 5/272 | 28/21697 | 2.32E-05 | 0.001175233 | 0.000989219 |
| | REACTOME_MHC_CLASS_II_ANTIGEN_PRESENTATION | 9/272 | 126/21697 | 3.07E-05 | 0.00150398 | 0.001265932 |
| | KEGG_ASTHMA | 5/272 | 30/21697 | 3.29E-05 | 0.001564249 | 0.001316661 |
| | KEGG_ALLOGRAFT_REJECTION | 5/272 | 37/21697 | 9.37E-05 | 0.003565296 | 0.003000985 |
| | KEGG_GRAFT_VERSUS_HOST_DISEASE | 5/272 | 41/21697 | 0.000155 | 0.005362662 | 0.004513866 |
| Cell Adhesion and Signaling | KEGG_CELL_ADHESION_MOLECULES_CAMS | 12/272 | 133/21697 | 1.18E-07 | 1.54175E-05 | 1.29773E-05 |
| | ZWANG_TRANSIENTLY_UP_BY_2ND_EGF_PULSE_ONLY | 50/272 | 1941/21697 | 7.24E-07 | 6.61582E-05 | 5.56867E-05 |
| | ONDER_CDH1_TARGETS_2_UP | 12/272 | 257/21697 | 0.000104 | 0.003918949 | 0.003298662 |
| | KEGG_FOCAL_ADHESION | 11/272 | 199/21697 | 4.51E-05 | 0.001992359 | 0.001677011 |
| TP53 and Apoptosis | PEREZ_TP53_TARGETS | 40/272 | 1198/21697 | 1.49E-08 | 2.39558E-06 | 2.01641E-06 |
| | BRUINS_UVC_RESPONSE_VIA_TP53_GROUP_A | 30/272 | 884/21697 | 7.99E-07 | 6.84384E-05 | 5.7606E-05 |
| | PEREZ_TP53_AND_TP63_TARGETS | 11/272 | 208/21697 | 6.73E-05 | 0.002792285 | 0.002350325 |
| | HANN_RESISTANCE_TO_BCL2_INHIBITOR_UP | 5/272 | 36/21697 | 8.19E-05 | 0.003298434 | 0.002776362 |
| Nervous System | REACTOME_NEURONAL_SYSTEM | 27/272 | 410/21697 | 2.36E-12 | 6.46166E-10 | 5.43892E-10 |
| | REACTOME_TRANSMISSION_ACROSS_CHEMICAL_SYNAPSES | 16/272 | 269/21697 | 3.22E-07 | 3.39241E-05 | 2.85546E-05 |
| | REACTOME_NERVOUS_SYSTEM_DEVELOPMENT | 23/272 | 580/21697 | 1.28E-06 | 0.000103351 | 8.69924E-05 |
| | REACTOME_NEUROTRANSMITTER_RECEPTORS_AND_POSTSYNAPTIC_SIGNAL_ | 10/272 | 205/21697 | 0.000276 | 0.008316188 | 0.006999911 |
| Polycomb Targets | BENPORATH_ES_WITH_H3K27ME3 | 76/272 | 1114/21697 | 3.02E-35 | 8.26451E-32 | 6.95641E-32 |
| | BENPORATH_SUZ12_TARGETS | 70/272 | 1033/21697 | 4.01E-32 | 5.49216E-29 | 4.62287E-29 |
| | MIKKELSEN_MEF_HCP_WITH_H3K27ME3 | 51/272 | 590/21697 | 4.28E-28 | 3.9059E-25 | 3.28768E-25 |
| | BENPORATH_EED_TARGETS | 64/272 | 1058/21697 | 1.75E-26 | 1.19586E-23 | 1.00658E-23 |
| | MEISSNER_BRAIN_HCP_WITH_H3K4ME3_AND_H3K27ME3 | 64/272 | 1073/21697 | 3.81E-26 | 2.08627E-23 | 1.75606E-23 |
| | MIKKELSEN_MCV6_HCP_WITH_H3K27ME3 | 43/272 | 437/21697 | 5.7E-26 | 2.60336E-23 | 2.1913E-23 |
| | BENPORATH_PRC2_TARGETS | 49/272 | 649/21697 | 2.32E-24 | 9.07209E-22 | 7.63617E-22 |
| | MIKKELSEN_NPC_HCP_WITH_H3K27ME3 | 26/272 | 345/21697 | 2.83E-13 | 9.6866E-11 | 8.15342E-11 |
| | MEISSNER_NPC_HCP_WITH_H3K4ME2_AND_H3K27ME3 | 26/272 | 350/21697 | 3.95E-13 | 1.20105E-10 | 1.01095E-10 |
| | MEISSNER_BRAIN_HCP_WITH_H3K27ME3 | 20/272 | 271/21697 | 2.52E-10 | 6.27713E-08 | 5.28359E-08 |
| | MIKKELSEN_NPC_HCP_WITH_H3K4ME3_AND_H3K27ME3 | 17/272 | 210/21697 | 1.42E-09 | 3.23121E-07 | 2.71978E-07 |
| | MEISSNER_NPC_HCP_WITH_H3K4ME3_AND_H3K27ME3 | 13/272 | 143/21697 | 3.19E-08 | 4.54878E-06 | 3.8288E-06 |

**Supplemental Table 3: Genes Highly Expressed in All 16 Clusters, Related to Figure 6**
**Log2FC=Log2-fold change between cluster and all others, pct=percentage of cells expressing**

| Cluster | Gene Symbol | Log2FC | pct.1 | pct.2 | P value | Adj P value | Cluster | Gene Symbol | Log2FC | pct.1 | pct.2 | P value | Adj P value |
|---|---|---|---|---|---|---|---|---|---|---|---|---|---|
| Tumor | Krt19 | 4.685 | 0.977 | 0.814 | 0.0E+00 | 0.0E+00 | Dendritic | Cd74 | 4.390 | 0.964 | 0.555 | 0.0E+00 | 0.0E+00 |
| | Ppbp | 4.050 | 0.781 | 0.673 | 0.0E+00 | 0.0E+00 | | H2-Ab1 | 4.061 | 0.934 | 0.629 | 0.0E+00 | 0.0E+00 |
| | S100a7a | 3.910 | 0.908 | 0.872 | 0.0E+00 | 0.0E+00 | | C1qa | 3.857 | 0.988 | 0.581 | 0.0E+00 | 0.0E+00 |
| | S100a10 | 3.589 | 0.968 | 0.700 | 0.0E+00 | 0.0E+00 | | H2-Aa | 3.618 | 0.918 | 0.556 | 0.0E+00 | 0.0E+00 |
| | Ly6d | 3.380 | 0.853 | 0.767 | 0.0E+00 | 0.0E+00 | | C1qb | 3.455 | 0.982 | 0.503 | 0.0E+00 | 0.0E+00 |
| | Prss27 | 3.341 | 0.899 | 0.825 | 0.0E+00 | 0.0E+00 | | Apoe | 3.074 | 0.985 | 0.781 | 0.0E+00 | 0.0E+00 |
| | Krt18 | 3.333 | 0.824 | 0.698 | 0.0E+00 | 0.0E+00 | | C1qc | 2.985 | 0.988 | 0.640 | 0.0E+00 | 0.0E+00 |
| | Plet1 | 3.301 | 0.954 | 0.684 | 0.0E+00 | 0.0E+00 | | Ms4a6c | 2.243 | 0.930 | 0.407 | 0.0E+00 | 0.0E+00 |
| | Nupr1 | 3.285 | 0.892 | 0.671 | 0.0E+00 | 0.0E+00 | | Ms4a7 | 2.238 | 0.969 | 0.433 | 0.0E+00 | 0.0E+00 |
| | S100a16 | 3.221 | 0.951 | 0.730 | 0.0E+00 | 0.0E+00 | | Ctss | 1.932 | 0.990 | 0.654 | 0.0E+00 | 0.0E+00 |
| | Krt23 | 3.208 | 0.894 | 0.638 | 0.0E+00 | 0.0E+00 | | Cst3 | 1.844 | 0.987 | 0.817 | 0.0E+00 | 0.0E+00 |
| | Ly6c1 | 2.943 | 0.832 | 0.652 | 0.0E+00 | 0.0E+00 | | Gatm | 1.752 | 0.937 | 0.517 | 0.0E+00 | 0.0E+00 |
| | Atp1b1 | 2.870 | 0.888 | 0.682 | 0.0E+00 | 0.0E+00 | | Lair1 | 1.530 | 0.895 | 0.546 | 0.0E+00 | 0.0E+00 |
| | Irf7 | 2.800 | 0.937 | 0.588 | 0.0E+00 | 0.0E+00 | | Pld4 | 1.525 | 0.899 | 0.187 | 0.0E+00 | 0.0E+00 |
| | Isg15 | 2.772 | 0.869 | 0.644 | 0.0E+00 | 0.0E+00 | | Tmem119 | 0.828 | 0.959 | 0.256 | 0.0E+00 | 0.0E+00 |
| | Tspan1 | 2.562 | 0.857 | 0.516 | 0.0E+00 | 0.0E+00 | | Cxcl9 | 0.789 | 0.946 | 0.644 | 0.0E+00 | 0.0E+00 |
| | Hspa5 | 2.539 | 0.951 | 0.818 | 0.0E+00 | 0.0E+00 | | Ccl8 | 0.651 | 0.962 | 0.382 | 0.0E+00 | 0.0E+00 |
| | Glrx | 2.448 | 0.869 | 0.594 | 0.0E+00 | 0.0E+00 | | Retnla | 0.821 | 0.957 | 0.743 | 2.9E-289 | 4.4E-286 |
| | Emp1 | 2.302 | 0.920 | 0.778 | 0.0E+00 | 0.0E+00 | | Cybb | 1.677 | 0.924 | 0.514 | 1.9E-288 | 2.9E-285 |
| | Hopx | 2.294 | 0.766 | 0.477 | 0.0E+00 | 0.0E+00 | | Gngt2 | 1.535 | 0.964 | 0.637 | 4.8E-284 | 7.2E-281 |
| | Cd81 | 2.276 | 0.909 | 0.547 | 0.0E+00 | 0.0E+00 | | Cxcl16 | 1.511 | 0.983 | 0.701 | 5.4E-279 | 8.1E-276 |
| | Arg1 | 2.255 | 0.872 | 0.739 | 0.0E+00 | 0.0E+00 | | Fyb | 1.709 | 0.949 | 0.688 | 1.7E-277 | 2.5E-274 |
| | Scgb1a1 | 2.246 | 0.831 | 0.809 | 0.0E+00 | 0.0E+00 | | Pou2f2 | 1.956 | 0.849 | 0.421 | 1.4E-270 | 2.1E-267 |
| | Lmo7 | 2.226 | 0.849 | 0.750 | 0.0E+00 | 0.0E+00 | | Acp5 | 1.404 | 0.968 | 0.607 | 4.2E-268 | 6.3E-265 |
| | Ceacam1 | 2.214 | 0.888 | 0.649 | 0.0E+00 | 0.0E+00 | | Lgmn | 1.372 | 0.964 | 0.624 | 2.1E-257 | 3.2E-254 |
| | Dusp14 | 2.072 | 0.836 | 0.675 | 0.0E+00 | 0.0E+00 | | Aif1 | 2.087 | 0.853 | 0.573 | 2.8E-255 | 4.3E-252 |
| | Wnt4 | 2.037 | 0.882 | 0.722 | 0.0E+00 | 0.0E+00 | | Atp1a1 | 1.325 | 0.957 | 0.700 | 3.7E-254 | 5.6E-251 |
| | Clu | 2.031 | 0.918 | 0.761 | 0.0E+00 | 0.0E+00 | | Lyz2 | 0.748 | 0.994 | 0.755 | 9.3E-252 | 1.4E-248 |
| | Epcam | 2.006 | 0.848 | 0.601 | 0.0E+00 | 0.0E+00 | | Trf | 1.377 | 0.891 | 0.499 | 4.4E-247 | 6.7E-244 |
| | Anxa1 | 2.002 | 0.919 | 0.772 | 0.0E+00 | 0.0E+00 | | Ms4a6d | 1.230 | 0.895 | 0.733 | 2.9E-240 | 4.4E-237 |
| | Cldn3 | 1.963 | 0.803 | 0.773 | 0.0E+00 | 0.0E+00 | | Csf1r | 1.731 | 0.824 | 0.437 | 2.0E-233 | 3.1E-230 |
| | 2310007B03Ril | 1.887 | 0.861 | 0.663 | 0.0E+00 | 0.0E+00 | | Bcl2a1d | 1.057 | 0.884 | 0.505 | 3.6E-223 | 5.4E-220 |
| | Pdzk1ip1 | 1.882 | 0.850 | 0.786 | 0.0E+00 | 0.0E+00 | | Fcgr1 | 0.866 | 0.839 | 0.480 | 3.3E-217 | 5.0E-214 |
| | Prss22 | 1.850 | 0.702 | 0.557 | 0.0E+00 | 0.0E+00 | | Napsa | 1.159 | 0.904 | 0.579 | 1.7E-213 | 2.5E-210 |
| | Aqp5 | 1.785 | 0.676 | 0.741 | 0.0E+00 | 0.0E+00 | | Ly86 | 1.085 | 0.807 | 0.287 | 7.3E-213 | 1.1E-209 |
| | Cldn7 | 1.777 | 0.732 | 0.679 | 0.0E+00 | 0.0E+00 | | Ctsc | 1.394 | 0.881 | 0.644 | 2.9E-207 | 4.4E-204 |
| | Jdp2 | 1.736 | 0.810 | 0.711 | 0.0E+00 | 0.0E+00 | | Gm2a | 1.234 | 0.909 | 0.624 | 5.8E-207 | 8.7E-204 |
| | Plpp1 | 1.699 | 0.783 | 0.653 | 0.0E+00 | 0.0E+00 | | Mafb | 1.508 | 0.853 | 0.588 | 4.2E-199 | 6.3E-196 |
| | Avpi1 | 1.689 | 0.843 | 0.712 | 0.0E+00 | 0.0E+00 | | Ccr2 | 0.905 | 0.788 | 0.255 | 3.8E-195 | 5.7E-192 |
| | Nccrp1 | 1.658 | 0.754 | 0.730 | 0.0E+00 | 0.0E+00 | | Ifitm3 | 1.495 | 0.943 | 0.695 | 1.8E-194 | 2.6E-191 |
| | Errfi1 | 1.618 | 0.894 | 0.564 | 0.0E+00 | 0.0E+00 | | Pgls | 0.758 | 0.921 | 0.629 | 5.1E-182 | 7.6E-179 |
| | Plat | 1.601 | 0.627 | 0.504 | 0.0E+00 | 0.0E+00 | | Itgb5 | 0.923 | 0.849 | 0.676 | 1.1E-177 | 1.6E-174 |
| | Gsta4 | 1.599 | 0.796 | 0.689 | 0.0E+00 | 0.0E+00 | | Rpl3 | 1.055 | 0.970 | 0.708 | 1.1E-177 | 1.7E-174 |
| | Tacstd2 | 1.567 | 0.736 | 0.538 | 0.0E+00 | 0.0E+00 | | Gpx1 | 0.883 | 0.965 | 0.833 | 4.2E-173 | 6.3E-170 |
| | Calr | 1.558 | 0.945 | 0.734 | 0.0E+00 | 0.0E+00 | | Clec4a3 | 1.528 | 0.770 | 0.440 | 5.1E-172 | 7.6E-169 |
| | Tubb4b | 1.556 | 0.918 | 0.674 | 0.0E+00 | 0.0E+00 | | Cd72 | 1.163 | 0.717 | 0.163 | 1.4E-171 | 2.1E-168 |
| | Wwtr1 | 1.537 | 0.890 | 0.574 | 0.0E+00 | 0.0E+00 | | Cx3cr1 | 1.589 | 0.759 | 0.360 | 7.3E-171 | 1.1E-167 |
| | Fxyd3 | 1.529 | 0.782 | 0.582 | 0.0E+00 | 0.0E+00 | | Tgfbi | 0.964 | 0.915 | 0.601 | 2.9E-170 | 4.4E-167 |
| | Csrp1 | 1.520 | 0.819 | 0.671 | 0.0E+00 | 0.0E+00 | | Asah1 | 0.779 | 0.879 | 0.564 | 1.0E-169 | 1.5E-166 |
| | Spns3 | 1.512 | 0.823 | 0.552 | 0.0E+00 | 0.0E+00 | | Ptma | 0.975 | 0.965 | 0.726 | 1.6E-165 | 2.4E-162 |
| | Mall | 1.506 | 0.859 | 0.721 | 0.0E+00 | 0.0E+00 | | Smpdl3a | 1.034 | 0.904 | 0.622 | 3.0E-163 | 4.5E-160 |
| | Gprc5a | 1.498 | 0.799 | 0.586 | 0.0E+00 | 0.0E+00 | | Ucp2 | 0.852 | 0.973 | 0.726 | 5.8E-162 | 8.7E-159 |
| | Herpud1 | 1.429 | 0.790 | 0.560 | 0.0E+00 | 0.0E+00 | | Bcl2a1a | 0.700 | 0.859 | 0.545 | 1.3E-159 | 1.9E-156 |
| | Ddit3 | 1.405 | 0.820 | 0.723 | 0.0E+00 | 0.0E+00 | | Cd300e | 1.156 | 0.727 | 0.244 | 4.8E-158 | 7.1E-155 |
| | Clec2g | 1.386 | 0.818 | 0.436 | 0.0E+00 | 0.0E+00 | | Ctsz | 0.849 | 0.994 | 0.790 | 2.0E-152 | 3.1E-149 |
| | Dnajb1 | 1.374 | 0.893 | 0.567 | 0.0E+00 | 0.0E+00 | | Laptm5 | 0.768 | 0.989 | 0.732 | 2.3E-151 | 3.5E-148 |
| | Cidea | 1.304 | 0.610 | 0.536 | 0.0E+00 | 0.0E+00 | | Rassf4 | 0.995 | 0.789 | 0.576 | 1.5E-149 | 2.3E-146 |
| | Lrrc26 | 1.302 | 0.812 | 0.724 | 0.0E+00 | 0.0E+00 | | Emp3 | 1.135 | 0.840 | 0.456 | 8.3E-148 | 1.2E-144 |
| | Cldn23 | 1.294 | 0.734 | 0.615 | 0.0E+00 | 0.0E+00 | | Erp29 | 1.067 | 0.922 | 0.709 | 9.2E-146 | 1.4E-142 |
| | Chac1 | 1.291 | 0.750 | 0.706 | 0.0E+00 | 0.0E+00 | | Sdf2l1 | 0.667 | 0.902 | 0.718 | 1.0E-143 | 1.5E-140 |
| | Rdh10 | 1.290 | 0.787 | 0.518 | 0.0E+00 | 0.0E+00 | | Cd300c2 | 0.691 | 0.932 | 0.651 | 5.4E-142 | 8.1E-139 |
| | Fam57a | 1.279 | 0.733 | 0.754 | 0.0E+00 | 0.0E+00 | | Clec4a1 | 1.686 | 0.720 | 0.281 | 1.3E-141 | 1.9E-138 |
| | Gadd45b | 1.263 | 0.948 | 0.711 | 0.0E+00 | 0.0E+00 | | Ms4a6b | 1.104 | 0.755 | 0.271 | 1.1E-138 | 1.6E-135 |
| | Cops6 | 1.253 | 0.833 | 0.647 | 0.0E+00 | 0.0E+00 | | Tmem160 | 0.781 | 0.834 | 0.577 | 2.9E-137 | 4.4E-134 |
| | Krt8 | 1.231 | 0.690 | 0.700 | 0.0E+00 | 0.0E+00 | | H2-DMa | 1.252 | 0.756 | 0.394 | 3.1E-136 | 4.7E-133 |
| | Gadd45g | 1.225 | 0.659 | 0.573 | 0.0E+00 | 0.0E+00 | | Abhd12 | 0.845 | 0.790 | 0.432 | 6.7E-135 | 1.0E-131 |
| | Fam110a | 1.163 | 0.787 | 0.499 | 0.0E+00 | 0.0E+00 | | Hexb | 1.403 | 0.863 | 0.751 | 6.5E-132 | 9.7E-129 |
| | Pls3 | 1.149 | 0.784 | 0.594 | 0.0E+00 | 0.0E+00 | | S100a4 | 1.042 | 0.790 | 0.500 | 1.6E-130 | 2.3E-127 |
| | Hbegf | 1.114 | 0.731 | 0.762 | 0.0E+00 | 0.0E+00 | | Flna | 0.830 | 0.835 | 0.487 | 2.2E-128 | 3.3E-125 |

| | | | | | | | | | | | | | |
|---|---|---|---|---|---|---|---|---|---|---|---|---|---|
| | Phlda1 | 1.100 | 0.759 | 0.510 | 0.0E+00 | 0.0E+00 | | Unc93b1 | 0.755 | 0.848 | 0.511 | 1.4E-127 | 2.0E-124 |
| | Krt80 | 1.097 | 0.845 | 0.529 | 0.0E+00 | 0.0E+00 | | Pid1 | 0.802 | 0.765 | 0.585 | 1.5E-127 | 2.2E-124 |
| | Manf | 1.096 | 0.869 | 0.829 | 0.0E+00 | 0.0E+00 | | Rras | 0.921 | 0.760 | 0.544 | 7.2E-126 | 1.1E-122 |
| | Parm1 | 1.078 | 0.778 | 0.721 | 0.0E+00 | 0.0E+00 | | Itgb2 | 0.950 | 0.910 | 0.585 | 2.3E-124 | 3.5E-121 |
| | Hsd17b2 | 1.076 | 0.686 | 0.537 | 0.0E+00 | 0.0E+00 | | Lilra5 | 1.100 | 0.724 | 0.321 | 2.1E-120 | 3.2E-117 |
| | Slco2a1 | 1.075 | 0.676 | 0.528 | 0.0E+00 | 0.0E+00 | | Mgst1 | 0.710 | 0.874 | 0.601 | 1.1E-119 | 1.6E-116 |
| | 2610528A11Ril | 1.070 | 0.701 | 0.641 | 0.0E+00 | 0.0E+00 | | Selenop | 0.924 | 0.776 | 0.519 | 2.1E-115 | 3.2E-112 |
| | Apol7a | 1.050 | 0.698 | 0.453 | 0.0E+00 | 0.0E+00 | | mt-Nd1 | 0.722 | 0.963 | 0.666 | 1.7E-114 | 2.6E-111 |
| | Mlph | 1.045 | 0.754 | 0.665 | 0.0E+00 | 0.0E+00 | | Fcgrt | 0.886 | 0.735 | 0.282 | 7.0E-114 | 1.0E-110 |
| | Sqstm1 | 1.023 | 0.873 | 0.581 | 0.0E+00 | 0.0E+00 | | Plekho1 | 0.926 | 0.756 | 0.553 | 7.7E-112 | 1.2E-108 |
| | Pmaip1 | 0.991 | 0.742 | 0.669 | 0.0E+00 | 0.0E+00 | | Ndufa4 | 0.708 | 0.879 | 0.666 | 2.0E-111 | 3.0E-108 |
| | Oasl1 | 0.984 | 0.620 | 0.575 | 0.0E+00 | 0.0E+00 | | Mef2c | 1.025 | 0.724 | 0.384 | 3.3E-106 | 5.0E-103 |
| | Plac8 | 0.977 | 0.735 | 0.523 | 0.0E+00 | 0.0E+00 | | Irf8 | 1.231 | 0.760 | 0.635 | 2.2E-105 | 3.3E-102 |
| | Ifi202b | 0.963 | 0.703 | 0.648 | 0.0E+00 | 0.0E+00 | | Npc2 | 0.824 | 0.958 | 0.889 | 2.4E-101 | 3.7E-98 |
| | Rbp1 | 0.960 | 0.608 | 0.545 | 0.0E+00 | 0.0E+00 | | Plac8 | 1.352 | 0.759 | 0.557 | 6.2E-100 | 9.3E-97 |
| | 2010109I03Rik | 0.930 | 0.607 | 0.662 | 0.0E+00 | 0.0E+00 | | H2-DMb1 | 1.111 | 0.707 | 0.428 | 3.2E-94 | 4.9E-91 |
| | Tspan8 | 0.903 | 0.766 | 0.646 | 0.0E+00 | 0.0E+00 | | Tifab | 0.751 | 0.686 | 0.354 | 2.7E-81 | 4.1E-78 |
| | Yod1 | 0.898 | 0.649 | 0.569 | 0.0E+00 | 0.0E+00 | | Pltp | 0.669 | 0.703 | 0.365 | 9.6E-81 | 1.4E-77 |
| | Cald1 | 0.888 | 0.677 | 0.690 | 0.0E+00 | 0.0E+00 | | Ccdc88a | 1.007 | 0.688 | 0.547 | 2.3E-70 | 3.5E-67 |
| | Plk2 | 0.881 | 0.723 | 0.493 | 0.0E+00 | 0.0E+00 | | Ear2 | 1.003 | 0.647 | 0.320 | 8.7E-66 | 1.3E-62 |
| | Ephb3 | 0.877 | 0.717 | 0.631 | 0.0E+00 | 0.0E+00 | | Ms4a4c | 0.756 | 0.630 | 0.135 | 1.7E-64 | 2.5E-61 |
| | Oit1 | 0.851 | 0.751 | 0.566 | 0.0E+00 | 0.0E+00 | | Ace | 1.683 | 0.644 | 0.536 | 6.6E-64 | 9.9E-61 |
| | Hsp90aa1 | 0.846 | 0.911 | 0.832 | 0.0E+00 | 0.0E+00 | | Evl | 0.693 | 0.688 | 0.340 | 1.6E-61 | 2.4E-58 |
| | Klk8 | 0.932 | 0.547 | 0.488 | 6.8E-289 | 1.0E-285 | | Ifi27l2a | 0.784 | 0.697 | 0.528 | 8.9E-55 | 1.3E-51 |
| | Hsp90b1 | 1.266 | 0.827 | 0.781 | 2.1E-272 | 3.1E-269 | | Gm15987 | 0.810 | 0.635 | 0.261 | 1.1E-54 | 1.6E-51 |
| | Tsc22d1 | 0.850 | 0.640 | 0.773 | 8.6E-178 | 1.3E-174 | | Klf4 | 0.724 | 0.648 | 0.422 | 4.9E-44 | 7.3E-41 |
| | Clic5 | 1.114 | 0.581 | 0.733 | 7.7E-159 | 1.2E-155 | | Slc12a2 | 1.262 | 0.706 | 0.578 | 4.8E-43 | 7.2E-40 |
| | Ifit1bl1 | 1.446 | 0.487 | 0.470 | 6.4E-149 | 9.6E-146 | | Adgre4 | 0.930 | 0.597 | 0.375 | 2.9E-42 | 4.4E-39 |
| | Cxcl5 | 1.588 | 0.446 | 0.399 | 6.7E-125 | 1.0E-121 | | Bst2 | 0.727 | 0.651 | 0.506 | 1.1E-30 | 1.7E-27 |
| | Pigr | 0.995 | 0.441 | 0.469 | 6.3E-68 | 9.5E-65 | | Tmem176b | 1.272 | 0.622 | 0.568 | 1.4E-22 | 2.1E-19 |
| | Cxcl3 | 1.035 | 0.364 | 0.517 | 5.7E-03 | 1.0E+00 | | Krt80 | 0.745 | 0.634 | 0.587 | 1.1E-20 | 1.6E-17 |
| **Neu-1** | Cst3 | 1.532 | 0.929 | 0.800 | 0.0E+00 | 0.0E+00 | **B cells** | Igkc | 4.317 | 0.964 | 0.279 | 0.0E+00 | 0.0E+00 |
| | Gngt2 | 1.374 | 0.812 | 0.614 | 0.0E+00 | 0.0E+00 | | Ighm | 3.141 | 0.908 | 0.317 | 0.0E+00 | 0.0E+00 |
| | Id2 | 1.172 | 0.925 | 0.753 | 0.0E+00 | 0.0E+00 | | Cd74 | 3.059 | 0.992 | 0.554 | 0.0E+00 | 0.0E+00 |
| | Ptgs1 | 1.172 | 0.772 | 0.678 | 0.0E+00 | 0.0E+00 | | H2-Ab1 | 2.828 | 0.975 | 0.628 | 0.0E+00 | 0.0E+00 |
| | Laptm5 | 0.838 | 0.829 | 0.722 | 0.0E+00 | 0.0E+00 | | H2-Aa | 2.789 | 0.983 | 0.554 | 0.0E+00 | 0.0E+00 |
| | Gm19951 | 0.675 | 0.840 | 0.656 | 0.0E+00 | 0.0E+00 | | Cd79a | 2.550 | 0.908 | 0.408 | 0.0E+00 | 0.0E+00 |
| | Ccl4 | 0.552 | 0.895 | 0.685 | 0.0E+00 | 0.0E+00 | | Cd79b | 2.207 | 0.854 | 0.309 | 0.0E+00 | 0.0E+00 |
| | Hist1h1e | 0.491 | 0.794 | 0.590 | 0.0E+00 | 0.0E+00 | | Ebf1 | 2.190 | 0.862 | 0.157 | 0.0E+00 | 0.0E+00 |
| | Asprv1 | 0.446 | 0.778 | 0.552 | 0.0E+00 | 0.0E+00 | | Iglc2 | 2.133 | 0.870 | 0.147 | 0.0E+00 | 0.0E+00 |
| | Egr2 | 0.310 | 0.274 | 0.401 | 3.3E-290 | 5.0E-287 | | Cd72 | 1.409 | 0.846 | 0.159 | 0.0E+00 | 0.0E+00 |
| | B930036N10Ri | 0.499 | 0.642 | 0.468 | 1.2E-283 | 1.8E-280 | | Serpina3g | 0.465 | 0.972 | 0.560 | 0.0E+00 | 0.0E+00 |
| | Id1 | 1.043 | 0.832 | 0.665 | 4.7E-266 | 7.1E-263 | | Gatm | 0.355 | 0.992 | 0.516 | 0.0E+00 | 0.0E+00 |
| | Gpx1 | 0.848 | 0.869 | 0.830 | 3.8E-265 | 5.7E-262 | | Ighd | 2.304 | 0.799 | 0.270 | 1.4E-299 | 2.1E-296 |
| | Ms4a6d | 0.565 | 0.805 | 0.725 | 7.4E-254 | 1.1E-250 | | Gm15987 | 0.802 | 0.838 | 0.255 | 1.2E-273 | 1.9E-270 |
| | Csf1r | 0.366 | 0.280 | 0.483 | 9.3E-226 | 1.4E-222 | | Jchain | 1.341 | 0.826 | 0.370 | 9.8E-271 | 1.5E-267 |
| | Gpc3 | 0.286 | 0.688 | 0.670 | 3.7E-198 | 5.5E-195 | | Ms4a1 | 1.914 | 0.819 | 0.197 | 6.8E-270 | 1.0E-266 |
| | Cd300c2 | 1.210 | 0.716 | 0.649 | 5.2E-190 | 7.8E-187 | | Ly6d | 0.446 | 0.929 | 0.779 | 5.9E-245 | 8.9E-242 |
| | Hist1h1d | 0.253 | 0.697 | 0.533 | 8.4E-188 | 1.3E-184 | | Rpl3 | 1.155 | 0.991 | 0.707 | 7.0E-233 | 1.1E-229 |
| | Ptgs2 | 0.526 | 0.234 | 0.306 | 4.4E-180 | 6.6E-177 | | Mef2c | 1.562 | 0.830 | 0.381 | 8.7E-231 | 1.3E-227 |
| | Pmaip1 | 0.518 | 0.787 | 0.662 | 1.1E-178 | 1.6E-175 | | Bank1 | 1.757 | 0.792 | 0.348 | 2.6E-210 | 3.9E-207 |
| | Ccl3 | 0.592 | 0.738 | 0.624 | 1.2E-148 | 1.8E-145 | | Ptma | 0.989 | 0.991 | 0.725 | 2.2E-205 | 3.3E-202 |
| | Osgin1 | 0.724 | 0.685 | 0.570 | 7.6E-146 | 1.1E-142 | | Igha | 1.645 | 0.780 | 0.378 | 1.0E-200 | 1.5E-197 |
| | Olr1 | 0.348 | 0.328 | 0.404 | 8.3E-139 | 1.2E-135 | | Ighg1 | 1.025 | 0.772 | 0.364 | 8.7E-190 | 1.3E-186 |
| | Pi16 | 0.312 | 0.619 | 0.443 | 4.3E-134 | 6.5E-131 | | H2-DMa | 0.962 | 0.810 | 0.392 | 7.6E-178 | 1.1E-174 |
| | Hexb | 0.810 | 0.794 | 0.746 | 4.2E-133 | 6.3E-130 | | Fam96a | 0.433 | 0.939 | 0.556 | 5.6E-177 | 8.5E-174 |
| | Fcgr4 | 0.354 | 0.750 | 0.685 | 1.7E-126 | 2.6E-123 | | Gpr65 | 0.440 | 0.838 | 0.448 | 3.8E-174 | 5.7E-171 |
| | Bcl2a1d | 0.547 | 0.380 | 0.543 | 1.6E-123 | 2.4E-120 | | Iglc3 | 1.588 | 0.712 | 0.157 | 2.9E-168 | 4.3E-165 |
| | Cd63 | 0.327 | 0.866 | 0.768 | 2.9E-120 | 4.4E-117 | | Napsa | 1.338 | 0.843 | 0.582 | 3.3E-163 | 5.0E-160 |
| | Ier3 | 0.771 | 0.802 | 0.786 | 5.3E-113 | 8.0E-110 | | mt-Nd1 | 0.665 | 0.967 | 0.666 | 1.3E-152 | 1.9E-149 |
| | Atp1a1 | 0.853 | 0.779 | 0.694 | 1.1E-110 | 1.7E-107 | | Ets1 | 1.116 | 0.792 | 0.447 | 1.3E-150 | 1.9E-147 |
| | Hk2 | 0.263 | 0.470 | 0.609 | 5.4E-104 | 8.0E-101 | | Mzb1 | 1.617 | 0.728 | 0.330 | 2.9E-146 | 4.3E-143 |
| | P2ry6 | 0.401 | 0.710 | 0.682 | 1.5E-100 | 2.3E-97 | | Rps2 | 0.667 | 0.989 | 0.892 | 8.8E-135 | 1.3E-131 |
| | Lrg1 | 0.265 | 0.765 | 0.686 | 1.4E-95 | 2.1E-92 | | Fcmr | 1.153 | 0.750 | 0.392 | 8.3E-132 | 1.2E-128 |
| | Gm12840 | 1.099 | 0.321 | 0.411 | 2.0E-95 | 3.0E-92 | | Gm31243 | 1.017 | 0.709 | 0.181 | 9.7E-121 | 1.5E-117 |
| | H1f0 | 0.599 | 0.700 | 0.568 | 2.2E-87 | 3.3E-84 | | H2-DMb2 | 1.138 | 0.718 | 0.272 | 2.2E-119 | 3.3E-116 |
| | Naaa | 0.359 | 0.655 | 0.568 | 4.8E-75 | 7.2E-72 | | Ranbp1 | 0.695 | 0.826 | 0.642 | 4.2E-105 | 6.4E-102 |
| | Ltc4s | 0.796 | 0.644 | 0.599 | 1.2E-72 | 1.9E-69 | | Gm8369 | 0.449 | 0.297 | 0.105 | 1.4E-101 | 2.1E-98 |
| | Reep5 | 0.646 | 0.749 | 0.711 | 5.2E-55 | 7.9E-52 | | Gimap6 | 1.256 | 0.710 | 0.321 | 1.5E-98 | 2.3E-95 |
| | Egr1 | 0.842 | 0.617 | 0.544 | 1.1E-51 | 1.6E-48 | | 4930523C07R | 1.050 | 0.716 | 0.510 | 6.0E-92 | 9.0E-89 |
| | Asah1 | 0.261 | 0.666 | 0.555 | 1.3E-49 | 2.0E-46 | | Ptprcap | 0.932 | 0.716 | 0.524 | 7.3E-82 | 1.1E-78 |
| | Gm2a | 0.855 | 0.637 | 0.632 | 1.0E-45 | 1.5E-42 | | Cd55 | 0.849 | 0.734 | 0.433 | 3.2E-81 | 4.8E-78 |
| | Hist1h4i | 1.255 | 0.646 | 0.671 | 1.8E-45 | 2.8E-42 | | Fcer2a | 1.434 | 0.669 | 0.501 | 4.6E-75 | 6.9E-72 |
| | Chil1 | 0.513 | 0.593 | 0.475 | 2.1E-43 | 3.2E-40 | | Erp29 | 0.518 | 0.910 | 0.710 | 1.4E-73 | 2.1E-70 |
| | Tgm2 | 0.664 | 0.463 | 0.641 | 5.8E-43 | 8.6E-40 | | Pax5 | 0.985 | 0.641 | 0.212 | 3.4E-73 | 5.1E-70 |

| | | | | | | |
|---|---|---|---|---|---|---|
| | Dpep2 | 0.374 | 0.652 | 0.687 | 7.8E-43 | 1.2E-39 |
| | Ctsa | 0.381 | 0.744 | 0.697 | 2.0E-42 | 2.9E-39 |
| | Fgd4 | 0.251 | 0.688 | 0.624 | 1.5E-41 | 2.3E-38 |
| | Fam20c | 0.442 | 0.792 | 0.763 | 5.8E-41 | 8.7E-38 |
| | Pou2f2 | 0.312 | 0.387 | 0.444 | 1.9E-34 | 2.9E-31 |
| | Cks2 | 1.189 | 0.534 | 0.460 | 3.2E-33 | 4.8E-30 |
| | Rhoc | 0.380 | 0.529 | 0.605 | 3.6E-33 | 5.3E-30 |
| | Fyb | 0.514 | 0.751 | 0.686 | 7.6E-32 | 1.1E-28 |
| | Fcgr2b | 0.347 | 0.713 | 0.648 | 2.8E-31 | 4.1E-28 |
| | C3 | 0.875 | 0.585 | 0.551 | 5.4E-30 | 8.1E-27 |
| | Slc7a11 | 0.272 | 0.428 | 0.478 | 6.5E-28 | 9.7E-25 |
| | Hist1h1c | 0.763 | 0.664 | 0.606 | 5.7E-26 | 8.5E-23 |
| | Tcirg1 | 0.280 | 0.550 | 0.636 | 9.7E-26 | 1.5E-22 |
| | Abca1 | 0.438 | 0.768 | 0.731 | 1.2E-25 | 1.7E-22 |
| | Rgs1 | 1.284 | 0.650 | 0.628 | 1.6E-25 | 2.4E-22 |
| | Csf2rb | 0.693 | 0.676 | 0.689 | 3.6E-25 | 5.5E-22 |
| | Bcl2a1b | 0.872 | 0.640 | 0.631 | 2.0E-23 | 2.9E-20 |
| | Ccng1 | 0.266 | 0.520 | 0.595 | 7.3E-23 | 1.1E-19 |
| | Agap1 | 0.867 | 0.586 | 0.610 | 2.6E-22 | 3.9E-19 |
| | Erp29 | 0.301 | 0.600 | 0.739 | 1.2E-21 | 1.7E-18 |
| | Slc6a6 | 0.338 | 0.463 | 0.522 | 2.5E-21 | 3.7E-18 |
| | Tnf | 0.725 | 0.327 | 0.308 | 9.4E-19 | 1.4E-15 |
| | Hist1h2ap | 0.263 | 0.508 | 0.479 | 6.2E-18 | 9.3E-15 |
| | Gm20186 | 0.564 | 0.404 | 0.440 | 6.6E-18 | 9.9E-15 |
| | Cfp | 0.354 | 0.543 | 0.434 | 1.4E-17 | 2.2E-14 |
| | Unc93b1 | 0.643 | 0.570 | 0.512 | 3.2E-16 | 4.8E-13 |
| | Ptma | 0.282 | 0.760 | 0.728 | 3.5E-15 | 5.2E-12 |
| | Itgax | 0.746 | 0.468 | 0.563 | 2.5E-14 | 3.8E-11 |
| | Aprt | 0.365 | 0.639 | 0.761 | 2.1E-12 | 3.1E-09 |
| | Itgb2 | 0.303 | 0.503 | 0.614 | 2.2E-12 | 3.3E-09 |
| | Ucp2 | 0.280 | 0.743 | 0.732 | 4.5E-11 | 6.8E-08 |
| | Bcl2a1a | 0.956 | 0.592 | 0.547 | 7.5E-10 | 1.1E-06 |
| | Arhgap25 | 0.475 | 0.546 | 0.497 | 4.7E-08 | 7.1E-05 |
| | G0s2 | 0.960 | 0.524 | 0.494 | 1.9E-06 | 2.8E-03 |
| | Gm26870 | 0.339 | 0.577 | 0.569 | 5.6E-05 | 8.3E-02 |
| | Cdkn1a | 0.436 | 0.825 | 0.760 | 5.5E-04 | 8.2E-01 |
| | Tgfbi | 0.656 | 0.586 | 0.616 | 5.7E-04 | 8.5E-01 |
| | Mpeg1 | 0.405 | 0.595 | 0.629 | 6.4E-04 | 9.6E-01 |
| | Csf2ra | 0.674 | 0.543 | 0.575 | 1.5E-03 | 1.0E+00 |
| | Tnfrsf23 | 0.465 | 0.654 | 0.712 | 5.8E-03 | 1.0E+00 |
| **Neu-2** | Gm5483 | 2.762 | 0.938 | 0.564 | 0.0E+00 | 0.0E+00 |
| | BC100530 | 2.671 | 0.815 | 0.520 | 0.0E+00 | 0.0E+00 |
| | Wfdc17 | 2.485 | 0.993 | 0.744 | 0.0E+00 | 0.0E+00 |
| | Cxcl2 | 2.340 | 0.951 | 0.469 | 0.0E+00 | 0.0E+00 |
| | Ifitm1 | 2.087 | 0.963 | 0.701 | 0.0E+00 | 0.0E+00 |
| | Retnlg | 2.057 | 0.960 | 0.576 | 0.0E+00 | 0.0E+00 |
| | Wfdc21 | 1.768 | 0.945 | 0.605 | 0.0E+00 | 0.0E+00 |
| | Lrg1 | 1.760 | 0.929 | 0.661 | 0.0E+00 | 0.0E+00 |
| | G0s2 | 1.756 | 0.819 | 0.446 | 0.0E+00 | 0.0E+00 |
| | Egr1 | 1.705 | 0.912 | 0.497 | 0.0E+00 | 0.0E+00 |
| | Lcn2 | 1.667 | 0.911 | 0.561 | 0.0E+00 | 0.0E+00 |
| | Ier3 | 1.643 | 0.972 | 0.758 | 0.0E+00 | 0.0E+00 |
| | Slpi | 1.397 | 0.904 | 0.602 | 0.0E+00 | 0.0E+00 |
| | Adam8 | 1.286 | 0.729 | 0.572 | 0.0E+00 | 0.0E+00 |
| | Ccl4 | 1.276 | 0.981 | 0.676 | 0.0E+00 | 0.0E+00 |
| | F630028O10Ri | 1.234 | 0.798 | 0.503 | 0.0E+00 | 0.0E+00 |
| | Slfn4 | 1.160 | 0.942 | 0.505 | 0.0E+00 | 0.0E+00 |
| | Id1 | 0.989 | 0.915 | 0.655 | 0.0E+00 | 0.0E+00 |
| | Csf2rb | 0.976 | 0.921 | 0.648 | 0.0E+00 | 0.0E+00 |
| | Steap4 | 0.965 | 0.732 | 0.447 | 0.0E+00 | 0.0E+00 |
| | Hcar2 | 0.955 | 0.882 | 0.648 | 0.0E+00 | 0.0E+00 |
| | Prok2 | 0.868 | 0.790 | 0.548 | 0.0E+00 | 0.0E+00 |
| | Olfm4 | 0.701 | 0.793 | 0.426 | 0.0E+00 | 0.0E+00 |
| | Dgat2 | 0.563 | 0.795 | 0.603 | 0.0E+00 | 0.0E+00 |
| | Tceal9 | 0.557 | 0.888 | 0.641 | 0.0E+00 | 0.0E+00 |
| | Cxcl3 | 0.533 | 0.840 | 0.429 | 0.0E+00 | 0.0E+00 |
| | Cd177 | 0.461 | 0.740 | 0.458 | 0.0E+00 | 0.0E+00 |
| | Tnfrsf23 | 0.440 | 0.910 | 0.668 | 0.0E+00 | 0.0E+00 |
| | Hk2 | 0.430 | 0.884 | 0.536 | 0.0E+00 | 0.0E+00 |
| | Ccl6 | 0.398 | 0.910 | 0.592 | 0.0E+00 | 0.0E+00 |
| | Cdkn1a | 0.298 | 0.945 | 0.742 | 0.0E+00 | 0.0E+00 |
| | Rgcc | 0.260 | 0.926 | 0.710 | 0.0E+00 | 0.0E+00 |
| | Slc7a11 | 0.868 | 0.715 | 0.429 | 1.1E-307 | 1.6E-304 |
| | Mmp8 | 0.729 | 0.731 | 0.597 | 8.0E-292 | 1.2E-288 |
| | Stfa2l1 | 2.238 | 0.682 | 0.448 | 1.4E-272 | 2.1E-269 |

| | | | | | | |
|---|---|---|---|---|---|---|
| | Ndufa4 | 0.509 | 0.843 | 0.667 | 4.6E-69 | 6.9E-66 |
| | Sdc4 | 0.893 | 0.745 | 0.674 | 2.2E-67 | 3.3E-64 |
| | Tnfrsf13c | 0.888 | 0.622 | 0.177 | 2.7E-67 | 4.1E-64 |
| | Scd1 | 1.063 | 0.661 | 0.348 | 8.6E-65 | 1.3E-61 |
| | Hmgn1 | 0.833 | 0.723 | 0.652 | 7.2E-61 | 1.1E-57 |
| | Hspe1 | 0.691 | 0.777 | 0.674 | 5.8E-60 | 8.7E-57 |
| | Hes1 | 0.392 | 0.338 | 0.532 | 7.1E-59 | 1.1E-55 |
| | Pxdc1 | 0.899 | 0.660 | 0.339 | 2.1E-58 | 3.2E-55 |
| | Cd180 | 0.448 | 0.352 | 0.326 | 1.2E-54 | 1.8E-51 |
| | Pgls | 0.647 | 0.753 | 0.634 | 2.2E-54 | 3.3E-51 |
| | Nap1l1 | 0.555 | 0.726 | 0.561 | 1.5E-44 | 2.2E-41 |
| | Blk | 0.482 | 0.380 | 0.328 | 1.6E-43 | 2.3E-40 |
| | Bcar3 | 0.432 | 0.387 | 0.408 | 3.1E-43 | 4.7E-40 |
| | Gimap3 | 0.924 | 0.632 | 0.218 | 1.3E-42 | 2.0E-39 |
| | AC149090.1 | 1.234 | 0.659 | 0.455 | 1.3E-41 | 1.9E-38 |
| | Irf8 | 0.826 | 0.671 | 0.638 | 8.6E-38 | 1.3E-34 |
| | Pou2f2 | 0.590 | 0.656 | 0.428 | 5.8E-36 | 8.7E-33 |
| | Ms4a4c | 0.498 | 0.386 | 0.143 | 8.2E-36 | 1.2E-32 |
| | Tubb5 | 0.433 | 0.727 | 0.599 | 1.1E-33 | 1.7E-30 |
| | Ly86 | 0.735 | 0.619 | 0.293 | 8.3E-27 | 1.2E-23 |
| | Aldh2 | 0.584 | 0.639 | 0.517 | 2.6E-26 | 3.9E-23 |
| | S1pr1 | 0.415 | 0.391 | 0.514 | 1.3E-24 | 1.9E-21 |
| | Id3 | 0.855 | 0.605 | 0.569 | 1.8E-24 | 2.7E-21 |
| | Pou2af1 | 1.186 | 0.585 | 0.245 | 5.4E-24 | 8.1E-21 |
| | Lmo2 | 0.870 | 0.616 | 0.400 | 1.0E-23 | 1.5E-20 |
| | Gimap1 | 0.903 | 0.615 | 0.455 | 3.2E-23 | 4.9E-20 |
| | C1qbp | 0.416 | 0.684 | 0.663 | 3.0E-22 | 4.5E-19 |
| | Slc25a4 | 0.458 | 0.662 | 0.537 | 2.2E-21 | 3.3E-18 |
| | Gimap7 | 0.681 | 0.581 | 0.231 | 4.9E-21 | 7.3E-18 |
| | Myo1e | 0.370 | 0.454 | 0.610 | 3.7E-20 | 5.5E-17 |
| | Siglecg | 0.415 | 0.590 | 0.505 | 4.4E-17 | 6.6E-14 |
| | Eprs | 0.470 | 0.654 | 0.584 | 2.2E-16 | 3.3E-13 |
| | Nucks1 | 0.526 | 0.636 | 0.534 | 2.7E-15 | 4.0E-12 |
| | Plekho1 | 0.670 | 0.621 | 0.557 | 1.2E-14 | 1.9E-11 |
| | Zbtb20 | 0.422 | 0.454 | 0.546 | 7.7E-14 | 1.2E-10 |
| | Evl | 0.561 | 0.600 | 0.343 | 3.7E-13 | 5.6E-10 |
| | Snx5 | 0.608 | 0.621 | 0.551 | 2.3E-12 | 3.5E-09 |
| | Ciita | 0.638 | 0.441 | 0.281 | 1.7E-11 | 2.6E-08 |
| | Blnk | 0.663 | 0.596 | 0.738 | 2.1E-10 | 3.2E-07 |
| | Sptbn1 | 0.654 | 0.623 | 0.580 | 6.2E-10 | 9.3E-07 |
| | Iglc1 | 1.620 | 0.546 | 0.204 | 9.9E-09 | 1.5E-05 |
| | Ms4a6c | 0.618 | 0.593 | 0.418 | 1.4E-08 | 2.1E-05 |
| | Cd2ap | 0.534 | 0.624 | 0.597 | 1.7E-08 | 2.6E-05 |
| | Cd69 | 0.605 | 0.558 | 0.334 | 5.9E-08 | 8.8E-05 |
| | Ms4a6b | 0.387 | 0.469 | 0.280 | 2.3E-07 | 3.4E-04 |
| | Bin1 | 0.731 | 0.578 | 0.566 | 2.8E-07 | 4.2E-04 |
| | Kcnq1ot1 | 0.583 | 0.624 | 0.657 | 3.6E-07 | 5.4E-04 |
| | Odc1 | 0.433 | 0.478 | 0.572 | 3.6E-07 | 5.4E-04 |
| | Vpreb3 | 1.232 | 0.539 | 0.135 | 4.7E-07 | 7.1E-04 |
| | Nr4a1 | 0.505 | 0.424 | 0.317 | 1.1E-06 | 1.7E-03 |
| | Rftn1 | 0.435 | 0.495 | 0.627 | 2.7E-06 | 4.1E-03 |
| | Pdia4 | 0.591 | 0.566 | 0.537 | 7.0E-06 | 1.1E-02 |
| | Gpr171 | 0.531 | 0.481 | 0.578 | 3.9E-05 | 5.9E-02 |
| | Ikzf3 | 0.915 | 0.544 | 0.267 | 4.1E-05 | 6.2E-02 |
| | Gpr183 | 0.771 | 0.535 | 0.267 | 4.2E-04 | 6.2E-01 |
| | Cd38 | 0.484 | 0.588 | 0.584 | 8.6E-03 | 1.0E+00 |
| **Lung Ep.** | Il33 | 1.978 | 0.892 | 0.537 | 0.0E+00 | 0.0E+00 |
| | Areg | 1.182 | 0.853 | 0.425 | 1.7E-291 | 2.6E-288 |
| | Fxyd3 | 2.925 | 0.930 | 0.612 | 1.3E-275 | 2.0E-272 |
| | Wfdc2 | 4.063 | 0.911 | 0.760 | 1.7E-275 | 2.5E-272 |
| | Col1a1 | 1.424 | 0.721 | 0.220 | 4.9E-249 | 7.3E-246 |
| | Dapl1 | 1.474 | 0.823 | 0.391 | 1.1E-243 | 1.6E-240 |
| | Col1a2 | 1.973 | 0.756 | 0.381 | 8.7E-239 | 1.3E-235 |
| | Btbd3 | 1.083 | 0.834 | 0.570 | 1.3E-220 | 1.9E-217 |
| | Ly6d | 1.756 | 0.948 | 0.779 | 8.8E-220 | 1.3E-216 |
| | Hspb1 | 1.934 | 0.823 | 0.582 | 6.5E-217 | 9.7E-214 |
| | Wfdc3 | 1.449 | 0.821 | 0.682 | 5.9E-205 | 8.8E-202 |
| | Ckmt1 | 1.370 | 0.883 | 0.732 | 1.1E-200 | 1.7E-197 |
| | Gpx2 | 1.928 | 0.819 | 0.571 | 1.7E-198 | 2.6E-195 |
| | Cdh1 | 1.240 | 0.859 | 0.604 | 1.2E-197 | 1.8E-194 |
| | Tmem176a | 2.040 | 0.873 | 0.582 | 7.6E-197 | 1.1E-193 |
| | Ehf | 2.134 | 0.855 | 0.799 | 5.1E-196 | 7.6E-193 |
| | Gsta4 | 2.610 | 0.871 | 0.705 | 1.8E-191 | 2.7E-188 |
| | Them5 | 1.260 | 0.732 | 0.268 | 1.7E-187 | 2.5E-184 |
| | Cldn7 | 1.072 | 0.896 | 0.684 | 1.1E-182 | 1.6E-179 |

| | | | | | | |
|---|---|---|---|---|---|---|
| | Osgin1 | 0.516 | 0.765 | 0.560 | 5.6E-269 | 8.5E-266 |
| | Stfa2 | 1.396 | 0.680 | 0.497 | 3.6E-261 | 5.4E-258 |
| | Il1f9 | 0.937 | 0.718 | 0.534 | 1.0E-248 | 1.6E-245 |
| | Gadd45a | 0.936 | 0.731 | 0.606 | 1.1E-241 | 1.7E-238 |
| | Ifitm3 | 0.397 | 0.844 | 0.679 | 2.8E-228 | 4.3E-225 |
| | Hip1 | 0.280 | 0.812 | 0.586 | 5.1E-221 | 7.7E-218 |
| | Ifitm6 | 0.479 | 0.683 | 0.435 | 1.5E-216 | 2.2E-213 |
| | Chil1 | 0.781 | 0.691 | 0.462 | 2.0E-216 | 2.9E-213 |
| | Asprv1 | 1.206 | 0.707 | 0.570 | 1.3E-199 | 1.9E-196 |
| | Tgm2 | 0.428 | 0.758 | 0.587 | 5.2E-159 | 7.8E-156 |
| | Acod1 | 0.849 | 0.624 | 0.389 | 1.6E-127 | 2.5E-124 |
| | Fyb | 0.331 | 0.786 | 0.682 | 1.7E-118 | 2.6E-115 |
| | Saa3 | 0.560 | 0.748 | 0.606 | 4.7E-83 | 7.1E-80 |
| | Ly6g | 0.520 | 0.596 | 0.425 | 4.1E-82 | 6.1E-79 |
| | Tgfbi | 0.391 | 0.696 | 0.596 | 1.9E-80 | 2.8E-77 |
| | Gm5416 | 0.363 | 0.605 | 0.486 | 5.7E-76 | 8.5E-73 |
| | Pi16 | 0.460 | 0.605 | 0.450 | 1.1E-70 | 1.6E-67 |
| | Fcgr4 | 0.341 | 0.715 | 0.693 | 1.4E-60 | 2.0E-57 |
| | Slc6a6 | 0.335 | 0.651 | 0.489 | 4.4E-51 | 6.6E-48 |
| | Csf2ra | 0.339 | 0.671 | 0.553 | 1.6E-47 | 2.4E-44 |
| | Fgd4 | 0.308 | 0.677 | 0.628 | 5.9E-39 | 8.9E-36 |
| | Stfa3 | 0.456 | 0.528 | 0.544 | 1.1E-08 | 1.7E-05 |
| | Hacd4 | 0.323 | 0.496 | 0.524 | 8.5E-08 | 1.3E-04 |
| | Ptgs2 | 0.592 | 0.407 | 0.275 | 1.3E-07 | 2.0E-04 |
| | Fosb | 0.422 | 0.476 | 0.366 | 3.8E-03 | 1.0E+00 |
| **Neu-3** | Ccl3 | 2.965 | 0.920 | 0.610 | 0.0E+00 | 0.0E+00 |
| | Hcar2 | 2.705 | 0.884 | 0.657 | 0.0E+00 | 0.0E+00 |
| | Ccl4 | 2.320 | 0.868 | 0.702 | 0.0E+00 | 0.0E+00 |
| | Nceh1 | 2.161 | 0.856 | 0.686 | 0.0E+00 | 0.0E+00 |
| | Gadd45b | 2.109 | 0.877 | 0.741 | 0.0E+00 | 0.0E+00 |
| | Ifrd1 | 1.995 | 0.819 | 0.702 | 0.0E+00 | 0.0E+00 |
| | Hilpda | 1.887 | 0.836 | 0.728 | 0.0E+00 | 0.0E+00 |
| | Atp6v1c1 | 1.859 | 0.857 | 0.741 | 0.0E+00 | 0.0E+00 |
| | Zeb2 | 1.831 | 0.829 | 0.607 | 0.0E+00 | 0.0E+00 |
| | Cd63 | 1.769 | 0.936 | 0.766 | 0.0E+00 | 0.0E+00 |
| | Ftl1 | 1.745 | 1.000 | 0.953 | 0.0E+00 | 0.0E+00 |
| | Id2 | 1.629 | 0.874 | 0.771 | 0.0E+00 | 0.0E+00 |
| | F10 | 1.608 | 0.863 | 0.694 | 0.0E+00 | 0.0E+00 |
| | Ctsz | 1.414 | 0.936 | 0.780 | 0.0E+00 | 0.0E+00 |
| | Ctsb | 1.241 | 0.977 | 0.941 | 0.0E+00 | 0.0E+00 |
| | Gas2l3 | 1.116 | 0.818 | 0.634 | 0.0E+00 | 0.0E+00 |
| | Dhfr | 0.942 | 0.789 | 0.739 | 0.0E+00 | 0.0E+00 |
| | Lhfpl2 | 0.915 | 0.784 | 0.664 | 0.0E+00 | 0.0E+00 |
| | Ctsd | 0.494 | 0.987 | 0.963 | 0.0E+00 | 0.0E+00 |
| | Fcgr2b | 0.726 | 0.838 | 0.637 | 1.2E-307 | 1.8E-304 |
| | Chka | 0.537 | 0.738 | 0.519 | 3.0E-302 | 4.5E-299 |
| | H2-Eb1 | 1.000 | 0.683 | 0.519 | 1.0E-285 | 1.5E-282 |
| | Cd274 | 1.549 | 0.725 | 0.579 | 2.4E-265 | 3.6E-262 |
| | Plcxd2 | 0.637 | 0.728 | 0.653 | 3.2E-265 | 4.8E-262 |
| | Hexa | 0.947 | 0.839 | 0.688 | 1.1E-248 | 1.7E-245 |
| | Dock10 | 0.893 | 0.783 | 0.666 | 2.7E-239 | 4.1E-236 |
| | Pdxk | 0.597 | 0.726 | 0.574 | 7.5E-236 | 1.1E-232 |
| | Cxcl2 | 1.230 | 0.759 | 0.512 | 6.9E-233 | 1.0E-229 |
| | Psap | 0.760 | 0.852 | 0.749 | 1.3E-230 | 1.9E-227 |
| | Gns | 1.339 | 0.782 | 0.688 | 1.3E-230 | 2.0E-227 |
| | Lamp1 | 1.194 | 0.885 | 0.861 | 2.6E-229 | 3.9E-226 |
| | Atf3 | 1.754 | 0.679 | 0.548 | 3.1E-224 | 4.7E-221 |
| | Npc1 | 1.211 | 0.749 | 0.677 | 5.2E-220 | 7.8E-217 |
| | Ccnf | 0.536 | 0.662 | 0.612 | 4.8E-210 | 7.3E-207 |
| | Plekhm2 | 1.096 | 0.785 | 0.726 | 1.7E-206 | 2.5E-203 |
| | Rgs1 | 1.675 | 0.774 | 0.615 | 1.1E-205 | 1.6E-202 |
| | Ier3 | 1.085 | 0.810 | 0.786 | 1.8E-195 | 2.7E-192 |
| | 9130230L23Rik | 0.335 | 0.648 | 0.524 | 1.2E-192 | 1.8E-189 |
| | P2rx7 | 0.626 | 0.760 | 0.609 | 9.9E-177 | 1.5E-173 |
| | Fam20c | 0.499 | 0.847 | 0.758 | 1.7E-170 | 2.6E-167 |
| | Gstm1 | 0.562 | 0.697 | 0.550 | 1.0E-164 | 1.6E-161 |
| | Hmox1 | 1.389 | 0.790 | 0.754 | 7.4E-133 | 1.1E-129 |
| | Aprt | 0.932 | 0.820 | 0.731 | 3.6E-132 | 5.4E-129 |
| | Tnfrsf12a | 0.301 | 0.721 | 0.595 | 1.8E-125 | 2.7E-122 |
| | Tpp1 | 0.909 | 0.716 | 0.608 | 4.3E-113 | 6.5E-110 |
| | Inhba | 0.910 | 0.622 | 0.550 | 4.4E-112 | 6.5E-109 |
| | Tst | 0.398 | 0.688 | 0.590 | 1.2E-95 | 1.9E-92 |
| | Tcirg1 | 1.119 | 0.670 | 0.616 | 9.4E-90 | 1.4E-86 |
| | Canx | 0.712 | 0.769 | 0.666 | 1.2E-87 | 1.8E-84 |
| | Mpeg1 | 0.696 | 0.701 | 0.614 | 6.5E-80 | 9.7E-77 |

| | | | | | | |
|---|---|---|---|---|---|---|
| | Clu | 2.509 | 0.912 | 0.787 | 1.7E-182 | 2.5E-179 |
| | Avpi1 | 1.828 | 0.900 | 0.732 | 2.4E-182 | 3.6E-179 |
| | Cp | 1.550 | 0.766 | 0.560 | 1.4E-168 | 2.1E-165 |
| | Cald1 | 1.304 | 0.837 | 0.684 | 1.2E-164 | 1.8E-161 |
| | Mgp | 2.757 | 0.817 | 0.644 | 3.8E-162 | 5.7E-159 |
| | Cbr2 | 3.030 | 0.806 | 0.745 | 6.7E-160 | 1.0E-156 |
| | Ybx3 | 1.340 | 0.857 | 0.598 | 6.7E-160 | 1.0E-156 |
| | Far1 | 1.422 | 0.882 | 0.509 | 6.9E-160 | 1.0E-156 |
| | Plac8 | 1.530 | 0.919 | 0.554 | 8.7E-160 | 1.3E-156 |
| | Serping1 | 1.481 | 0.693 | 0.306 | 5.8E-159 | 8.7E-156 |
| | Akr1b3 | 1.620 | 0.866 | 0.517 | 1.5E-157 | 2.2E-154 |
| | Cd81 | 1.439 | 0.922 | 0.607 | 1.0E-155 | 1.5E-152 |
| | Scgb1a1 | 3.980 | 0.915 | 0.811 | 7.6E-155 | 1.1E-151 |
| | Id3 | 1.470 | 0.769 | 0.565 | 1.6E-149 | 2.5E-146 |
| | Perp | 1.659 | 0.834 | 0.733 | 1.2E-147 | 1.8E-144 |
| | Ppp1r14b | 1.481 | 0.892 | 0.660 | 6.9E-146 | 1.0E-142 |
| | Tpm2 | 1.173 | 0.815 | 0.658 | 4.9E-138 | 7.4E-135 |
| | Ceacam1 | 1.300 | 0.918 | 0.689 | 2.4E-135 | 3.5E-132 |
| | Igfbp4 | 1.213 | 0.714 | 0.516 | 3.6E-125 | 5.4E-122 |
| | Rpl3 | 1.482 | 0.892 | 0.712 | 1.4E-118 | 2.1E-115 |
| | Ptprf | 1.412 | 0.746 | 0.622 | 2.3E-117 | 3.4E-114 |
| | Rps2 | 1.263 | 0.934 | 0.894 | 1.5E-115 | 2.2E-112 |
| | Nenf | 1.188 | 0.790 | 0.520 | 6.5E-109 | 9.7E-106 |
| | Arg1 | 1.174 | 0.919 | 0.760 | 1.7E-105 | 2.6E-102 |
| | Nedd4 | 1.526 | 0.696 | 0.323 | 1.8E-105 | 2.8E-102 |
| | Dmkn | 1.828 | 0.776 | 0.719 | 1.0E-104 | 1.6E-101 |
| | Fabp5 | 2.134 | 0.923 | 0.803 | 7.8E-103 | 1.2E-99 |
| | Gm26870 | 1.224 | 0.813 | 0.564 | 1.2E-101 | 1.8E-98 |
| | Eln | 1.022 | 0.671 | 0.350 | 3.1E-100 | 4.6E-97 |
| | Scgb3a1 | 4.567 | 0.740 | 0.635 | 1.2E-99 | 1.8E-96 |
| | Pam | 1.544 | 0.788 | 0.651 | 1.9E-97 | 2.9E-94 |
| | Aqp5 | 1.174 | 0.777 | 0.728 | 4.3E-96 | 6.5E-93 |
| | Phlda1 | 1.105 | 0.823 | 0.551 | 4.7E-95 | 7.1E-92 |
| | Prdx2 | 1.395 | 0.805 | 0.642 | 2.1E-89 | 3.1E-86 |
| | Epcam | 1.512 | 0.786 | 0.644 | 4.0E-89 | 6.0E-86 |
| | Dcxr | 1.400 | 0.768 | 0.619 | 4.5E-87 | 6.8E-84 |
| | Serpinb6b | 1.063 | 0.769 | 0.688 | 5.7E-86 | 8.6E-83 |
| | Rbp1 | 1.619 | 0.722 | 0.552 | 4.9E-85 | 7.3E-82 |
| | Muc5b | 2.403 | 0.675 | 0.506 | 7.2E-84 | 1.1E-80 |
| | Scd1 | 1.002 | 0.226 | 0.360 | 1.4E-83 | 2.2E-80 |
| | Tsc22d1 | 1.408 | 0.780 | 0.747 | 1.2E-81 | 1.9E-78 |
| | Socs2 | 1.111 | 0.732 | 0.667 | 2.5E-80 | 3.8E-77 |
| | Selenbp1 | 2.144 | 0.707 | 0.645 | 4.2E-80 | 6.3E-77 |
| | Krt8 | 1.959 | 0.720 | 0.698 | 5.2E-77 | 7.9E-74 |
| | Atp5g1 | 1.265 | 0.834 | 0.645 | 8.9E-76 | 1.3E-72 |
| | Prdx1 | 1.233 | 0.822 | 0.667 | 1.8E-74 | 2.6E-71 |
| | Sftpb | 1.227 | 0.664 | 0.439 | 2.9E-74 | 4.4E-71 |
| | Retnla | 1.246 | 0.738 | 0.750 | 1.6E-71 | 2.3E-68 |
| | Nfib | 1.180 | 0.680 | 0.619 | 2.7E-70 | 4.1E-67 |
| | Aldh1a1 | 1.071 | 0.648 | 0.384 | 9.0E-69 | 1.3E-65 |
| | Nupr1 | 1.463 | 0.795 | 0.711 | 1.2E-68 | 1.8E-65 |
| | Ifitm3 | 1.335 | 0.809 | 0.700 | 6.7E-68 | 1.0E-64 |
| | Sdc4 | 1.174 | 0.760 | 0.674 | 1.7E-65 | 2.5E-62 |
| | Cldn3 | 1.119 | 0.776 | 0.778 | 5.3E-64 | 8.0E-61 |
| | Scd2 | 1.073 | 0.705 | 0.541 | 6.5E-63 | 9.7E-60 |
| | Timp3 | 1.525 | 0.657 | 0.569 | 1.1E-59 | 1.6E-56 |
| | Scgb3a2 | 2.769 | 0.668 | 0.618 | 4.3E-58 | 6.5E-55 |
| | Dbi | 1.376 | 0.776 | 0.646 | 7.1E-52 | 1.1E-48 |
| | Tmem176b | 1.986 | 0.660 | 0.567 | 2.6E-51 | 3.9E-48 |
| | Sdc1 | 1.649 | 0.700 | 0.709 | 1.3E-43 | 2.0E-40 |
| | Cyp2f2 | 2.315 | 0.663 | 0.704 | 2.5E-42 | 3.7E-39 |
| | Bgn | 1.569 | 0.550 | 0.243 | 1.1E-39 | 1.7E-36 |
| | Trf | 1.356 | 0.684 | 0.507 | 1.6E-37 | 2.5E-34 |
| | Wfdc18 | 1.473 | 0.361 | 0.563 | 6.2E-34 | 9.4E-31 |
| | Sparc | 2.443 | 0.542 | 0.358 | 1.0E-32 | 1.6E-29 |
| | Foxq1 | 1.159 | 0.583 | 0.537 | 2.1E-25 | 3.2E-22 |
| | Tff2 | 1.506 | 0.544 | 0.381 | 1.1E-24 | 1.7E-21 |
| | Ndufc2 | 1.065 | 0.660 | 0.625 | 1.1E-23 | 1.7E-20 |
| | Chchd10 | 1.241 | 0.644 | 0.644 | 2.8E-22 | 4.3E-19 |
| | Igfbp5 | 1.952 | 0.347 | 0.441 | 9.8E-22 | 1.5E-18 |
| | Ccnd1 | 1.370 | 0.572 | 0.514 | 2.1E-19 | 3.2E-16 |
| | Tagln2 | 1.084 | 0.659 | 0.593 | 5.6E-18 | 8.5E-15 |
| | Hmgn1 | 1.450 | 0.616 | 0.655 | 9.1E-17 | 1.4E-13 |
| | Snhg18 | 1.056 | 0.551 | 0.593 | 1.2E-10 | 1.7E-07 |
| | C3 | 1.081 | 0.651 | 0.554 | 3.2E-06 | 4.8E-03 |

| | | | | | | |
|---|---|---|---|---|---|---|
| | Tmem86a | 0.586 | 0.724 | 0.687 | 1.8E-78 | 2.7E-75 |
| | Osm | 0.502 | 0.286 | 0.341 | 5.9E-76 | 8.8E-73 |
| | C3 | 0.873 | 0.632 | 0.548 | 3.4E-72 | 5.1E-69 |
| | Slc7a11 | 1.622 | 0.565 | 0.458 | 4.8E-70 | 7.2E-67 |
| | Acod1 | 1.455 | 0.548 | 0.408 | 1.2E-65 | 1.8E-62 |
| | Ftl1-ps1 | 0.517 | 0.338 | 0.440 | 5.7E-65 | 8.5E-62 |
| | Naglu | 0.833 | 0.616 | 0.547 | 7.7E-61 | 1.2E-57 |
| | Gm5416 | 0.294 | 0.605 | 0.491 | 3.2E-55 | 4.8E-52 |
| | Prok2 | 0.362 | 0.643 | 0.575 | 3.5E-50 | 5.2E-47 |
| | Cd300c2 | 0.878 | 0.675 | 0.658 | 2.2E-49 | 3.3E-46 |
| | Hk2 | 0.718 | 0.466 | 0.600 | 6.9E-44 | 1.0E-40 |
| | Slc43a3 | 0.347 | 0.577 | 0.585 | 8.1E-44 | 1.2E-40 |
| | Creg1 | 1.161 | 0.702 | 0.668 | 6.8E-39 | 1.0E-35 |
| | Cldn1 | 0.605 | 0.582 | 0.623 | 1.3E-37 | 2.0E-34 |
| | Hspa1b | 0.760 | 0.659 | 0.609 | 2.5E-37 | 3.8E-34 |
| | G0s2 | 0.733 | 0.597 | 0.488 | 6.5E-37 | 9.7E-34 |
| | Tgm2 | 0.386 | 0.453 | 0.630 | 1.1E-35 | 1.7E-32 |
| | Ctsa | 0.424 | 0.748 | 0.700 | 1.5E-35 | 2.2E-32 |
| | Cxcr1 | 0.882 | 0.643 | 0.718 | 9.1E-35 | 1.4E-31 |
| | Gadd45g | 0.961 | 0.634 | 0.584 | 2.1E-31 | 3.2E-28 |
| | Cd68 | 0.694 | 0.714 | 0.715 | 6.3E-31 | 9.4E-28 |
| | Tnfrsf23 | 0.997 | 0.661 | 0.707 | 8.2E-31 | 1.2E-27 |
| | Hal | 0.570 | 0.514 | 0.710 | 1.2E-30 | 1.8E-27 |
| | Syngr1 | 0.408 | 0.721 | 0.777 | 1.4E-28 | 2.0E-25 |
| | Ptgs2 | 0.446 | 0.299 | 0.293 | 4.0E-27 | 6.0E-24 |
| | Slc37a2 | 0.659 | 0.609 | 0.589 | 5.6E-27 | 8.4E-24 |
| | Dpp7 | 0.410 | 0.624 | 0.628 | 5.7E-25 | 8.6E-22 |
| | Laptm5 | 0.357 | 0.748 | 0.739 | 3.4E-23 | 5.1E-20 |
| | Hpgds | 0.365 | 0.680 | 0.634 | 4.8E-21 | 7.2E-18 |
| | Rps6ka2 | 0.407 | 0.521 | 0.487 | 7.5E-20 | 1.1E-16 |
| | Slc6a6 | 0.352 | 0.448 | 0.520 | 6.6E-18 | 9.9E-15 |
| | Syne1 | 0.775 | 0.511 | 0.619 | 2.1E-17 | 3.1E-14 |
| | Vegfa | 0.957 | 0.495 | 0.433 | 7.9E-17 | 1.2E-13 |
| | Asprv1 | 0.304 | 0.639 | 0.584 | 1.2E-14 | 1.8E-11 |
| | Sqstm1 | 0.695 | 0.608 | 0.639 | 3.5E-14 | 5.2E-11 |
| | Tnf | 0.606 | 0.311 | 0.311 | 1.1E-12 | 1.6E-09 |
| | Gm26870 | 0.659 | 0.567 | 0.570 | 1.1E-11 | 1.7E-08 |
| | Hspa9 | 0.318 | 0.651 | 0.648 | 1.1E-09 | 1.6E-06 |
| | Osgin1 | 0.594 | 0.500 | 0.600 | 5.8E-08 | 8.7E-05 |
| | Csf2rb | 0.453 | 0.681 | 0.687 | 7.0E-07 | 1.1E-03 |
| | Egr1 | 0.431 | 0.574 | 0.554 | 8.2E-07 | 1.2E-03 |
| | Dhrs3 | 0.461 | 0.674 | 0.676 | 1.1E-06 | 1.7E-03 |
| | Thbs1 | 1.309 | 0.451 | 0.550 | 7.4E-06 | 1.1E-02 |
| | Cdkn1a | 0.339 | 0.776 | 0.770 | 4.9E-05 | 7.3E-02 |
| | Gadd45a | 0.424 | 0.639 | 0.622 | 1.8E-04 | 2.6E-01 |
| | Hist1h4i | 0.531 | 0.648 | 0.669 | 3.2E-04 | 4.8E-01 |
| | Slpi | 0.477 | 0.653 | 0.644 | 3.7E-04 | 5.5E-01 |
| | Ddit3 | 0.444 | 0.598 | 0.758 | 3.7E-04 | 5.6E-01 |
| | Il1f9 | 0.929 | 0.529 | 0.564 | 5.8E-04 | 8.7E-01 |
| | Sirpa | 0.299 | 0.560 | 0.634 | 5.2E-03 | 1.0E+00 |
| **T cells** | Cd3g | 2.602 | 0.918 | 0.436 | 0.0E+00 | 0.0E+00 |
| | Trbc2 | 2.530 | 0.904 | 0.229 | 0.0E+00 | 0.0E+00 |
| | Il7r | 2.308 | 0.820 | 0.669 | 0.0E+00 | 0.0E+00 |
| | Bcl2 | 2.093 | 0.817 | 0.414 | 0.0E+00 | 0.0E+00 |
| | Rpl3 | 1.972 | 0.988 | 0.699 | 0.0E+00 | 0.0E+00 |
| | Cd3e | 1.953 | 0.856 | 0.286 | 0.0E+00 | 0.0E+00 |
| | Cd3d | 1.904 | 0.890 | 0.375 | 0.0E+00 | 0.0E+00 |
| | Ptprcap | 1.902 | 0.828 | 0.511 | 0.0E+00 | 0.0E+00 |
| | Gimap1 | 1.875 | 0.851 | 0.436 | 0.0E+00 | 0.0E+00 |
| | Cxcr6 | 1.867 | 0.946 | 0.284 | 0.0E+00 | 0.0E+00 |
| | Ctla2a | 1.782 | 0.926 | 0.738 | 0.0E+00 | 0.0E+00 |
| | Trac | 1.722 | 0.781 | 0.126 | 0.0E+00 | 0.0E+00 |
| | Gimap3 | 1.701 | 0.790 | 0.197 | 0.0E+00 | 0.0E+00 |
| | AW112010 | 1.634 | 0.891 | 0.489 | 0.0E+00 | 0.0E+00 |
| | Ets1 | 1.568 | 0.849 | 0.434 | 0.0E+00 | 0.0E+00 |
| | Ms4a6b | 1.553 | 0.803 | 0.254 | 0.0E+00 | 0.0E+00 |
| | Skap1 | 1.513 | 0.806 | 0.163 | 0.0E+00 | 0.0E+00 |
| | Lck | 1.488 | 0.779 | 0.264 | 0.0E+00 | 0.0E+00 |
| | Ramp1 | 1.368 | 0.783 | 0.405 | 0.0E+00 | 0.0E+00 |
| | Rps2 | 1.301 | 0.993 | 0.889 | 0.0E+00 | 0.0E+00 |
| | Nkg7 | 1.264 | 0.876 | 0.521 | 0.0E+00 | 0.0E+00 |
| | Ptma | 1.233 | 0.983 | 0.718 | 0.0E+00 | 0.0E+00 |
| | Maf | 1.184 | 0.936 | 0.701 | 0.0E+00 | 0.0E+00 |
| | Thy1 | 1.172 | 0.878 | 0.165 | 0.0E+00 | 0.0E+00 |
| | Hspe1 | 1.096 | 0.879 | 0.665 | 0.0E+00 | 0.0E+00 |

| | | | | | | |
|---|---|---|---|---|---|---|
| | Tst | 1.319 | 0.571 | 0.602 | 1.0E-05 | 1.6E-02 |
| | Pmepa1 | 1.556 | 0.563 | 0.622 | 5.8E-05 | 8.7E-02 |
| | Fmo2 | 1.044 | 0.413 | 0.343 | 2.5E-04 | 3.8E-01 |
| | Crip2 | 1.269 | 0.502 | 0.560 | 1.6E-03 | 1.0E+00 |
| | Igfbp7 | 1.823 | 0.413 | 0.473 | 5.7E-03 | 1.0E+00 |
| | Sftpd | 1.945 | 0.489 | 0.461 | 8.2E-03 | 1.0E+00 |
| **T cells 2** | Ccl5 | 6.355 | 0.988 | 0.505 | 0.0E+00 | 0.0E+00 |
| | Nkg7 | 4.438 | 0.997 | 0.532 | 0.0E+00 | 0.0E+00 |
| | AW112010 | 3.298 | 0.997 | 0.502 | 0.0E+00 | 0.0E+00 |
| | Cd3g | 2.927 | 0.963 | 0.453 | 0.0E+00 | 0.0E+00 |
| | Klrd1 | 2.649 | 0.930 | 0.383 | 0.0E+00 | 0.0E+00 |
| | Gimap4 | 2.407 | 0.904 | 0.222 | 0.0E+00 | 0.0E+00 |
| | Cxcr6 | 2.039 | 0.977 | 0.308 | 0.0E+00 | 0.0E+00 |
| | Ikzf3 | 1.577 | 0.956 | 0.261 | 0.0E+00 | 0.0E+00 |
| | Pdcd1 | 1.413 | 0.989 | 0.234 | 0.0E+00 | 0.0E+00 |
| | Rgs16 | 0.578 | 0.966 | 0.464 | 0.0E+00 | 0.0E+00 |
| | Bcl2 | 2.127 | 0.963 | 0.426 | 1.1E-305 | 1.6E-302 |
| | Klre1 | 1.502 | 0.905 | 0.518 | 5.3E-294 | 8.0E-291 |
| | Gimap1 | 2.138 | 0.918 | 0.450 | 1.1E-276 | 1.6E-273 |
| | Ptprcap | 2.054 | 0.919 | 0.521 | 6.8E-265 | 1.0E-261 |
| | Gzma | 3.710 | 0.885 | 0.291 | 3.3E-263 | 5.0E-260 |
| | Serpinb9 | 1.165 | 0.887 | 0.348 | 3.6E-245 | 5.4E-242 |
| | Wls | 1.997 | 0.925 | 0.525 | 5.8E-242 | 8.8E-239 |
| | Ctsw | 2.064 | 0.826 | 0.206 | 9.3E-235 | 1.4E-231 |
| | Gpr171 | 0.922 | 0.902 | 0.568 | 2.0E-234 | 3.1E-231 |
| | Camk2n1 | 0.674 | 0.893 | 0.584 | 6.5E-232 | 9.8E-229 |
| | Klrb1c | 0.994 | 0.792 | 0.241 | 1.3E-217 | 2.0E-214 |
| | Thy1 | 1.399 | 0.805 | 0.194 | 2.8E-212 | 4.2E-209 |
| | Itga1 | 1.174 | 0.858 | 0.540 | 2.9E-207 | 4.3E-204 |
| | Lck | 1.936 | 0.843 | 0.282 | 9.4E-198 | 1.4E-194 |
| | Cd3e | 1.702 | 0.850 | 0.307 | 4.9E-196 | 7.4E-193 |
| | Bin1 | 1.028 | 0.889 | 0.559 | 4.2E-175 | 6.3E-172 |
| | Rpl3 | 1.327 | 0.992 | 0.710 | 1.8E-173 | 2.7E-170 |
| | mt-Nd1 | 1.057 | 0.991 | 0.669 | 1.2E-168 | 1.8E-165 |
| | Ctla2a | 1.995 | 0.849 | 0.746 | 4.1E-167 | 6.2E-164 |
| | Gbp4 | 0.491 | 0.908 | 0.599 | 6.1E-164 | 9.2E-161 |
| | Skap1 | 1.419 | 0.785 | 0.187 | 8.8E-160 | 1.3E-156 |
| | Lgals1 | 1.245 | 0.915 | 0.575 | 1.1E-159 | 1.6E-156 |
| | Gimap3 | 1.887 | 0.779 | 0.219 | 8.9E-157 | 1.3E-153 |
| | Gpr65 | 0.821 | 0.870 | 0.451 | 4.5E-153 | 6.7E-150 |
| | Ets1 | 1.314 | 0.838 | 0.449 | 9.4E-153 | 1.4E-149 |
| | Ptma | 1.057 | 0.983 | 0.728 | 1.0E-151 | 1.5E-148 |
| | Il2rb | 1.677 | 0.756 | 0.242 | 1.1E-147 | 1.7E-144 |
| | Crip1 | 0.729 | 0.947 | 0.503 | 4.0E-140 | 5.9E-137 |
| | Hspe1 | 0.981 | 0.928 | 0.671 | 4.3E-139 | 6.5E-136 |
| | Gzmb | 2.047 | 0.733 | 0.217 | 1.6E-136 | 2.4E-133 |
| | Bcl2a1d | 0.761 | 0.902 | 0.508 | 6.0E-129 | 9.1E-126 |
| | Ly9 | 0.743 | 0.798 | 0.321 | 9.2E-125 | 1.4E-121 |
| | Laptm5 | 0.968 | 0.994 | 0.735 | 1.4E-124 | 2.2E-121 |
| | Trbc2 | 2.552 | 0.763 | 0.257 | 3.1E-120 | 4.7E-117 |
| | Gimap6 | 1.119 | 0.771 | 0.323 | 5.1E-115 | 7.7E-112 |
| | Rps2 | 0.852 | 0.998 | 0.893 | 4.5E-114 | 6.7E-111 |
| | Trac | 1.458 | 0.734 | 0.151 | 8.0E-114 | 1.2E-110 |
| | AU020206 | 0.620 | 0.934 | 0.698 | 2.9E-110 | 4.4E-107 |
| | Irf8 | 0.532 | 0.853 | 0.635 | 3.3E-106 | 4.9E-103 |
| | H2afz | 0.948 | 0.944 | 0.804 | 2.5E-104 | 3.7E-101 |
| | Ccnd2 | 1.530 | 0.785 | 0.608 | 3.8E-104 | 5.7E-101 |
| | Dut | 0.533 | 0.774 | 0.448 | 1.5E-98 | 2.3E-95 |
| | Emp3 | 0.746 | 0.876 | 0.460 | 5.1E-97 | 7.6E-94 |
| | Id2 | 0.628 | 0.977 | 0.778 | 1.4E-96 | 2.0E-93 |
| | Ebpl | 0.549 | 0.782 | 0.407 | 2.9E-88 | 4.4E-85 |
| | Ndufa4 | 0.747 | 0.879 | 0.668 | 4.0E-88 | 6.0E-85 |
| | Cd2 | 1.412 | 0.725 | 0.421 | 1.2E-85 | 1.8E-82 |
| | Ifi47 | 0.906 | 0.766 | 0.648 | 1.7E-82 | 2.6E-79 |
| | Klrk1 | 1.640 | 0.666 | 0.100 | 2.0E-81 | 3.0E-78 |
| | Serpinb6b | 0.741 | 0.769 | 0.688 | 8.2E-73 | 1.2E-69 |
| | Cx3cr1 | 0.855 | 0.716 | 0.365 | 3.6E-72 | 5.4E-69 |
| | Reep5 | 0.686 | 0.904 | 0.713 | 1.6E-71 | 2.4E-68 |
| | Pla2g16 | 0.823 | 0.780 | 0.548 | 3.0E-65 | 4.5E-62 |
| | Racgap1 | 0.500 | 0.689 | 0.255 | 1.8E-63 | 2.7E-60 |
| | Cd7 | 1.311 | 0.653 | 0.123 | 3.7E-59 | 5.5E-56 |
| | F2r | 1.007 | 0.699 | 0.655 | 1.7E-57 | 2.5E-54 |
| | Itm2c | 0.598 | 0.789 | 0.595 | 2.3E-57 | 3.5E-54 |
| | Maf | 0.484 | 0.773 | 0.713 | 6.1E-56 | 9.2E-53 |
| | Sh2d1a | 1.028 | 0.637 | 0.160 | 4.7E-49 | 7.0E-46 |

| | | | | | | |
|---|---|---|---|---|---|---|
| | Ccl5 | 1.061 | 0.893 | 0.492 | 0.0E+00 | 0.0E+00 |
| | Pdcd1 | 1.058 | 0.965 | 0.206 | 0.0E+00 | 0.0E+00 |
| | Icos | 1.040 | 0.721 | 0.224 | 0.0E+00 | 0.0E+00 |
| | Il2rb | 0.938 | 0.822 | 0.219 | 0.0E+00 | 0.0E+00 |
| | Trdc | 0.926 | 0.888 | 0.260 | 0.0E+00 | 0.0E+00 |
| | Ikzf3 | 0.876 | 0.895 | 0.238 | 0.0E+00 | 0.0E+00 |
| | F2r | 0.833 | 0.807 | 0.646 | 0.0E+00 | 0.0E+00 |
| | mt-Nd1 | 0.778 | 0.971 | 0.657 | 0.0E+00 | 0.0E+00 |
| | Tcrg-C1 | 0.619 | 0.895 | 0.175 | 0.0E+00 | 0.0E+00 |
| | Cd163l1 | 0.484 | 0.811 | 0.167 | 0.0E+00 | 0.0E+00 |
| | Camk2n1 | 0.476 | 0.974 | 0.567 | 0.0E+00 | 0.0E+00 |
| | Il2ra | 0.462 | 0.861 | 0.138 | 0.0E+00 | 0.0E+00 |
| | Gbp4 | 0.441 | 0.879 | 0.589 | 0.0E+00 | 0.0E+00 |
| | AU020206 | 0.440 | 0.957 | 0.687 | 0.0E+00 | 0.0E+00 |
| | Nrp1 | 0.439 | 0.886 | 0.725 | 0.0E+00 | 0.0E+00 |
| | Gpr171 | 0.417 | 0.782 | 0.562 | 3.2E-288 | 4.8E-285 |
| | S100a4 | 1.528 | 0.865 | 0.488 | 9.4E-267 | 1.4E-263 |
| | Crip1 | 0.519 | 0.881 | 0.490 | 1.0E-230 | 1.6E-227 |
| | Ctsw | 0.610 | 0.694 | 0.191 | 3.5E-222 | 5.3E-219 |
| | Slc25a4 | 0.912 | 0.791 | 0.526 | 7.1E-222 | 1.1E-218 |
| | Lat | 0.926 | 0.715 | 0.457 | 1.0E-202 | 1.6E-199 |
| | Gimap4 | 1.294 | 0.695 | 0.208 | 3.3E-200 | 5.0E-197 |
| | Tcf7 | 0.897 | 0.659 | 0.050 | 2.1E-199 | 3.1E-196 |
| | Gimap6 | 0.852 | 0.713 | 0.309 | 4.4E-194 | 6.6E-191 |
| | Cd28 | 1.003 | 0.702 | 0.529 | 1.9E-184 | 2.8E-181 |
| | Bcl11b | 1.081 | 0.686 | 0.332 | 1.0E-181 | 1.6E-178 |
| | Ebpl | 0.434 | 0.744 | 0.395 | 1.2E-177 | 1.7E-174 |
| | Itgb7 | 1.071 | 0.709 | 0.520 | 5.7E-170 | 8.5E-167 |
| | Wls | 0.694 | 0.775 | 0.518 | 1.0E-164 | 1.6E-161 |
| | Cd2 | 1.084 | 0.694 | 0.411 | 6.8E-162 | 1.0E-158 |
| | Gpr65 | 0.404 | 0.737 | 0.442 | 2.0E-150 | 3.0E-147 |
| | 4930523C07Rik | 0.862 | 0.693 | 0.505 | 1.3E-137 | 1.9E-134 |
| | Pla2g16 | 0.610 | 0.767 | 0.540 | 8.1E-130 | 1.2E-126 |
| | Pgls | 0.429 | 0.820 | 0.627 | 5.4E-121 | 8.1E-118 |
| | Dut | 0.633 | 0.647 | 0.443 | 7.5E-88 | 1.1E-84 |
| | Cd7 | 0.448 | 0.374 | 0.119 | 7.0E-70 | 1.1E-66 |
| | Ifi203 | 0.760 | 0.642 | 0.560 | 2.2E-69 | 3.3E-66 |
| | Il18r1 | 0.468 | 0.359 | 0.433 | 9.1E-69 | 1.4E-65 |
| | Ranbp1 | 0.564 | 0.715 | 0.643 | 6.1E-68 | 9.1E-65 |
| | Mllt3 | 0.629 | 0.666 | 0.631 | 2.6E-67 | 3.8E-64 |
| | Ar | 0.464 | 0.380 | 0.208 | 8.1E-65 | 1.2E-61 |
| | Ccnd2 | 0.721 | 0.677 | 0.608 | 5.6E-64 | 8.3E-61 |
| | Tmem176a | 0.693 | 0.737 | 0.580 | 1.5E-62 | 2.3E-59 |
| | Igfbp4 | 0.442 | 0.371 | 0.530 | 4.0E-61 | 6.0E-58 |
| | Hmgn1 | 0.444 | 0.702 | 0.651 | 8.9E-57 | 1.3E-53 |
| | Nucks1 | 0.698 | 0.651 | 0.530 | 2.2E-56 | 3.2E-53 |
| | Ifi27l2a | 0.810 | 0.649 | 0.527 | 1.1E-55 | 1.7E-52 |
| | Nrip1 | 0.824 | 0.664 | 0.612 | 7.0E-54 | 1.1E-50 |
| | Esyt1 | 0.550 | 0.636 | 0.410 | 6.1E-53 | 9.1E-50 |
| | Ifi47 | 0.471 | 0.662 | 0.649 | 1.9E-50 | 2.9E-47 |
| | Dock10 | 0.556 | 0.688 | 0.678 | 8.2E-47 | 1.2E-43 |
| | Trbc1 | 1.908 | 0.580 | 0.202 | 3.3E-44 | 4.9E-41 |
| | C1qbp | 0.468 | 0.666 | 0.664 | 9.5E-43 | 1.4E-39 |
| | Ms4a4b | 0.952 | 0.551 | 0.088 | 2.1E-41 | 3.2E-38 |
| | Gm12840 | 1.196 | 0.572 | 0.386 | 9.3E-39 | 1.4E-35 |
| | Gimap7 | 0.743 | 0.566 | 0.222 | 1.6E-32 | 2.4E-29 |
| | Klk8 | 0.546 | 0.598 | 0.493 | 8.8E-32 | 1.3E-28 |
| | Rexo2 | 0.515 | 0.636 | 0.637 | 1.1E-25 | 1.6E-22 |
| | Zbtb20 | 0.641 | 0.604 | 0.540 | 1.0E-24 | 1.5E-21 |
| | Evl | 0.589 | 0.582 | 0.337 | 5.4E-18 | 8.2E-15 |
| | Las1l | 0.388 | 0.576 | 0.490 | 5.6E-17 | 8.4E-14 |
| | Kcnq1ot1 | 0.582 | 0.619 | 0.658 | 8.6E-16 | 1.3E-12 |
| | S1pr1 | 0.739 | 0.552 | 0.507 | 8.8E-16 | 1.3E-12 |
| | Lef1 | 0.517 | 0.517 | 0.035 | 3.6E-14 | 5.4E-11 |
| | Gpr183 | 0.598 | 0.436 | 0.265 | 9.7E-11 | 1.5E-07 |
| | Kcnn4 | 0.510 | 0.468 | 0.579 | 5.0E-10 | 7.6E-07 |
| | Gm8369 | 0.566 | 0.443 | 0.091 | 1.8E-09 | 2.8E-06 |
| | Tnfsf8 | 0.669 | 0.451 | 0.396 | 5.5E-09 | 8.3E-06 |
| | Ramp3 | 0.410 | 0.540 | 0.413 | 1.2E-07 | 1.8E-04 |
| | Ccr2 | 0.445 | 0.449 | 0.261 | 4.5E-07 | 6.8E-04 |
| | Tmem160 | 0.367 | 0.600 | 0.584 | 5.6E-07 | 8.5E-04 |
| | Dapl1 | 0.839 | 0.526 | 0.394 | 4.6E-06 | 6.9E-03 |
| | Cblb | 0.388 | 0.487 | 0.637 | 1.4E-04 | 2.1E-01 |
| | Odc1 | 0.426 | 0.573 | 0.569 | 1.2E-03 | 1.0E+00 |
| | Tmem176b | 0.675 | 0.581 | 0.569 | 5.8E-03 | 1.0E+00 |

| | | | | | | |
|---|---|---|---|---|---|---|
| | Cd3d | 1.570 | 0.660 | 0.399 | 1.5E-48 | 2.3E-45 |
| | Nme1 | 0.638 | 0.798 | 0.715 | 6.0E-44 | 9.0E-41 |
| | Cd48 | 1.146 | 0.658 | 0.445 | 9.0E-42 | 1.3E-38 |
| | Pycard | 0.597 | 0.864 | 0.778 | 3.7E-39 | 5.5E-36 |
| | Zbp1 | 0.508 | 0.708 | 0.633 | 3.6E-36 | 5.5E-33 |
| | Itgb7 | 0.911 | 0.655 | 0.528 | 4.8E-34 | 7.2E-31 |
| | Gimap7 | 1.216 | 0.618 | 0.234 | 2.7E-31 | 4.0E-28 |
| | Tmem160 | 0.727 | 0.687 | 0.583 | 2.4E-28 | 3.7E-25 |
| | Phf11b | 0.550 | 0.623 | 0.316 | 8.7E-28 | 1.3E-24 |
| | Dock10 | 0.734 | 0.696 | 0.678 | 3.1E-27 | 4.7E-24 |
| | Klra4 | 1.070 | 0.347 | 0.141 | 2.8E-26 | 4.2E-23 |
| | Kcnq1ot1 | 0.612 | 0.692 | 0.655 | 1.8E-18 | 2.6E-15 |
| | Nucks1 | 0.618 | 0.643 | 0.535 | 1.4E-17 | 2.1E-14 |
| | Ranbp1 | 0.579 | 0.682 | 0.647 | 3.9E-15 | 5.9E-12 |
| | Trbc1 | 1.609 | 0.582 | 0.216 | 9.1E-15 | 1.4E-11 |
| | Osbpl3 | 0.642 | 0.631 | 0.611 | 1.3E-13 | 1.9E-10 |
| | AC149090.1 | 0.829 | 0.602 | 0.458 | 1.3E-12 | 2.0E-09 |
| | Bcl11b | 1.186 | 0.573 | 0.347 | 3.4E-12 | 5.1E-09 |
| | Sept9 | 0.475 | 0.618 | 0.475 | 1.9E-10 | 2.8E-07 |
| | Il18r1 | 0.691 | 0.586 | 0.426 | 6.5E-10 | 9.8E-07 |
| | Il7r | 0.709 | 0.631 | 0.679 | 2.1E-09 | 3.1E-06 |
| | S100a4 | 1.032 | 0.598 | 0.508 | 3.6E-09 | 5.4E-06 |
| | Evl | 0.553 | 0.597 | 0.346 | 4.4E-08 | 6.6E-05 |
| | Dok2 | 0.675 | 0.563 | 0.438 | 3.4E-07 | 5.0E-04 |
| | Sms | 0.505 | 0.435 | 0.441 | 5.8E-07 | 8.7E-04 |
| | Ms4a6b | 0.880 | 0.574 | 0.280 | 7.1E-07 | 1.1E-03 |
| | Ybx3 | 0.569 | 0.449 | 0.607 | 4.5E-06 | 6.7E-03 |
| | Cblb | 0.518 | 0.449 | 0.632 | 7.0E-06 | 1.0E-02 |
| | Esyt1 | 0.697 | 0.566 | 0.420 | 1.5E-05 | 2.2E-02 |
| | Sept11 | 0.805 | 0.568 | 0.594 | 8.0E-05 | 1.2E-01 |
| | Slamf7 | 0.845 | 0.534 | 0.547 | 8.7E-03 | 1.0E+00 |
| **Prol.** | Pclaf | 2.491 | 0.958 | 0.145 | 0.0E+00 | 0.0E+00 |
| | Tpx2 | 1.659 | 0.979 | 0.461 | 0.0E+00 | 0.0E+00 |
| | Hmmr | 1.416 | 0.965 | 0.417 | 0.0E+00 | 0.0E+00 |
| | Cks1b | 1.929 | 0.966 | 0.434 | 5.1E-307 | 7.7E-304 |
| | Ccnb1 | 1.412 | 0.897 | 0.279 | 9.5E-295 | 1.4E-291 |
| | Spc25 | 1.049 | 0.954 | 0.304 | 7.3E-293 | 1.1E-289 |
| | Stmn1 | 2.414 | 0.965 | 0.511 | 2.1E-290 | 3.1E-287 |
| | Mki67 | 2.558 | 0.959 | 0.570 | 2.8E-289 | 4.3E-286 |
| | Nusap1 | 1.629 | 0.920 | 0.239 | 1.2E-271 | 1.7E-268 |
| | Cdca8 | 1.719 | 0.927 | 0.543 | 1.2E-265 | 1.8E-262 |
| | Tuba1b | 2.789 | 0.975 | 0.597 | 1.6E-247 | 2.4E-244 |
| | Cenpe | 1.483 | 0.904 | 0.409 | 3.6E-245 | 5.4E-242 |
| | Racgap1 | 1.111 | 0.892 | 0.252 | 2.0E-234 | 3.0E-231 |
| | Ptma | 2.106 | 0.961 | 0.729 | 9.1E-234 | 1.4E-230 |
| | Lig1 | 1.309 | 0.922 | 0.604 | 1.5E-224 | 2.2E-221 |
| | Comt | 1.054 | 0.981 | 0.624 | 1.5E-214 | 2.2E-211 |
| | Fn1 | 1.115 | 0.950 | 0.408 | 2.4E-214 | 3.6E-211 |
| | Mt1 | 1.467 | 0.989 | 0.771 | 5.7E-214 | 8.6E-211 |
| | Tacc3 | 1.088 | 0.885 | 0.465 | 6.6E-212 | 1.0E-208 |
| | Nucks1 | 2.033 | 0.961 | 0.529 | 6.2E-211 | 9.3E-208 |
| | H2afz | 1.932 | 0.933 | 0.805 | 2.0E-204 | 3.0E-201 |
| | Tagln2 | 2.069 | 0.991 | 0.587 | 7.3E-196 | 1.1E-192 |
| | Snx5 | 1.077 | 0.986 | 0.545 | 5.5E-194 | 8.2E-191 |
| | Tubb5 | 2.638 | 0.927 | 0.597 | 1.1E-193 | 1.6E-190 |
| | Hmgn1 | 1.810 | 0.970 | 0.649 | 2.7E-191 | 4.0E-188 |
| | Atad2 | 1.082 | 0.929 | 0.479 | 9.4E-191 | 1.4E-187 |
| | Ppp1r14b | 1.888 | 0.981 | 0.660 | 1.3E-189 | 1.9E-186 |
| | Lgals1 | 1.754 | 0.982 | 0.574 | 1.6E-188 | 2.4E-185 |
| | Slc25a4 | 1.447 | 0.961 | 0.533 | 1.2E-185 | 1.8E-182 |
| | Ccnb2 | 1.586 | 0.846 | 0.228 | 4.4E-185 | 6.7E-182 |
| | Hebp1 | 1.259 | 0.989 | 0.644 | 2.7E-183 | 4.1E-180 |
| | Ezh2 | 1.288 | 0.869 | 0.522 | 1.3E-180 | 2.0E-177 |
| | Crip1 | 1.624 | 0.982 | 0.504 | 1.2E-176 | 1.8E-173 |
| | Tmem256 | 1.475 | 0.968 | 0.545 | 1.8E-173 | 2.7E-170 |
| | Prdx1 | 1.850 | 0.996 | 0.665 | 5.2E-173 | 7.7E-170 |
| | Selenoh | 1.849 | 0.871 | 0.469 | 1.9E-168 | 2.8E-165 |
| | Cenpa | 1.756 | 0.867 | 0.426 | 3.0E-166 | 4.4E-163 |
| | Cenpx | 1.309 | 0.959 | 0.664 | 6.9E-165 | 1.0E-161 |
| | Rexo2 | 1.263 | 0.965 | 0.631 | 8.8E-164 | 1.3E-160 |
| | Ndufa4 | 2.100 | 0.982 | 0.667 | 3.8E-158 | 5.8E-155 |
| | Pycard | 1.529 | 0.989 | 0.776 | 2.6E-157 | 4.0E-154 |
| | Smc4 | 1.764 | 0.924 | 0.614 | 5.8E-155 | 8.7E-152 |
| | Fabp5 | 1.449 | 0.993 | 0.803 | 1.1E-154 | 1.7E-151 |
| | Cks2 | 1.430 | 0.931 | 0.464 | 1.4E-152 | 2.2E-149 |

| | | | | | | |
|---|---|---|---|---|---|---|
| | Igfbp4 | 0.442 | 0.371 | 0.530 | 4.0E-61 | 6.0E-58 |
| | Hmgn1 | 0.444 | 0.702 | 0.651 | 8.9E-57 | 1.3E-53 |
| | Nucks1 | 0.698 | 0.651 | 0.530 | 2.2E-56 | 3.2E-53 |
| | Ifi27l2a | 0.810 | 0.649 | 0.527 | 1.1E-55 | 1.7E-52 |
| | Nrip1 | 0.824 | 0.664 | 0.612 | 7.0E-54 | 1.1E-50 |
| | Esyt1 | 0.550 | 0.636 | 0.410 | 6.1E-53 | 9.1E-50 |
| | Ifi47 | 0.471 | 0.662 | 0.649 | 1.9E-50 | 2.9E-47 |
| | Dock10 | 0.556 | 0.688 | 0.678 | 8.2E-47 | 1.2E-43 |
| | Trbc1 | 1.908 | 0.580 | 0.202 | 3.3E-44 | 4.9E-41 |
| | C1qbp | 0.468 | 0.666 | 0.664 | 9.5E-43 | 1.4E-39 |
| | Ms4a4b | 0.952 | 0.551 | 0.088 | 2.1E-41 | 3.2E-38 |
| | Gm12840 | 1.196 | 0.572 | 0.386 | 9.3E-39 | 1.4E-35 |
| | Gimap7 | 0.743 | 0.566 | 0.222 | 1.6E-32 | 2.4E-29 |
| | Klk8 | 0.546 | 0.598 | 0.493 | 8.8E-32 | 1.3E-28 |
| | Rexo2 | 0.515 | 0.636 | 0.637 | 1.1E-25 | 1.6E-22 |
| | Zbtb20 | 0.641 | 0.604 | 0.540 | 1.0E-24 | 1.5E-21 |
| | Evl | 0.589 | 0.582 | 0.337 | 5.4E-18 | 8.2E-15 |
| | Las1l | 0.388 | 0.576 | 0.490 | 5.6E-17 | 8.4E-14 |
| | Kcnq1ot1 | 0.582 | 0.619 | 0.658 | 8.6E-16 | 1.3E-12 |
| | S1pr1 | 0.739 | 0.552 | 0.507 | 8.8E-16 | 1.3E-12 |
| | Lef1 | 0.517 | 0.517 | 0.035 | 3.6E-14 | 5.4E-11 |
| | Gpr183 | 0.598 | 0.436 | 0.265 | 9.7E-11 | 1.5E-07 |
| | Kcnn4 | 0.510 | 0.468 | 0.579 | 5.0E-10 | 7.6E-07 |
| | Gm8369 | 0.566 | 0.443 | 0.091 | 1.8E-09 | 2.8E-06 |
| | Tnfsf8 | 0.669 | 0.451 | 0.396 | 5.5E-09 | 8.3E-06 |
| | Ramp3 | 0.410 | 0.540 | 0.413 | 1.2E-07 | 1.8E-04 |
| | Ccr2 | 0.445 | 0.449 | 0.261 | 4.5E-07 | 6.8E-04 |
| | Tmem160 | 0.367 | 0.600 | 0.584 | 5.6E-07 | 8.5E-04 |
| | Dapl1 | 0.839 | 0.526 | 0.394 | 4.6E-06 | 6.9E-03 |
| | Cblb | 0.388 | 0.487 | 0.637 | 1.4E-04 | 2.1E-01 |
| | Odc1 | 0.426 | 0.573 | 0.569 | 1.2E-03 | 1.0E+00 |
| | Tmem176b | 0.675 | 0.581 | 0.569 | 5.8E-03 | 1.0E+00 |
| **Mac 1** | Apoe | 3.645 | 0.999 | 0.776 | 0.0E+00 | 0.0E+00 |
| | Ctss | 3.156 | 1.000 | 0.646 | 0.0E+00 | 0.0E+00 |
| | Fn1 | 3.070 | 0.961 | 0.389 | 0.0E+00 | 0.0E+00 |
| | Lyz2 | 2.912 | 1.000 | 0.750 | 0.0E+00 | 0.0E+00 |
| | Trem2 | 2.398 | 0.999 | 0.563 | 0.0E+00 | 0.0E+00 |
| | Lgals1 | 2.386 | 1.000 | 0.559 | 0.0E+00 | 0.0E+00 |
| | Spp1 | 2.356 | 0.996 | 0.780 | 0.0E+00 | 0.0E+00 |
| | Mafb | 2.251 | 0.977 | 0.576 | 0.0E+00 | 0.0E+00 |
| | C1qa | 2.202 | 0.980 | 0.573 | 0.0E+00 | 0.0E+00 |
| | Fabp5 | 2.165 | 0.994 | 0.796 | 0.0E+00 | 0.0E+00 |
| | C1qb | 2.126 | 0.982 | 0.493 | 0.0E+00 | 0.0E+00 |
| | Psap | 2.095 | 1.000 | 0.747 | 0.0E+00 | 0.0E+00 |
| | Ctsl | 2.035 | 0.999 | 0.840 | 0.0E+00 | 0.0E+00 |
| | C1qc | 2.002 | 0.983 | 0.633 | 0.0E+00 | 0.0E+00 |
| | Gpnmb | 1.975 | 0.994 | 0.642 | 0.0E+00 | 0.0E+00 |
| | Vim | 1.894 | 0.998 | 0.631 | 0.0E+00 | 0.0E+00 |
| | Ms4a6c | 1.887 | 0.981 | 0.393 | 0.0E+00 | 0.0E+00 |
| | Emp3 | 1.866 | 0.993 | 0.440 | 0.0E+00 | 0.0E+00 |
| | Plin2 | 1.859 | 1.000 | 0.843 | 0.0E+00 | 0.0E+00 |
| | Cd68 | 1.847 | 0.999 | 0.699 | 0.0E+00 | 0.0E+00 |
| | F13a1 | 1.817 | 0.892 | 0.611 | 0.0E+00 | 0.0E+00 |
| | Lgmn | 1.795 | 0.994 | 0.615 | 0.0E+00 | 0.0E+00 |
| | Grn | 1.771 | 1.000 | 0.840 | 0.0E+00 | 0.0E+00 |
| | S100a4 | 1.755 | 0.940 | 0.486 | 0.0E+00 | 0.0E+00 |
| | Lrp1 | 1.691 | 0.995 | 0.503 | 0.0E+00 | 0.0E+00 |
| | Ms4a7 | 1.620 | 0.981 | 0.421 | 0.0E+00 | 0.0E+00 |
| | mt-Nd1 | 1.606 | 0.996 | 0.658 | 0.0E+00 | 0.0E+00 |
| | Hexa | 1.594 | 0.999 | 0.688 | 0.0E+00 | 0.0E+00 |
| | Npc2 | 1.576 | 1.000 | 0.885 | 0.0E+00 | 0.0E+00 |
| | Hexb | 1.575 | 0.998 | 0.741 | 0.0E+00 | 0.0E+00 |
| | Crip1 | 1.546 | 0.994 | 0.486 | 0.0E+00 | 0.0E+00 |
| | Lamp1 | 1.530 | 1.000 | 0.856 | 0.0E+00 | 0.0E+00 |
| | Saa3 | 1.523 | 0.868 | 0.613 | 0.0E+00 | 0.0E+00 |
| | Mgst1 | 1.502 | 0.992 | 0.589 | 0.0E+00 | 0.0E+00 |
| | Syngr1 | 1.501 | 0.994 | 0.759 | 0.0E+00 | 0.0E+00 |
| | Ctsz | 1.495 | 1.000 | 0.786 | 0.0E+00 | 0.0E+00 |
| | Ctsb | 1.482 | 1.000 | 0.942 | 0.0E+00 | 0.0E+00 |
| | Ccl9 | 1.476 | 0.919 | 0.528 | 0.0E+00 | 0.0E+00 |
| | C3ar1 | 1.475 | 0.992 | 0.552 | 0.0E+00 | 0.0E+00 |
| | Pycard | 1.449 | 0.984 | 0.769 | 0.0E+00 | 0.0E+00 |
| | Ctsd | 1.430 | 1.000 | 0.964 | 0.0E+00 | 0.0E+00 |
| | Smpdl3a | 1.420 | 0.995 | 0.611 | 0.0E+00 | 0.0E+00 |
| | Anxa5 | 1.419 | 0.999 | 0.669 | 0.0E+00 | 0.0E+00 |

| | | | | | | |
|---|---|---|---|---|---|---|
| | Racgap1 | 1.111 | 0.892 | 0.252 | 2.0E-234 | 3.0E-231 |
| | Ptma | 2.106 | 0.961 | 0.729 | 9.1E-234 | 1.4E-230 |
| | Lig1 | 1.309 | 0.922 | 0.604 | 1.5E-224 | 2.2E-221 |
| | Comt | 1.054 | 0.981 | 0.624 | 1.5E-214 | 2.2E-211 |
| | Fn1 | 1.115 | 0.950 | 0.408 | 2.4E-214 | 3.6E-211 |
| | Mt1 | 1.467 | 0.989 | 0.771 | 5.7E-214 | 8.6E-211 |
| | Tacc3 | 1.088 | 0.885 | 0.465 | 6.6E-212 | 1.0E-208 |
| | Nucks1 | 2.033 | 0.961 | 0.529 | 6.2E-211 | 9.3E-208 |
| | H2afz | 1.932 | 0.933 | 0.805 | 2.0E-204 | 3.0E-201 |
| | Tagln2 | 2.069 | 0.991 | 0.587 | 7.3E-196 | 1.1E-192 |
| | Snx5 | 1.077 | 0.986 | 0.545 | 5.5E-194 | 8.2E-191 |
| | Tubb5 | 2.638 | 0.927 | 0.597 | 1.1E-193 | 1.6E-190 |
| | Hmgn1 | 1.810 | 0.970 | 0.649 | 2.7E-191 | 4.0E-188 |
| | Atad2 | 1.082 | 0.929 | 0.479 | 9.4E-191 | 1.4E-187 |
| | Ppp1r14b | 1.888 | 0.981 | 0.660 | 1.3E-189 | 1.9E-186 |
| | Lgals1 | 1.754 | 0.982 | 0.574 | 1.6E-188 | 2.4E-185 |
| | Slc25a4 | 1.447 | 0.961 | 0.533 | 1.2E-185 | 1.8E-182 |
| | Ccnb2 | 1.586 | 0.846 | 0.228 | 4.4E-185 | 6.7E-182 |
| | Hebp1 | 1.259 | 0.989 | 0.644 | 2.7E-183 | 4.1E-180 |
| | Ezh2 | 1.288 | 0.869 | 0.522 | 1.3E-180 | 2.0E-177 |
| | Crip1 | 1.624 | 0.982 | 0.504 | 1.2E-176 | 1.8E-173 |
| | Tmem256 | 1.475 | 0.968 | 0.545 | 1.8E-173 | 2.7E-170 |
| | Prdx1 | 1.850 | 0.996 | 0.665 | 5.2E-173 | 7.7E-170 |
| | Selenoh | 1.849 | 0.871 | 0.469 | 1.9E-168 | 2.8E-165 |
| | Cenpa | 1.756 | 0.867 | 0.426 | 3.0E-166 | 4.4E-163 |
| | Cenpx | 1.309 | 0.959 | 0.664 | 6.9E-165 | 1.0E-161 |
| | Rexo2 | 1.263 | 0.965 | 0.631 | 8.8E-164 | 1.3E-160 |
| | Ndufa4 | 2.100 | 0.982 | 0.667 | 3.8E-158 | 5.8E-155 |
| | Pycard | 1.529 | 0.989 | 0.776 | 2.6E-157 | 4.0E-154 |
| | Smc4 | 1.764 | 0.924 | 0.614 | 5.8E-155 | 8.7E-152 |
| | Fabp5 | 1.449 | 0.993 | 0.803 | 1.1E-154 | 1.7E-151 |
| | Cks2 | 1.430 | 0.931 | 0.464 | 1.4E-152 | 2.2E-149 |
| | Nap1l1 | 1.688 | 0.954 | 0.558 | 1.7E-148 | 2.5E-145 |
| | Hspa9 | 1.285 | 0.938 | 0.643 | 9.9E-148 | 1.5E-144 |
| | Nme1 | 1.906 | 0.965 | 0.712 | 1.0E-146 | 1.5E-143 |
| | Prdx2 | 1.788 | 0.940 | 0.640 | 2.6E-141 | 3.9E-138 |
| | Rpl3 | 1.645 | 0.952 | 0.712 | 2.0E-140 | 3.0E-137 |
| | Rps2 | 1.458 | 0.940 | 0.894 | 2.3E-130 | 3.5E-127 |
| | Spp1 | 1.101 | 0.991 | 0.788 | 3.2E-128 | 4.8E-125 |
| | Tubb4b | 1.816 | 0.977 | 0.715 | 3.8E-128 | 5.6E-125 |
| | Cybb | 1.037 | 0.947 | 0.519 | 8.9E-128 | 1.3E-124 |
| | Vim | 1.633 | 0.988 | 0.644 | 2.4E-125 | 3.5E-122 |
| | Mcm7 | 1.071 | 0.851 | 0.556 | 1.5E-124 | 2.2E-121 |
| | Ybx3 | 1.869 | 0.865 | 0.599 | 1.9E-123 | 2.8E-120 |
| | Siva1 | 1.363 | 0.931 | 0.730 | 6.4E-120 | 9.6E-117 |
| | Pgls | 1.338 | 0.950 | 0.632 | 5.3E-119 | 8.0E-116 |
| | Ptms | 1.038 | 0.988 | 0.620 | 2.7E-116 | 4.1E-113 |
| | Fkbp2 | 1.125 | 0.931 | 0.533 | 4.2E-114 | 6.3E-111 |
| | Tyms | 1.094 | 0.796 | 0.470 | 1.6E-111 | 2.5E-108 |
| | Lamtor4 | 1.288 | 0.982 | 0.702 | 9.9E-107 | 1.5E-103 |
| | Hist1h4d | 1.284 | 0.839 | 0.676 | 1.0E-105 | 1.5E-102 |
| | Dbi | 1.394 | 0.936 | 0.644 | 2.0E-103 | 3.0E-100 |
| | Top2a | 2.140 | 0.749 | 0.304 | 1.5E-100 | 2.3E-97 |
| | mt-Nd1 | 1.535 | 0.881 | 0.672 | 2.2E-98 | 3.2E-95 |
| | Birc5 | 2.119 | 0.756 | 0.487 | 3.7E-96 | 5.6E-93 |
| | Hsp90b1 | 1.288 | 0.991 | 0.786 | 4.3E-96 | 6.5E-93 |
| | Cdca3 | 1.395 | 0.733 | 0.222 | 6.4E-91 | 9.6E-88 |
| | Reep5 | 1.089 | 0.966 | 0.713 | 1.3E-90 | 2.0E-87 |
| | Gm26917 | 1.427 | 0.839 | 0.566 | 1.8E-88 | 2.7E-85 |
| | Aprt | 1.160 | 0.943 | 0.737 | 1.0E-83 | 1.6E-80 |
| | Gzma | 1.044 | 0.791 | 0.294 | 5.1E-75 | 7.7E-72 |
| | C1qbp | 1.452 | 0.809 | 0.661 | 1.8E-71 | 2.7E-68 |
| | Ranbp1 | 1.666 | 0.798 | 0.645 | 3.8E-71 | 5.8E-68 |
| | Cdc20 | 1.193 | 0.713 | 0.484 | 2.7E-70 | 4.1E-67 |
| | Tmed3 | 1.101 | 0.873 | 0.617 | 4.5E-69 | 6.7E-66 |
| | Pdia6 | 1.130 | 0.973 | 0.783 | 2.8E-67 | 4.2E-64 |
| | Cenpf | 1.498 | 0.697 | 0.197 | 9.5E-65 | 1.4E-61 |
| | Calr | 1.114 | 0.973 | 0.770 | 1.3E-59 | 1.9E-56 |
| | Ramp1 | 1.043 | 0.756 | 0.421 | 6.4E-56 | 9.5E-53 |
| | Atp5g1 | 1.539 | 0.749 | 0.648 | 3.7E-50 | 5.5E-47 |
| | Hspd1 | 1.695 | 0.731 | 0.563 | 1.7E-49 | 2.6E-46 |
| | Igkc | 2.677 | 0.216 | 0.301 | 7.5E-49 | 1.1E-45 |
| | Igha | 1.754 | 0.694 | 0.385 | 6.0E-45 | 9.0E-42 |
| | Hspe1 | 1.292 | 0.779 | 0.675 | 1.7E-44 | 2.6E-41 |
| | Dtymk | 1.346 | 0.694 | 0.574 | 2.7E-43 | 4.1E-40 |

| | | | | | | |
|---|---|---|---|---|---|---|
| | Cybb | 1.397 | 0.993 | 0.501 | 0.0E+00 | 0.0E+00 |
| | Ctsa | 1.395 | 0.999 | 0.689 | 0.0E+00 | 0.0E+00 |
| | Fcgr2b | 1.362 | 0.991 | 0.641 | 0.0E+00 | 0.0E+00 |
| | Ms4a6d | 1.355 | 0.957 | 0.726 | 0.0E+00 | 0.0E+00 |
| | Ctsc | 1.326 | 0.970 | 0.634 | 0.0E+00 | 0.0E+00 |
| | Rpl3 | 1.324 | 0.999 | 0.701 | 0.0E+00 | 0.0E+00 |
| | Tmem256 | 1.321 | 0.978 | 0.530 | 0.0E+00 | 0.0E+00 |
| | Erp29 | 1.320 | 0.994 | 0.701 | 0.0E+00 | 0.0E+00 |
| | Lrpap1 | 1.315 | 0.993 | 0.571 | 0.0E+00 | 0.0E+00 |
| | Gpx1 | 1.307 | 1.000 | 0.828 | 0.0E+00 | 0.0E+00 |
| | Prdx1 | 1.306 | 0.998 | 0.653 | 0.0E+00 | 0.0E+00 |
| | Selenop | 1.301 | 0.939 | 0.505 | 0.0E+00 | 0.0E+00 |
| | Ccl6 | 1.287 | 0.993 | 0.618 | 0.0E+00 | 0.0E+00 |
| | Itgb5 | 1.270 | 0.995 | 0.664 | 0.0E+00 | 0.0E+00 |
| | Snx5 | 1.244 | 0.989 | 0.529 | 0.0E+00 | 0.0E+00 |
| | Gusb | 1.240 | 0.981 | 0.609 | 0.0E+00 | 0.0E+00 |
| | Aprt | 1.235 | 0.992 | 0.727 | 0.0E+00 | 0.0E+00 |
| | Dbi | 1.213 | 0.986 | 0.631 | 0.0E+00 | 0.0E+00 |
| | Rps2 | 1.206 | 0.998 | 0.889 | 0.0E+00 | 0.0E+00 |
| | Cst3 | 1.200 | 0.999 | 0.812 | 0.0E+00 | 0.0E+00 |
| | Atp5g1 | 1.188 | 0.996 | 0.631 | 0.0E+00 | 0.0E+00 |
| | Plekho1 | 1.164 | 0.955 | 0.538 | 0.0E+00 | 0.0E+00 |
| | Myof | 1.160 | 0.994 | 0.484 | 0.0E+00 | 0.0E+00 |
| | Abca1 | 1.158 | 0.989 | 0.723 | 0.0E+00 | 0.0E+00 |
| | Mpeg1 | 1.121 | 0.999 | 0.603 | 0.0E+00 | 0.0E+00 |
| | Creg1 | 1.120 | 0.999 | 0.654 | 0.0E+00 | 0.0E+00 |
| | Irf8 | 1.119 | 0.934 | 0.623 | 0.0E+00 | 0.0E+00 |
| | Cd93 | 1.117 | 0.916 | 0.618 | 0.0E+00 | 0.0E+00 |
| | Lamtor4 | 1.117 | 0.985 | 0.692 | 0.0E+00 | 0.0E+00 |
| | Pdia6 | 1.103 | 0.993 | 0.775 | 0.0E+00 | 0.0E+00 |
| | Fam20c | 1.100 | 0.997 | 0.755 | 0.0E+00 | 0.0E+00 |
| | Hebp1 | 1.091 | 0.971 | 0.633 | 0.0E+00 | 0.0E+00 |
| | Hspe1 | 1.089 | 0.996 | 0.660 | 0.0E+00 | 0.0E+00 |
| | Fam96a | 1.086 | 0.958 | 0.547 | 0.0E+00 | 0.0E+00 |
| | Tubb5 | 1.084 | 0.991 | 0.582 | 0.0E+00 | 0.0E+00 |
| | Sash1 | 1.076 | 0.988 | 0.700 | 0.0E+00 | 0.0E+00 |
| | Msr1 | 1.069 | 0.965 | 0.286 | 0.0E+00 | 0.0E+00 |
| | Nme1 | 1.065 | 0.992 | 0.702 | 0.0E+00 | 0.0E+00 |
| | Fkbp2 | 1.048 | 0.993 | 0.516 | 0.0E+00 | 0.0E+00 |
| | Tmem160 | 1.046 | 0.970 | 0.564 | 0.0E+00 | 0.0E+00 |
| | Gyg | 1.041 | 0.960 | 0.639 | 0.0E+00 | 0.0E+00 |
| | Gatm | 1.037 | 0.904 | 0.510 | 0.0E+00 | 0.0E+00 |
| | Anxa4 | 1.008 | 0.987 | 0.484 | 0.0E+00 | 0.0E+00 |
| | Pgls | 1.005 | 0.988 | 0.619 | 0.0E+00 | 0.0E+00 |
| | Pid1 | 1.002 | 0.863 | 0.576 | 0.0E+00 | 0.0E+00 |
| | Ppp1r14b | 1.000 | 0.991 | 0.648 | 0.0E+00 | 0.0E+00 |
| | Ucp2 | 0.994 | 0.999 | 0.719 | 0.0E+00 | 0.0E+00 |
| | Aldh2 | 0.986 | 0.970 | 0.496 | 0.0E+00 | 0.0E+00 |
| | Tuba1b | 0.983 | 0.991 | 0.583 | 0.0E+00 | 0.0E+00 |
| | Timp2 | 0.980 | 0.994 | 0.767 | 0.0E+00 | 0.0E+00 |
| | Tmed3 | 0.973 | 0.994 | 0.602 | 0.0E+00 | 0.0E+00 |
| | Zeb2 | 0.969 | 0.991 | 0.611 | 0.0E+00 | 0.0E+00 |
| | Sgk1 | 0.967 | 0.986 | 0.632 | 0.0E+00 | 0.0E+00 |
| | Lat2 | 0.960 | 0.968 | 0.664 | 0.0E+00 | 0.0E+00 |
| | Hal | 0.960 | 0.984 | 0.673 | 0.0E+00 | 0.0E+00 |
| | Ccr2 | 0.957 | 0.715 | 0.248 | 2.0E-241 | 3.0E-238 |
| | Ccl2 | 1.048 | 0.587 | 0.454 | 4.3E-39 | 6.5E-36 |
| **Macs 2** | Chil3 | 4.719 | 0.975 | 0.671 | 0.0E+00 | 0.0E+00 |
| | Gpnmb | 3.826 | 1.000 | 0.645 | 0.0E+00 | 0.0E+00 |
| | Lpl | 3.697 | 0.989 | 0.496 | 0.0E+00 | 0.0E+00 |
| | Ctsk | 3.692 | 0.959 | 0.550 | 0.0E+00 | 0.0E+00 |
| | Mmp12 | 3.321 | 0.870 | 0.537 | 0.0E+00 | 0.0E+00 |
| | Lyz2 | 3.173 | 1.000 | 0.752 | 0.0E+00 | 0.0E+00 |
| | Psap | 3.172 | 1.000 | 0.749 | 0.0E+00 | 0.0E+00 |
| | Cybb | 2.910 | 1.000 | 0.504 | 0.0E+00 | 0.0E+00 |
| | Crip1 | 2.846 | 0.996 | 0.490 | 0.0E+00 | 0.0E+00 |
| | Atp6v0d2 | 2.784 | 0.999 | 0.677 | 0.0E+00 | 0.0E+00 |
| | Spp1 | 2.744 | 0.996 | 0.782 | 0.0E+00 | 0.0E+00 |
| | Mfge8 | 2.720 | 0.895 | 0.672 | 0.0E+00 | 0.0E+00 |
| | Ctss | 2.600 | 0.999 | 0.649 | 0.0E+00 | 0.0E+00 |
| | Fabp5 | 2.562 | 0.999 | 0.797 | 0.0E+00 | 0.0E+00 |
| | Ctsl | 2.553 | 0.996 | 0.841 | 0.0E+00 | 0.0E+00 |
| | Ctsd | 2.452 | 1.000 | 0.964 | 0.0E+00 | 0.0E+00 |

| | | | | | | |
|---|---|---|---|---|---|---|
| | Camp | 4.266 | 0.267 | 0.373 | 3.4E-41 | 5.1E-38 |
| | Hist1h2ap | 1.712 | 0.662 | 0.481 | 8.0E-37 | 1.2E-33 |
| | Erp29 | 1.042 | 0.758 | 0.715 | 5.1E-28 | 7.7E-25 |
| | Ube2c | 1.866 | 0.621 | 0.424 | 1.2E-24 | 1.7E-21 |
| | Serpinb1a | 1.100 | 0.363 | 0.542 | 1.4E-23 | 2.2E-20 |
| | Ngp | 3.416 | 0.294 | 0.366 | 1.9E-22 | 2.8E-19 |
| | H2afx | 1.423 | 0.598 | 0.441 | 1.4E-15 | 2.1E-12 |
| | Prc1 | 1.160 | 0.595 | 0.429 | 2.0E-15 | 2.9E-12 |
| | Hist1h2ae | 1.473 | 0.600 | 0.599 | 3.2E-09 | 4.8E-06 |
| | Hist1h1b | 2.008 | 0.556 | 0.500 | 1.6E-06 | 2.4E-03 |
| | Car2 | 1.424 | 0.451 | 0.501 | 5.3E-05 | 7.9E-02 |
| | Smc2 | 1.419 | 0.545 | 0.391 | 1.3E-04 | 2.0E-01 |
| | Jchain | 1.375 | 0.566 | 0.380 | 9.6E-04 | 1.0E+00 |
| **Neu-5 IFN** | Rsad2 | 3.882 | 0.893 | 0.683 | 2.2E-159 | 3.2E-156 |
| | Isg15 | 2.862 | 0.914 | 0.683 | 9.1E-136 | 1.4E-132 |
| | Ifitm3 | 2.426 | 0.937 | 0.699 | 9.8E-131 | 1.5E-127 |
| | Gbp2 | 3.508 | 0.883 | 0.647 | 3.1E-123 | 4.6E-120 |
| | Slfn4 | 3.032 | 0.848 | 0.564 | 3.0E-118 | 4.5E-115 |
| | Ifi47 | 2.533 | 0.858 | 0.647 | 4.5E-108 | 6.7E-105 |
| | Ifit2 | 1.494 | 0.827 | 0.689 | 1.6E-98 | 2.5E-95 |
| | Rtp4 | 2.650 | 0.820 | 0.679 | 1.4E-96 | 2.1E-93 |
| | Ifit1bl2 | 1.106 | 0.794 | 0.562 | 1.8E-92 | 2.6E-89 |
| | Slfn5 | 2.500 | 0.787 | 0.570 | 5.6E-89 | 8.5E-86 |
| | Ifit3b | 2.020 | 0.698 | 0.421 | 5.4E-66 | 8.2E-63 |
| | Ccl4 | 1.652 | 0.921 | 0.717 | 1.5E-61 | 2.2E-58 |
| | Acod1 | 1.515 | 0.764 | 0.419 | 5.1E-59 | 7.7E-56 |
| | Ifit1 | 2.933 | 0.693 | 0.445 | 1.8E-55 | 2.7E-52 |
| | Isg20 | 2.176 | 0.731 | 0.572 | 2.0E-51 | 3.0E-48 |
| | Cldn1 | 0.347 | 0.777 | 0.616 | 3.0E-47 | 4.4E-44 |
| | Fcgr4 | 1.090 | 0.812 | 0.694 | 1.6E-45 | 2.3E-42 |
| | Zbp1 | 1.271 | 0.789 | 0.633 | 5.8E-45 | 8.7E-42 |
| | Cmpk2 | 1.335 | 0.647 | 0.457 | 6.9E-42 | 1.0E-38 |
| | Wfdc17 | 0.879 | 0.916 | 0.778 | 1.7E-38 | 2.6E-35 |
| | Oasl1 | 2.003 | 0.668 | 0.582 | 1.0E-34 | 1.5E-31 |
| | Hcar2 | 0.498 | 0.830 | 0.679 | 1.6E-30 | 2.4E-27 |
| | Gbp5 | 1.549 | 0.660 | 0.546 | 6.0E-30 | 9.0E-27 |
| | Ifit3 | 2.987 | 0.612 | 0.399 | 9.5E-30 | 1.4E-26 |
| | Gm4316 | 0.371 | 0.279 | 0.493 | 1.4E-29 | 2.1E-26 |
| | Gm12840 | 0.275 | 0.201 | 0.399 | 4.9E-28 | 7.3E-25 |
| | Retnlg | 0.406 | 0.835 | 0.628 | 9.9E-28 | 1.5E-24 |
| | G0s2 | 1.012 | 0.711 | 0.497 | 7.3E-27 | 1.1E-23 |
| | Fyb | 0.799 | 0.835 | 0.695 | 1.1E-26 | 1.6E-23 |
| | Ffar2 | 0.272 | 0.701 | 0.484 | 2.4E-26 | 3.6E-23 |
| | Cd274 | 1.280 | 0.736 | 0.593 | 5.4E-26 | 8.2E-23 |
| | Bst2 | 1.457 | 0.657 | 0.508 | 1.2E-23 | 1.7E-20 |
| | Egr1 | 0.572 | 0.756 | 0.554 | 1.1E-22 | 1.7E-19 |
| | Ccl3 | 0.827 | 0.799 | 0.641 | 1.1E-22 | 1.7E-19 |
| | Cxcl2 | 1.001 | 0.741 | 0.536 | 4.1E-22 | 6.2E-19 |
| | Cxcl10 | 1.921 | 0.551 | 0.311 | 1.4E-21 | 2.2E-18 |
| | Ier3 | 0.726 | 0.820 | 0.788 | 1.7E-21 | 2.5E-18 |
| | Gm20234 | 0.309 | 0.609 | 0.491 | 1.1E-18 | 1.6E-15 |
| | Ly6i | 0.652 | 0.586 | 0.455 | 1.5E-18 | 2.2E-15 |
| | Usp18 | 1.392 | 0.604 | 0.553 | 2.2E-18 | 3.4E-15 |
| | Clec4a3 | 0.263 | 0.614 | 0.448 | 7.7E-17 | 1.1E-13 |
| | Slpi | 0.712 | 0.797 | 0.643 | 1.8E-16 | 2.6E-13 |
| | Gm5483 | 0.695 | 0.728 | 0.616 | 2.9E-16 | 4.3E-13 |
| | Olfm4 | 0.430 | 0.635 | 0.476 | 1.4E-15 | 2.1E-12 |
| | Saa3 | 0.296 | 0.764 | 0.624 | 1.7E-15 | 2.6E-12 |
| | Asprv1 | 0.545 | 0.734 | 0.588 | 2.1E-15 | 3.1E-12 |
| | Gm19951 | 0.676 | 0.769 | 0.685 | 5.5E-15 | 8.3E-12 |
| | Ifi202b | 0.285 | 0.739 | 0.658 | 5.6E-15 | 8.4E-12 |
| | Ifitm1 | 0.593 | 0.807 | 0.738 | 8.3E-15 | 1.2E-11 |
| | Osgin1 | 0.663 | 0.678 | 0.588 | 7.4E-14 | 1.1E-10 |
| | Fgd4 | 0.355 | 0.713 | 0.634 | 7.6E-14 | 1.1E-10 |
| | Lcn2 | 0.607 | 0.683 | 0.610 | 1.0E-11 | 1.5E-08 |
| | Cst3 | 0.478 | 0.858 | 0.821 | 2.9E-11 | 4.3E-08 |
| | Fam20c | 0.267 | 0.858 | 0.766 | 2.1E-10 | 3.2E-07 |
| | Bcl2a1a | 0.523 | 0.398 | 0.556 | 5.1E-10 | 7.6E-07 |
| | Id2 | 0.498 | 0.881 | 0.780 | 1.8E-09 | 2.6E-06 |
| | Wfdc21 | 0.437 | 0.713 | 0.653 | 1.8E-09 | 2.6E-06 |
| | Il1f9 | 0.718 | 0.632 | 0.560 | 2.7E-09 | 4.0E-06 |
| | Laptm5 | 0.349 | 0.812 | 0.739 | 3.7E-08 | 5.6E-05 |
| | Id1 | 0.788 | 0.688 | 0.693 | 5.4E-08 | 8.1E-05 |

| | | | | | | |
|---|---|---|---|---|---|---|
| | Fabp4 | 2.405 | 0.976 | 0.611 | 0.0E+00 | 0.0E+00 |
| | Lipa | 2.389 | 0.997 | 0.513 | 0.0E+00 | 0.0E+00 |
| | Mgll | 2.382 | 0.973 | 0.587 | 0.0E+00 | 0.0E+00 |
| | Trem2 | 2.379 | 0.995 | 0.566 | 0.0E+00 | 0.0E+00 |
| | Mpeg1 | 2.367 | 0.999 | 0.606 | 0.0E+00 | 0.0E+00 |
| | Vim | 2.330 | 0.999 | 0.634 | 0.0E+00 | 0.0E+00 |
| | Sgk1 | 2.323 | 0.999 | 0.634 | 0.0E+00 | 0.0E+00 |
| | mt-Nd1 | 2.303 | 0.998 | 0.660 | 0.0E+00 | 0.0E+00 |
| | S100a1 | 2.291 | 0.994 | 0.634 | 0.0E+00 | 0.0E+00 |
| | Ccl6 | 2.274 | 0.994 | 0.621 | 0.0E+00 | 0.0E+00 |
| | Abcg1 | 2.256 | 0.997 | 0.645 | 0.0E+00 | 0.0E+00 |
| | Mrc1 | 2.227 | 0.894 | 0.408 | 0.0E+00 | 0.0E+00 |
| | Myof | 2.219 | 0.995 | 0.488 | 0.0E+00 | 0.0E+00 |
| | Selenop | 2.137 | 0.991 | 0.506 | 0.0E+00 | 0.0E+00 |
| | Pld3 | 2.132 | 0.995 | 0.574 | 0.0E+00 | 0.0E+00 |
| | Plin2 | 2.121 | 0.999 | 0.844 | 0.0E+00 | 0.0E+00 |
| | Apoe | 2.006 | 0.979 | 0.778 | 0.0E+00 | 0.0E+00 |
| | Anxa5 | 1.983 | 0.999 | 0.671 | 0.0E+00 | 0.0E+00 |
| | Cd68 | 1.963 | 0.998 | 0.702 | 0.0E+00 | 0.0E+00 |
| | Aig1 | 1.926 | 0.981 | 0.665 | 0.0E+00 | 0.0E+00 |
| | Mgst1 | 1.906 | 0.996 | 0.592 | 0.0E+00 | 0.0E+00 |
| | Slc7a2 | 1.859 | 0.953 | 0.593 | 0.0E+00 | 0.0E+00 |
| | Ear2 | 1.849 | 0.881 | 0.305 | 0.0E+00 | 0.0E+00 |
| | Sh3bgrl | 1.816 | 0.991 | 0.540 | 0.0E+00 | 0.0E+00 |
| | Lrp1 | 1.810 | 0.995 | 0.507 | 0.0E+00 | 0.0E+00 |
| | Hexa | 1.807 | 0.999 | 0.690 | 0.0E+00 | 0.0E+00 |
| | Gns | 1.748 | 1.000 | 0.684 | 0.0E+00 | 0.0E+00 |
| | Abhd12 | 1.739 | 0.990 | 0.418 | 0.0E+00 | 0.0E+00 |
| | Ahnak2 | 1.738 | 0.981 | 0.542 | 0.0E+00 | 0.0E+00 |
| | Creg1 | 1.723 | 0.999 | 0.657 | 0.0E+00 | 0.0E+00 |
| | Prdx1 | 1.712 | 0.996 | 0.656 | 0.0E+00 | 0.0E+00 |
| | Lgals1 | 1.693 | 0.991 | 0.563 | 0.0E+00 | 0.0E+00 |
| | Lgmn | 1.687 | 0.991 | 0.618 | 0.0E+00 | 0.0E+00 |
| | Gstm1 | 1.679 | 0.977 | 0.546 | 0.0E+00 | 0.0E+00 |
| | Trf | 1.674 | 0.974 | 0.490 | 0.0E+00 | 0.0E+00 |
| | Gm26917 | 1.673 | 0.905 | 0.556 | 0.0E+00 | 0.0E+00 |
| | Gusb | 1.658 | 0.988 | 0.611 | 0.0E+00 | 0.0E+00 |
| | Lamp1 | 1.650 | 0.999 | 0.858 | 0.0E+00 | 0.0E+00 |
| | Acp5 | 1.634 | 0.995 | 0.601 | 0.0E+00 | 0.0E+00 |
| | Vat1 | 1.632 | 0.981 | 0.642 | 0.0E+00 | 0.0E+00 |
| | F7 | 1.632 | 0.925 | 0.406 | 0.0E+00 | 0.0E+00 |
| | Axl | 1.612 | 0.892 | 0.422 | 0.0E+00 | 0.0E+00 |
| | Ctsb | 1.608 | 1.000 | 0.943 | 0.0E+00 | 0.0E+00 |
| | Ctsz | 1.605 | 1.000 | 0.787 | 0.0E+00 | 0.0E+00 |
| | Snx5 | 1.602 | 0.992 | 0.532 | 0.0E+00 | 0.0E+00 |
| | Ctsa | 1.599 | 0.999 | 0.692 | 0.0E+00 | 0.0E+00 |
| | Dusp3 | 1.564 | 0.986 | 0.628 | 0.0E+00 | 0.0E+00 |
| | Cd36 | 1.561 | 0.877 | 0.480 | 0.0E+00 | 0.0E+00 |
| | Myo5a | 1.560 | 0.983 | 0.491 | 0.0E+00 | 0.0E+00 |
| | Aplp2 | 1.550 | 0.995 | 0.630 | 0.0E+00 | 0.0E+00 |
| | Il11ra1 | 1.532 | 0.958 | 0.470 | 0.0E+00 | 0.0E+00 |
| | Kcnq1ot1 | 1.510 | 0.945 | 0.642 | 0.0E+00 | 0.0E+00 |
| | Sirpa | 1.506 | 0.996 | 0.609 | 0.0E+00 | 0.0E+00 |
| | Itgax | 1.499 | 0.986 | 0.527 | 0.0E+00 | 0.0E+00 |
| | Anxa4 | 1.494 | 0.988 | 0.487 | 0.0E+00 | 0.0E+00 |
| | Fstl1 | 1.486 | 0.862 | 0.466 | 0.0E+00 | 0.0E+00 |
| | Dnmt3a | 1.473 | 0.946 | 0.462 | 0.0E+00 | 0.0E+00 |
| | Cdo1 | 1.466 | 0.918 | 0.461 | 0.0E+00 | 0.0E+00 |
| | Shtn1 | 1.463 | 0.978 | 0.464 | 0.0E+00 | 0.0E+00 |
| | Laptm5 | 1.462 | 0.999 | 0.728 | 0.0E+00 | 0.0E+00 |
| | Grn | 1.460 | 0.996 | 0.841 | 0.0E+00 | 0.0E+00 |
| | Mertk | 1.458 | 0.955 | 0.387 | 0.0E+00 | 0.0E+00 |
| | Slc6a6 | 1.456 | 0.997 | 0.490 | 0.0E+00 | 0.0E+00 |
| | Smpdl3a | 1.454 | 0.996 | 0.614 | 0.0E+00 | 0.0E+00 |
| | Il18 | 1.438 | 0.916 | 0.498 | 0.0E+00 | 0.0E+00 |
| | Abcc5 | 1.434 | 0.946 | 0.408 | 0.0E+00 | 0.0E+00 |
| | Colgalt1 | 1.434 | 0.980 | 0.501 | 0.0E+00 | 0.0E+00 |
| | Csf2rb | 0.657 | 0.721 | 0.686 | 8.4E-08 | 1.3E-04 |
| | Bcl2a1b | 0.643 | 0.751 | 0.631 | 3.8E-07 | 5.7E-04 |
| | Ddx60 | 1.401 | 0.528 | 0.488 | 5.3E-07 | 8.0E-04 |
| | Mpeg1 | 0.362 | 0.726 | 0.622 | 1.2E-06 | 1.7E-03 |
| | Tnfrsf23 | 0.333 | 0.741 | 0.702 | 5.0E-06 | 7.5E-03 |
| | Osm | 0.260 | 0.475 | 0.334 | 5.3E-06 | 8.0E-03 |
| | Ifi27l2a | 2.007 | 0.553 | 0.533 | 5.5E-06 | 8.2E-03 |
| | Cdc42ep3 | 0.290 | 0.622 | 0.527 | 7.5E-06 | 1.1E-02 |
| | Il18bp | 0.429 | 0.508 | 0.450 | 1.3E-05 | 2.0E-02 |
| | Irf7 | 0.399 | 0.642 | 0.654 | 2.9E-05 | 4.4E-02 |
| | Lrg1 | 0.382 | 0.668 | 0.699 | 2.1E-04 | 3.1E-01 |
| | Cd300c2 | 0.587 | 0.657 | 0.660 | 2.3E-04 | 3.4E-01 |
| | Hk2 | 0.508 | 0.497 | 0.587 | 2.8E-04 | 4.2E-01 |
| | Steap4 | 0.298 | 0.586 | 0.487 | 3.4E-04 | 5.1E-01 |
| | Ifi209 | 0.898 | 0.563 | 0.599 | 4.8E-04 | 7.3E-01 |
| | Ly6c2 | 1.077 | 0.482 | 0.398 | 5.2E-04 | 7.8E-01 |
| | Chil1 | 0.308 | 0.429 | 0.495 | 7.0E-04 | 1.0E+00 |
| | Il18 | 0.297 | 0.556 | 0.516 | 8.5E-04 | 1.0E+00 |
| | Ly6g | 0.309 | 0.497 | 0.449 | 1.5E-03 | 1.0E+00 |
| | Rgs1 | 0.948 | 0.627 | 0.632 | 1.9E-03 | 1.0E+00 |
| | Unc93b1 | 0.563 | 0.569 | 0.521 | 2.0E-03 | 1.0E+00 |
| | Hist1h1c | 0.327 | 0.497 | 0.617 | 2.6E-03 | 1.0E+00 |
| | Ifitm6 | 0.346 | 0.391 | 0.472 | 4.6E-03 | 1.0E+00 |
| **Eryth.** | Hbb-bs | 9.669 | 1.000 | 0.782 | 0.0E+00 | 0.0E+00 |
| | Hba-a1 | 8.916 | 1.000 | 0.499 | 0.0E+00 | 0.0E+00 |
| | Hbb-bt | 8.788 | 0.996 | 0.481 | 0.0E+00 | 0.0E+00 |
| | Hba-a2 | 8.609 | 1.000 | 0.450 | 0.0E+00 | 0.0E+00 |
| | Bpgm | 4.001 | 0.837 | 0.481 | 0.0E+00 | 0.0E+00 |
| | Alas2 | 3.365 | 0.939 | 0.531 | 0.0E+00 | 0.0E+00 |
| | Snca | 2.876 | 0.949 | 0.366 | 0.0E+00 | 0.0E+00 |
| | Fech | 2.293 | 0.957 | 0.604 | 0.0E+00 | 0.0E+00 |
| | Fam46c | 1.861 | 0.916 | 0.610 | 0.0E+00 | 0.0E+00 |
| | Rsad2 | 1.161 | 0.865 | 0.679 | 0.0E+00 | 0.0E+00 |
| | Gypa | 0.702 | 0.791 | 0.028 | 0.0E+00 | 0.0E+00 |
| | Slc4a1 | 0.430 | 0.640 | 0.027 | 0.0E+00 | 0.0E+00 |
| | Prdx2 | 1.533 | 0.891 | 0.636 | 2.0E-288 | 3.0E-285 |
| | Fam213a | 0.874 | 0.916 | 0.568 | 2.3E-282 | 3.5E-279 |
| | Isg20 | 1.050 | 0.910 | 0.561 | 2.0E-245 | 2.9E-242 |
| | Ncoa4 | 0.766 | 0.907 | 0.558 | 3.4E-228 | 5.0E-225 |
| | Aldh1a1 | 0.377 | 0.748 | 0.376 | 5.6E-212 | 8.4E-209 |
| | Car2 | 0.406 | 0.781 | 0.489 | 6.8E-212 | 1.0E-208 |
| | Ube2c | 0.553 | 0.642 | 0.419 | 4.5E-87 | 6.8E-84 |
| | Gpx1 | 0.918 | 0.898 | 0.834 | 2.7E-17 | 4.0E-14 |
| **Macs 3** | Lyz2 | 3.103 | 0.994 | 0.755 | 0.0E+00 | 0.0E+00 |
| | Atp6v0d2 | 2.835 | 0.929 | 0.683 | 0.0E+00 | 0.0E+00 |
| | Fabp5 | 2.818 | 0.976 | 0.801 | 0.0E+00 | 0.0E+00 |
| | Fabp4 | 2.446 | 0.907 | 0.618 | 0.0E+00 | 0.0E+00 |
| | Trem2 | 2.287 | 0.948 | 0.573 | 0.0E+00 | 0.0E+00 |
| | Mmp12 | 2.260 | 0.889 | 0.540 | 0.0E+00 | 0.0E+00 |
| | Vim | 2.236 | 0.943 | 0.640 | 0.0E+00 | 0.0E+00 |
| | Gpnmb | 2.203 | 0.955 | 0.651 | 0.0E+00 | 0.0E+00 |
| | Ftl1 | 1.976 | 1.000 | 0.956 | 0.0E+00 | 0.0E+00 |
| | C1qa | 1.299 | 0.907 | 0.583 | 0.0E+00 | 0.0E+00 |
| | Il11ra1 | 1.068 | 0.868 | 0.479 | 1.4E-302 | 2.1E-299 |
| | Lgals1 | 2.459 | 0.899 | 0.571 | 2.8E-293 | 4.2E-290 |
| | C1qb | 1.319 | 0.890 | 0.505 | 4.2E-288 | 6.2E-285 |
| | Lpl | 2.532 | 0.875 | 0.506 | 1.6E-278 | 2.4E-275 |
| | Cdkn2a | 0.952 | 0.852 | 0.657 | 1.9E-278 | 2.9E-275 |
| | Pld3 | 1.832 | 0.903 | 0.582 | 6.5E-275 | 9.7E-272 |
| | Crip1 | 2.187 | 0.866 | 0.500 | 1.9E-269 | 2.8E-266 |
| | Serpinb6a | 2.298 | 0.917 | 0.672 | 1.5E-268 | 2.3E-265 |
| | Spp1 | 2.531 | 0.948 | 0.786 | 4.3E-264 | 6.4E-261 |
| | Prdx1 | 2.470 | 0.877 | 0.664 | 4.4E-258 | 6.7E-255 |
| | Sdc3 | 1.041 | 0.862 | 0.567 | 9.7E-254 | 1.5E-250 |
| | S100a1 | 2.732 | 0.862 | 0.643 | 1.9E-248 | 2.8E-245 |
| | Myof | 1.428 | 0.860 | 0.499 | 6.7E-247 | 1.0E-243 |
| | Ctsk | 2.393 | 0.836 | 0.559 | 2.3E-238 | 3.5E-235 |

| | | | | | | | | | | | | | |
|---|---|---|---|---|---|---|---|---|---|---|---|---|---|
| | Ucp2 | 1.423 | 0.999 | 0.721 | 0.0E+00 | 0.0E+00 | | Mt2 | 1.490 | 0.817 | 0.645 | 2.6E-238 | 3.8E-235 |
| | Serpinb6a | 1.419 | 0.997 | 0.665 | 0.0E+00 | 0.0E+00 | | Ccl6 | 2.295 | 0.913 | 0.628 | 1.9E-236 | 2.9E-233 |
| | Itgb2 | 1.419 | 0.996 | 0.577 | 0.0E+00 | 0.0E+00 | | Cd63 | 2.069 | 0.968 | 0.778 | 4.8E-228 | 7.2E-225 |
| | Cd63 | 1.419 | 0.999 | 0.774 | 0.0E+00 | 0.0E+00 | | Apoe | 1.530 | 0.947 | 0.782 | 7.2E-227 | 1.1E-223 |
| | Dbi | 1.415 | 0.987 | 0.634 | 0.0E+00 | 0.0E+00 | | Chil3 | 2.328 | 0.881 | 0.677 | 1.7E-224 | 2.5E-221 |
| | Tgfbr2 | 1.413 | 0.988 | 0.488 | 0.0E+00 | 0.0E+00 | | C1qc | 1.093 | 0.861 | 0.644 | 9.6E-215 | 1.4E-211 |
| | Sdc3 | 1.412 | 0.957 | 0.559 | 0.0E+00 | 0.0E+00 | | Lgmn | 1.340 | 0.917 | 0.625 | 6.9E-209 | 1.0E-205 |
| | Npc2 | 1.410 | 1.000 | 0.886 | 0.0E+00 | 0.0E+00 | | Plin2 | 1.944 | 0.957 | 0.847 | 9.8E-201 | 1.5E-197 |
| | Dhrs3 | 1.394 | 0.995 | 0.661 | 0.0E+00 | 0.0E+00 | | Mfge8 | 2.225 | 0.887 | 0.675 | 2.6E-200 | 3.9E-197 |
| | Serpine1 | 1.356 | 0.860 | 0.281 | 0.0E+00 | 0.0E+00 | | Gngt2 | 2.023 | 0.880 | 0.639 | 1.5E-198 | 2.2E-195 |
| | Syngr1 | 1.354 | 0.971 | 0.762 | 0.0E+00 | 0.0E+00 | | Cd36 | 1.210 | 0.816 | 0.487 | 1.8E-196 | 2.6E-193 |
| | Atp13a2 | 1.353 | 0.978 | 0.468 | 0.0E+00 | 0.0E+00 | | Bhlhe41 | 1.075 | 0.805 | 0.561 | 8.0E-189 | 1.2E-185 |
| | Tcf7l2 | 1.342 | 0.915 | 0.455 | 0.0E+00 | 0.0E+00 | | Syngr1 | 1.161 | 0.875 | 0.768 | 9.8E-184 | 1.5E-180 |
| | Sort1 | 1.301 | 0.946 | 0.559 | 0.0E+00 | 0.0E+00 | | Mt1 | 2.231 | 0.865 | 0.772 | 4.2E-183 | 6.3E-180 |
| | Lrpap1 | 1.294 | 0.948 | 0.577 | 0.0E+00 | 0.0E+00 | | Psap | 1.901 | 0.939 | 0.754 | 5.6E-181 | 8.4E-178 |
| | Bhlhe41 | 1.284 | 0.943 | 0.552 | 0.0E+00 | 0.0E+00 | | Comt | 1.663 | 0.818 | 0.624 | 8.2E-180 | 1.2E-176 |
| | AU020206 | 1.254 | 0.941 | 0.692 | 0.0E+00 | 0.0E+00 | | Ucp2 | 1.750 | 0.878 | 0.729 | 1.1E-168 | 1.6E-165 |
| **Neu-4 classic** | Retnlg | 4.320 | 0.993 | 0.616 | 0.0E+00 | 0.0E+00 | | Ctsz | 1.495 | 0.959 | 0.791 | 1.0E-166 | 1.5E-163 |
| | Ifitm6 | 3.584 | 0.992 | 0.450 | 0.0E+00 | 0.0E+00 | | Vat1 | 1.214 | 0.850 | 0.651 | 4.9E-166 | 7.4E-163 |
| | Lcn2 | 3.566 | 0.999 | 0.596 | 0.0E+00 | 0.0E+00 | | Ctss | 1.571 | 0.856 | 0.658 | 2.9E-164 | 4.3E-161 |
| | Wfdc21 | 3.439 | 0.999 | 0.640 | 0.0E+00 | 0.0E+00 | | Rps2 | 1.322 | 0.923 | 0.894 | 6.8E-152 | 1.0E-148 |
| | Mmp8 | 2.882 | 0.971 | 0.602 | 0.0E+00 | 0.0E+00 | | Mgll | 1.555 | 0.775 | 0.599 | 2.6E-149 | 3.9E-146 |
| | Wfdc17 | 2.808 | 0.997 | 0.771 | 0.0E+00 | 0.0E+00 | | Lipa | 1.354 | 0.858 | 0.524 | 4.6E-147 | 6.9E-144 |
| | Ifitm1 | 2.636 | 0.992 | 0.728 | 0.0E+00 | 0.0E+00 | | Nme1 | 1.536 | 0.864 | 0.712 | 2.6E-139 | 3.8E-136 |
| | Lrg1 | 2.580 | 0.992 | 0.687 | 0.0E+00 | 0.0E+00 | | Npc2 | 1.668 | 0.952 | 0.889 | 1.9E-138 | 2.9E-135 |
| | Prok2 | 2.540 | 0.960 | 0.568 | 0.0E+00 | 0.0E+00 | | Cd68 | 1.715 | 0.860 | 0.710 | 4.7E-137 | 7.0E-134 |
| | Ly6g | 2.372 | 0.944 | 0.430 | 0.0E+00 | 0.0E+00 | | Ndufc2 | 1.120 | 0.806 | 0.620 | 1.0E-130 | 1.6E-127 |
| | Slpi | 1.882 | 0.978 | 0.632 | 0.0E+00 | 0.0E+00 | | Ctsd | 1.460 | 0.982 | 0.965 | 5.3E-126 | 8.0E-123 |
| | Ifitm3 | 1.752 | 0.989 | 0.691 | 0.0E+00 | 0.0E+00 | | Chchd10 | 0.992 | 0.770 | 0.640 | 1.8E-116 | 2.7E-113 |
| | Stfa2 | 1.639 | 0.909 | 0.508 | 0.0E+00 | 0.0E+00 | | Acp5 | 1.333 | 0.859 | 0.610 | 1.3E-111 | 2.0E-108 |
| | Cd177 | 1.499 | 0.907 | 0.482 | 0.0E+00 | 0.0E+00 | | Blvra | 1.204 | 0.765 | 0.688 | 3.1E-107 | 4.6E-104 |
| | Chil1 | 1.440 | 0.942 | 0.477 | 0.0E+00 | 0.0E+00 | | Ckb | 1.006 | 0.730 | 0.512 | 4.7E-105 | 7.1E-102 |
| | Anxa1 | 1.274 | 0.997 | 0.792 | 0.0E+00 | 0.0E+00 | | Lrpap1 | 1.029 | 0.784 | 0.587 | 1.6E-100 | 2.4E-97 |
| | Steap4 | 1.122 | 0.940 | 0.470 | 0.0E+00 | 0.0E+00 | | Marco | 1.092 | 0.638 | 0.382 | 1.4E-99 | 2.1E-96 |
| | Ggt1 | 1.101 | 0.897 | 0.480 | 0.0E+00 | 0.0E+00 | | Hebp1 | 1.673 | 0.752 | 0.647 | 4.2E-98 | 6.3E-95 |
| | Il1f9 | 0.835 | 0.940 | 0.545 | 3.3E-294 | 4.9E-291 | | Mrc1 | 1.060 | 0.678 | 0.421 | 4.6E-96 | 6.8E-93 |
| | Slfn4 | 0.805 | 0.912 | 0.554 | 3.6E-286 | 5.4E-283 | | Creg1 | 1.335 | 0.866 | 0.665 | 2.1E-95 | 3.1E-92 |
| | Tgm1 | 0.511 | 0.865 | 0.592 | 3.6E-260 | 5.4E-257 | | Mpeg1 | 1.104 | 0.872 | 0.615 | 1.0E-91 | 1.5E-88 |
| | Ngp | 1.375 | 0.786 | 0.348 | 1.6E-249 | 2.3E-246 | | Rpl3 | 1.021 | 0.848 | 0.712 | 2.1E-91 | 3.2E-88 |
| | Gm5483 | 1.224 | 0.916 | 0.606 | 2.1E-249 | 3.2E-246 | | Ear2 | 1.521 | 0.627 | 0.320 | 7.5E-87 | 1.1E-83 |
| | Gbp2 | 0.381 | 0.899 | 0.640 | 3.9E-247 | 5.9E-244 | | Gpx1 | 1.184 | 0.891 | 0.835 | 1.3E-82 | 2.0E-79 |
| | BC100530 | 2.269 | 0.877 | 0.550 | 4.7E-247 | 7.0E-244 | | Ctsl | 1.931 | 0.937 | 0.845 | 3.9E-82 | 5.8E-79 |
| | Tgfbi | 0.991 | 0.922 | 0.598 | 1.6E-246 | 2.5E-243 | | Anxa4 | 1.516 | 0.722 | 0.503 | 2.2E-78 | 3.2E-75 |
| | Flna | 0.907 | 0.875 | 0.483 | 5.7E-215 | 8.5E-212 | | Gyg | 1.106 | 0.792 | 0.651 | 6.1E-76 | 9.1E-73 |
| | Csf2rb | 0.767 | 0.972 | 0.676 | 1.4E-214 | 2.1E-211 | | Akr1b3 | 1.361 | 0.704 | 0.519 | 2.5E-74 | 3.7E-71 |
| | Hacd4 | 1.020 | 0.822 | 0.508 | 3.9E-214 | 5.8E-211 | | Abcg1 | 1.286 | 0.825 | 0.655 | 1.8E-73 | 2.7E-70 |
| | Ccl6 | 0.890 | 0.959 | 0.624 | 1.5E-213 | 2.3E-210 | | Cybb | 1.621 | 0.701 | 0.520 | 1.8E-71 | 2.7E-68 |
| | Gyg | 0.741 | 0.863 | 0.647 | 1.3E-200 | 2.0E-197 | | Anxa5 | 1.578 | 0.758 | 0.684 | 1.8E-70 | 2.7E-67 |
| | Stfa2l1 | 1.303 | 0.835 | 0.467 | 2.0E-193 | 2.9E-190 | | Il18 | 1.245 | 0.662 | 0.512 | 9.3E-69 | 1.4E-65 |
| | Pi16 | 0.851 | 0.785 | 0.460 | 4.3E-188 | 6.4E-185 | | Ftl1-ps1 | 0.990 | 0.663 | 0.422 | 6.5E-67 | 9.7E-64 |
| | Tceal9 | 0.260 | 0.937 | 0.666 | 1.7E-185 | 2.6E-182 | | Lamtor4 | 1.225 | 0.821 | 0.703 | 5.3E-66 | 7.9E-63 |
| | Id1 | 0.309 | 0.894 | 0.685 | 8.3E-174 | 1.2E-170 | | Mgst1 | 1.663 | 0.691 | 0.607 | 2.6E-64 | 4.0E-61 |
| | Syne1 | 0.622 | 0.810 | 0.599 | 1.2E-162 | 1.8E-159 | | Hexa | 1.098 | 0.839 | 0.699 | 4.3E-62 | 6.4E-59 |
| | F630028O10Ri | 0.753 | 0.828 | 0.534 | 8.0E-158 | 1.2E-154 | | Sh3bgrl | 1.604 | 0.671 | 0.556 | 1.4E-58 | 2.1E-55 |
| | Mgst1 | 0.476 | 0.870 | 0.599 | 2.7E-148 | 4.1E-145 | | Cndp2 | 1.101 | 0.673 | 0.547 | 2.4E-57 | 3.5E-54 |
| | Gadd45a | 0.597 | 0.832 | 0.616 | 9.7E-141 | 1.5E-137 | | Lmna | 1.256 | 0.730 | 0.630 | 2.9E-50 | 4.4E-47 |
| | Tuba1a | 0.562 | 0.780 | 0.518 | 2.4E-137 | 3.7E-134 | | Gstm1 | 1.617 | 0.640 | 0.563 | 2.5E-49 | 3.7E-46 |
| | Napsa | 0.305 | 0.855 | 0.579 | 6.0E-121 | 9.0E-118 | | Ptms | 1.387 | 0.727 | 0.623 | 3.7E-49 | 5.5E-46 |
| | Adam8 | 0.628 | 0.807 | 0.586 | 3.7E-118 | 5.5E-115 | | Sept9 | 1.139 | 0.627 | 0.473 | 6.6E-45 | 9.8E-42 |
| | Ly6c2 | 0.635 | 0.708 | 0.387 | 1.9E-116 | 2.9E-113 | | Lamp1 | 0.986 | 0.887 | 0.863 | 7.9E-42 | 1.2E-38 |
| | Arhgap25 | 0.285 | 0.808 | 0.494 | 8.0E-109 | 1.2E-105 | | Nenf | 0.948 | 0.658 | 0.522 | 8.2E-42 | 1.2E-38 |
| | Fgd4 | 0.382 | 0.790 | 0.629 | 7.6E-99 | 1.1E-95 | | Selenop | 1.239 | 0.639 | 0.523 | 4.1E-40 | 6.2E-37 |
| | Lmo4 | 0.595 | 0.715 | 0.404 | 6.3E-96 | 9.5E-93 | | Ctsb | 1.179 | 0.945 | 0.945 | 9.1E-39 | 1.4E-35 |
| | Smpdl3a | 0.448 | 0.826 | 0.623 | 7.2E-91 | 1.1E-87 | | Dbi | 1.285 | 0.650 | 0.649 | 5.4E-32 | 8.1E-29 |
| | Itgb2 | 0.339 | 0.867 | 0.585 | 4.4E-86 | 6.6E-83 | | Ccl9 | 1.120 | 0.623 | 0.545 | 1.0E-31 | 1.5E-28 |
| | Acvrl1 | 0.628 | 0.673 | 0.488 | 1.1E-76 | 1.7E-73 | | Ctsa | 0.950 | 0.810 | 0.702 | 4.3E-31 | 6.5E-28 |
| | Saa3 | 0.352 | 0.854 | 0.617 | 1.3E-74 | 2.0E-71 | | Camk1 | 1.023 | 0.632 | 0.532 | 6.6E-31 | 9.9E-28 |
| | Sirpa | 0.296 | 0.858 | 0.617 | 5.0E-72 | 7.5E-69 | | Hspe1 | 1.301 | 0.691 | 0.676 | 9.4E-31 | 1.4E-27 |
| | Olfm4 | 1.468 | 0.683 | 0.470 | 6.3E-67 | 9.5E-64 | | Smpdl3a | 0.946 | 0.721 | 0.628 | 2.8E-25 | 4.3E-22 |
| | Asprv1 | 0.875 | 0.706 | 0.585 | 2.1E-58 | 3.2E-55 | | Pgls | 1.000 | 0.682 | 0.636 | 5.1E-23 | 7.6E-20 |

| | | | | | | | | | | | |
|---|---|---|---|---|---|---|---|---|---|---|---|
| Olfm4 | 1.468 | 0.683 | 0.470 | 6.3E-67 | 9.5E-64 | Smpdl3a | 0.946 | 0.721 | 0.628 | 2.8E-25 | 4.3E-22 |
| Asprv1 | 0.875 | 0.706 | 0.585 | 2.1E-58 | 3.2E-55 | Pgls | 1.000 | 0.682 | 0.636 | 5.1E-23 | 7.6E-20 |
| Camp | 0.968 | 0.597 | 0.362 | 2.9E-50 | 4.4E-47 | Atp5g1 | 1.239 | 0.649 | 0.649 | 5.3E-21 | 8.0E-18 |
| Serpinb1a | 0.640 | 0.616 | 0.536 | 9.8E-38 | 1.5E-34 | Trf | 0.948 | 0.589 | 0.509 | 6.0E-18 | 9.0E-15 |
| Cfp | 0.367 | 0.625 | 0.445 | 1.9E-37 | 2.9E-34 | Fam96a | 0.940 | 0.409 | 0.573 | 6.5E-15 | 9.8E-12 |
| Stfa3 | 0.514 | 0.382 | 0.548 | 5.2E-34 | 7.8E-31 | Ppp1r14b | 1.198 | 0.602 | 0.668 | 1.6E-10 | 2.4E-07 |
| Abcd2 | 0.264 | 0.621 | 0.524 | 2.9E-28 | 4.3E-25 | Rexo2 | 1.352 | 0.558 | 0.640 | 1.2E-09 | 1.8E-06 |
| Dgat2 | 0.299 | 0.683 | 0.628 | 9.4E-26 | 1.4E-22 | Aig1 | 1.010 | 0.561 | 0.683 | 5.7E-03 | 1.0E+00 |
| Rab27a | 0.295 | 0.545 | 0.430 | 4.6E-11 | 6.8E-08 | | | | | | |
| Osm | 0.446 | 0.514 | 0.328 | 4.6E-06 | 7.0E-03 | | | | | | |

**Supplemental Table 4: GSEA on scRNAseq Populations, related to Figure 6**
**NES=Normalized Enrichment Score, FDR=False Discovery Rate**

| | | Combo vs aPD1 | | | | | | | | |
|---|---|---|---|---|---|---|---|---|---|---|
| | | Macs/Dend | | | Neus | | | Tumor | | |
| Group | MSigDB Signature Name | NES | FDR q | -Log(q) | NES | FDR q | -Log(q) | NES | FDR q | -Log(q) |
| DNA Replication and Damage | REACTOME_G1_S_DNA_DAMAGE_CHECKPOINTS.v2022.1.Hs.grp | 1.64 | 0.02 | 1.65 | 1.30 | 0.18 | 0.74 | 1.46 | 0.03 | 1.57 |
| | REACTOME_ORC1_REMOVAL_FROM_CHROMATIN.v2022.1.Hs.grp | 1.51 | 0.04 | 1.36 | 1.64 | 0.04 | 1.39 | 1.49 | 0.02 | 1.68 |
| | REACTOME_SWITCHING_OF_ORIGINS_TO_A_POST_REPLICATIVE_STATE.v2022.1.Hs.grp | 1.42 | 0.07 | 1.15 | 1.59 | 0.05 | 1.33 | 1.43 | 0.03 | 1.46 |
| | REACTOME_CDT1_ASSOCIATION_WITH_THE_CDC6_ORC_ORIGIN_COMPLEX.v2022.1.Hs.grp | 1.55 | 0.03 | 1.46 | 1.69 | 0.04 | 1.40 | 1.41 | 0.04 | 1.36 |
| | REACTOME_SYNTHESIS_OF_DNA.v2022.1.Hs.grp | 1.37 | 0.10 | 1.01 | 1.63 | 0.04 | 1.39 | 1.37 | 0.06 | 1.23 |
| | REACTOME_APC_C_MEDIATED_DEGRADATION_OF_CELL_CYCLE_PROTEINS.v2022.1.Hs.grp | 1.45 | 0.06 | 1.24 | 1.52 | 0.07 | 1.18 | 1.40 | 0.05 | 1.34 |
| Protein and RNA Processing | KEGG_RIBOSOME.v2022.1.Hs.grp | 2.56 | 0.00 | 4.00 | 1.61 | 0.04 | 1.37 | -2.28 | 0.00 | 4.00 |
| | REACTOME_EUKARYOTIC_TRANSLATION_ELONGATION.v2022.1.Hs.grp | 2.56 | 0.00 | 4.00 | 1.52 | 0.07 | 1.17 | -2.42 | 0.00 | 4.00 |
| | Hs.grp | 2.54 | 0.00 | 4.00 | 1.64 | 0.04 | 1.39 | -2.02 | 0.00 | 2.85 |
| | REACTOME_RESPONSE_OF_EIF2AK4_GCN2_TO_AMINO_ACID_DEFICIENCY.v2022.1.Hs.grp | 2.41 | 0.00 | 4.00 | 1.36 | 0.14 | 0.84 | -2.12 | 0.00 | 3.34 |
| | REACTOME_EUKARYOTIC_TRANSLATION_INITIATION.v2022.1.Hs.grp | 2.40 | 0.00 | 4.00 | 1.25 | 0.22 | 0.65 | -2.11 | 0.00 | 3.37 |
| | REACTOME_INFLUENZA_INFECTION.v2022.1.Hs.grp | 2.40 | 0.00 | 4.00 | 1.27 | 0.21 | 0.67 | -2.19 | 0.00 | 3.84 |
| | GOBP_CYTOPLASMIC_TRANSLATION.v2022.1.Hs.grp | 2.24 | 0.00 | 4.00 | 1.04 | 0.45 | 0.34 | -1.94 | 0.00 | 2.60 |
| | REACTOME_TRANSLATION.v2022.1.Hs.grp | 2.01 | 0.00 | 3.05 | 1.23 | 0.24 | 0.61 | -1.62 | 0.02 | 1.69 |
| | REACTOME_SELENOAMINO_ACID_METABOLISM.v2022.1.Hs.grp | 2.38 | 0.00 | 4.00 | 1.51 | 0.07 | 1.17 | -2.38 | 0.00 | 4.00 |
| | REACTOME_CELLULAR_RESPONSE_TO_STARVATION.v2022.1.Hs.grp | 2.30 | 0.00 | 4.00 | 1.04 | 0.45 | 0.35 | -1.80 | 0.01 | 2.14 |
| | REACTOME_METABOLISM_OF_AMINO_ACIDS_AND_DERIVATIVES.v2022.1.Hs.grp | 1.98 | 0.00 | 2.99 | 1.22 | 0.25 | 0.61 | -1.56 | 0.03 | 1.54 |
| | REACTOME_NONSENSE_MEDIATED_DECAY_NMD.v2022.1.Hs.grp | 2.48 | 0.00 | 4.00 | 1.42 | 0.11 | 0.96 | -2.21 | 0.00 | 4.00 |
| | REACTOME_RRNA_PROCESSING.v2022.1.Hs.grp | 2.19 | 0.00 | 4.00 | 1.40 | 0.12 | 0.91 | -2.01 | 0.00 | 2.79 |
| Oxidative Phosphorylation | GOBP_AEROBIC_RESPIRATION.v2022.1.Hs.grp | 1.77 | 0.01 | 2.11 | 2.53 | 0.00 | 4.00 | 1.63 | 0.01 | 2.28 |
| | GOBP_OXIDATIVE_PHOSPHORYLATION.v2022.1.Hs.grp | 1.95 | 0.00 | 2.80 | 2.62 | 0.00 | 4.00 | 1.72 | 0.00 | 2.67 |
| | GOBP_ATP_SYNTHESIS_COUPLED_ELECTRON_TRANSPORT.v2022.1.Hs.grp | 2.02 | 0.00 | 3.01 | 2.64 | 0.00 | 4.00 | 1.62 | 0.01 | 2.22 |
| | GOBP_RESPIRATORY_ELECTRON_TRANSPORT_CHAIN.v2022.1.Hs.grp | 1.89 | 0.00 | 2.60 | 2.48 | 0.00 | 4.00 | 1.67 | 0.00 | 2.42 |
| | REACTOME_RESPIRATORY_ELECTRON_TRANSPORT_ATP_SYNTHESIS_BY_CHEMIOSMOTIC_COUPLING_AND_HEAT_PRODUCTION_BY_UNCOUPLING_PROTEINS.v2022.1.Hs.grp | 1.81 | 0.01 | 2.27 | 2.69 | 0.00 | 4.00 | 1.74 | 0.00 | 2.77 |
| | GOBP_ATP_BIOSYNTHETIC_PROCESS.v2022.1.Hs.grp | 1.76 | 0.01 | 2.05 | 2.08 | 0.00 | 3.19 | 1.66 | 0.00 | 2.38 |
| | Hs.grp | 1.54 | 0.04 | 1.44 | 2.31 | 0.00 | 4.00 | 1.60 | 0.01 | 2.16 |
| | GOBP_ELECTRON_TRANSPORT_CHAIN.v2022.1.Hs.grp | 1.59 | 0.03 | 1.52 | 2.30 | 0.00 | 4.00 | 1.72 | 0.00 | 2.69 |
| | REACTOME_RESPIRATORY_ELECTRON_TRANSPORT.v2022.1.Hs.grp | 1.83 | 0.00 | 2.36 | 2.53 | 0.00 | 4.00 | 1.62 | 0.01 | 2.22 |
| | KEGG_OXIDATIVE_PHOSPHORYLATION.v2022.1.Hs.grp | 1.85 | 0.00 | 2.41 | 2.31 | 0.00 | 4.00 | 1.69 | 0.00 | 2.57 |
| | REACTOME_COMPLEX_I_BIOGENESIS.v2022.1.Hs.grp | 1.83 | 0.00 | 2.37 | 2.13 | 0.00 | 3.23 | 1.58 | 0.01 | 2.10 |
| Myeloid Migration | GOBP_NEUTROPHIL_MIGRATION.v2022.1.Hs.grp | 2.02 | 0.00 | 3.03 | -0.94 | 0.81 | 0.09 | 1.97 | 0.00 | 4.00 |
| | GOBP_MYELOID_LEUKOCYTE_MIGRATION.v2022.1.Hs.grp | 1.99 | 0.00 | 2.98 | -0.95 | 0.83 | 0.08 | 1.84 | 0.00 | 3.20 |
| | GOBP_NEUTROPHIL_CHEMOTAXIS.v2022.1.Hs.grp | 1.99 | 0.00 | 3.00 | 0.88 | 0.72 | 0.14 | 2.12 | 0.00 | 4.00 |
| | GOBP_GRANULOCYTE_MIGRATION.v2022.1.Hs.grp | 1.97 | 0.00 | 2.99 | 1.08 | 0.42 | 0.38 | 1.94 | 0.00 | 4.00 |
| | GOBP_LEUKOCYTE_CHEMOTAXIS.v2022.1.Hs.grp | 1.90 | 0.00 | 2.61 | -0.94 | 0.79 | 0.10 | 1.73 | 0.00 | 2.68 |
| | GOBP_GRANULOCYTE_CHEMOTAXIS.v2022.1.Hs.grp | 1.90 | 0.00 | 2.59 | 1.06 | 0.43 | 0.37 | 2.07 | 0.00 | 4.00 |
| | GOBP_CELL_CHEMOTAXIS.v2022.1.Hs.grp | 1.89 | 0.00 | 2.61 | 1.05 | 0.44 | 0.35 | 1.72 | 0.00 | 2.68 |
| | GOBP_LEUKOCYTE_MIGRATION.v2022.1.Hs.grp | 1.81 | 0.01 | 2.26 | -1.04 | 0.76 | 0.12 | 1.68 | 0.00 | 2.45 |
| | GOBP_TAXIS.v2022.1.Hs.grp | 1.69 | 0.01 | 1.85 | 1.04 | 0.45 | 0.34 | 1.62 | 0.01 | 2.22 |
| Myeloid Activation | GOBP_GRANULOCYTE_ACTIVATION.v2022.1.Hs.grp | 1.47 | 0.06 | 1.26 | 1.16 | 0.31 | 0.50 | 1.52 | 0.02 | 1.79 |
| | GOBP_POSITIVE_REGULATION_OF_MYELOID_CELL_DIFFERENTIATION.v2022.1.Hs.grp | 1.53 | 0.04 | 1.39 | 1.03 | 0.47 | 0.33 | 1.49 | 0.02 | 1.69 |
| | GOBP_REGULATION_OF_MYELOID_LEUKOCYTE_MEDIATED_IMMUNITY.v2022.1.Hs.grp | 0.91 | 0.64 | 0.19 | -0.80 | 0.92 | 0.03 | 1.48 | 0.02 | 1.65 |
| | GOBP_LEUKOCYTE_MEDIATED_CYTOTOXICITY.v2022.1.Hs.grp | 1.29 | 0.15 | 0.82 | -0.85 | 0.89 | 0.05 | 1.88 | 0.00 | 3.52 |
| | GOBP_FC_RECEPTOR_SIGNALING_PATHWAY.v2022.1.Hs.grp | 1.12 | 0.33 | 0.48 | -0.99 | 0.80 | 0.09 | 1.39 | 0.05 | 1.30 |
| | GOBP_REGULATION_OF_LEUKOCYTE_MEDIATED_CYTOTOXICITY.v2022.1.Hs.grp | 1.16 | 0.28 | 0.55 | -1.03 | 0.73 | 0.14 | 1.72 | 0.00 | 2.66 |
| Inflammation | GOBP_RESPONSE_TO_CHEMOKINE.v2022.1.Hs.grp | 1.75 | 0.01 | 2.02 | -1.26 | 0.38 | 0.42 | 1.69 | 0.00 | 2.57 |
| | KEGG_CYTOKINE_CYTOKINE_RECEPTOR_INTERACTION.v2022.1.Hs.grp | 1.56 | 0.04 | 1.46 | -1.40 | 0.24 | 0.62 | 1.52 | 0.02 | 1.79 |
| | HALLMARK_IL6_JAK_STAT3_SIGNALING.v2022.1.Hs.grp | 1.55 | 0.04 | 1.45 | -1.48 | 0.17 | 0.77 | 1.48 | 0.02 | 1.63 |
| | GOBP_POSITIVE_REGULATION_OF_TUMOR_NECROSIS_FACTOR_SUPERFAMILY_CYTOKINE_PRODUCTION.v2022.1.Hs.grp | 1.57 | 0.03 | 1.47 | -1.26 | 0.37 | 0.44 | 1.59 | 0.01 | 2.12 |
| | GOBP_NEGATIVE_REGULATION_OF_VIRAL_GENOME_REPLICATION.v2022.1.Hs.grp | 1.93 | 0.00 | 2.72 | 1.53 | 0.06 | 1.20 | 1.35 | 0.07 | 1.17 |
| | GOBP_TUMOR_NECROSIS_FACTOR_SUPERFAMILY_CYTOKINE_PRODUCTION.v2022.1.Hs.grp | 1.81 | 0.01 | 2.26 | 0.97 | 0.56 | 0.25 | 1.56 | 0.01 | 1.98 |
| | HALLMARK_TNFA_SIGNALING_VIA_NFKB.v2022.1.Hs.grp | 1.61 | 0.03 | 1.58 | -2.35 | 0.00 | 4.00 | 1.44 | 0.03 | 1.49 |
| | REACTOME_INTERFERON_ALPHA_BETA_SIGNALING.v2022.1.Hs.grp | 1.96 | 0.00 | 2.88 | 1.87 | 0.01 | 2.14 | 1.58 | 0.01 | 2.09 |
| | HALLMARK_INTERFERON_ALPHA_RESPONSE.v2022.1.Hs.grp | 2.17 | 0.00 | 3.82 | 1.79 | 0.02 | 1.76 | 1.57 | 0.01 | 2.06 |
| | HALLMARK_INTERFERON_GAMMA_RESPONSE.v2022.1.Hs.grp | 2.04 | 0.00 | 3.09 | 0.92 | 0.64 | 0.19 | 1.47 | 0.03 | 1.60 |
| | GOBP_ACUTE_INFLAMMATORY_RESPONSE.v2022.1.Hs.grp | 2.02 | 0.00 | 2.99 | 1.41 | 0.12 | 0.93 | 1.76 | 0.00 | 2.79 |
| | HALLMARK_ALLOGRAFT_REJECTION.v2022.1.Hs.grp | 1.72 | 0.01 | 1.92 | -0.92 | 0.80 | 0.10 | 1.59 | 0.01 | 2.11 |
| | HALLMARK_INFLAMMATORY_RESPONSE.v2022.1.Hs.grp | 1.97 | 0.00 | 2.96 | -1.51 | 0.15 | 0.82 | 1.49 | 0.02 | 1.67 |

### Supplemental Table 4 Continued: GSEA on scRNAseq Populations, related to Figure 6
### NES=Normalized Enrichment Score, FDR=False Discovery Rate

| Group | MSigDB Signature Name | Combo vs GSK | | | | | | | | |
|---|---|---|---|---|---|---|---|---|---|---|
| | | Macs/Dend | | | Neus | | | Tumor | | |
| | | NES | FDR q | -Log(q) | NES | FDR q | -Log(q) | NES | FDR q | -Log(q) |
| DNA Replication and Damage | REACTOME_G1_S_DNA_DAMAGE_CHECKPOINTS.v2022.1.Hs.grp | 1.72 | 0.01 | 2.20 | 1.09 | 0.39 | | 1.30 | 0.12 | 0.92 |
| | REACTOME_ORC1_REMOVAL_FROM_CHROMATIN.v2022.1.Hs.grp | 1.76 | 0.00 | 2.38 | 1.29 | 0.20 | 0.69 | 1.27 | 0.14 | 0.84 |
| | REACTOME_SWITCHING_OF_ORIGINS_TO_A_POST_REPLICATIVE_STATE.v2022.1.Hs.grp | 1.65 | 0.01 | 1.96 | 1.20 | 0.26 | 0.58 | 1.23 | 0.18 | 0.75 |
| | REACTOME_CDT1_ASSOCIATION_WITH_THE_CDC6_ORC_ORIGIN_COMPLEX.v2022.1.Hs.grp | 1.76 | 0.00 | 2.37 | 1.30 | 0.20 | 0.69 | 1.21 | 0.19 | 0.71 |
| | REACTOME_SYNTHESIS_OF_DNA.v2022.1.Hs.grp | 1.62 | 0.01 | 1.84 | 1.18 | 0.29 | 0.54 | 1.20 | 0.20 | 0.69 |
| | REACTOME_APC_C_MEDIATED_DEGRADATION_OF_CELL_CYCLE_PROTEINS.v2022.1.Hs.grp | 1.66 | 0.01 | 2.00 | 1.22 | 0.25 | 0.60 | 1.20 | 0.20 | 0.69 |
| Protein and RNA Processing | KEGG_RIBOSOME.v2022.1.Hs.grp | 2.59 | 0.00 | 4.00 | -1.30 | 0.31 | 0.50 | -2.15 | 0.00 | 3.86 |
| | REACTOME_EUKARYOTIC_TRANSLATION_ELONGATION.v2022.1.Hs.grp | 2.61 | 0.00 | 4.00 | -1.43 | 0.28 | 0.55 | -2.28 | 0.00 | 4.00 |
| | REACTOME_SRP_DEPENDENT_COTRANSLATIONAL_PROTEIN_TARGETING_TO_MEMBRANE.v2022.1. | 2.51 | 0.00 | 4.00 | -1.25 | 0.33 | 0.48 | -2.12 | 0.00 | 3.70 |
| | REACTOME_RESPONSE_OF_EIF2AK4_GCN2_TO_AMINO_ACID_DEFICIENCY.v2022.1.Hs.grp | 2.63 | 0.00 | 4.00 | -1.40 | 0.33 | 0.49 | -2.27 | 0.00 | 4.00 |
| | REACTOME_EUKARYOTIC_TRANSLATION_INITIATION.v2022.1.Hs.grp | 2.52 | 0.00 | 4.00 | -1.38 | 0.27 | 0.57 | -2.09 | 0.00 | 3.48 |
| | REACTOME_INFLUENZA_INFECTION.v2022.1.Hs.grp | 2.53 | 0.00 | 4.00 | -1.40 | 0.30 | 0.52 | -2.18 | 0.00 | 3.76 |
| | GOBP_CYTOPLASMIC_TRANSLATION.v2022.1.Hs.grp | 2.41 | 0.00 | 4.00 | -1.46 | 0.25 | 0.60 | -2.18 | 0.00 | 3.81 |
| | REACTOME_TRANSLATION.v2022.1.Hs.grp | 2.29 | 0.00 | 4.00 | -1.29 | 0.33 | 0.48 | -2.02 | 0.00 | 3.36 |
| | REACTOME_SELENOAMINO_ACID_METABOLISM.v2022.1.Hs.grp | 2.56 | 0.00 | 4.00 | -1.36 | 0.29 | 0.54 | -2.24 | 0.00 | 4.00 |
| | REACTOME_CELLULAR_RESPONSE_TO_STARVATION.v2022.1.Hs.grp | 2.39 | 0.00 | 4.00 | -1.59 | 0.24 | 0.61 | -2.05 | 0.00 | 3.31 |
| | REACTOME_METABOLISM_OF_AMINO_ACIDS_AND_DERIVATIVES.v2022.1.Hs.grp | 2.21 | 0.00 | 4.00 | -1.09 | 0.54 | 0.27 | -1.66 | 0.01 | 1.85 |
| | REACTOME_NONSENSE_MEDIATED_DECAY_NMD.v2022.1.Hs.grp | 2.58 | 0.00 | 4.00 | -1.34 | 0.32 | 0.49 | -2.14 | 0.00 | 3.90 |
| | REACTOME_RRNA_PROCESSING.v2022.1.Hs.grp | 2.39 | 0.00 | 4.00 | -1.38 | 0.28 | 0.55 | -2.13 | 0.00 | 3.67 |
| Oxidative Phosphorylation | GOBP_AEROBIC_RESPIRATION.v2022.1.Hs.grp | 2.05 | 0.00 | 3.93 | 1.93 | 0.09 | 1.06 | 1.82 | 0.00 | 3.70 |
| | GOBP_OXIDATIVE_PHOSPHORYLATION.v2022.1.Hs.grp | 2.14 | 0.00 | 4.00 | 1.90 | 0.06 | 1.26 | 1.92 | 0.00 | 3.82 |
| | GOBP_ATP_SYNTHESIS_COUPLED_ELECTRON_TRANSPORT.v2022.1.Hs.grp | 2.14 | 0.00 | 4.00 | 1.83 | 0.06 | 1.23 | 1.88 | 0.00 | 3.97 |
| | GOBP_RESPIRATORY_ELECTRON_TRANSPORT_CHAIN.v2022.1.Hs.grp | 2.06 | 0.00 | 3.91 | 1.81 | 0.05 | 1.28 | 1.88 | 0.00 | 3.94 |
| | REACTOME_RESPIRATORY_ELECTRON_TRANSPORT_ATP_SYNTHESIS_BY_CHEMIOSMOTIC_COUPLING_AND_HEAT_PRODUCTION_BY_UNCOUPLING_PROTEINS.v2022.1.Hs.grp | 2.07 | 0.00 | 3.88 | 1.78 | 0.06 | 1.19 | 1.96 | 0.00 | 4.00 |
| | GOBP_ATP_BIOSYNTHETIC_PROCESS.v2022.1.Hs.grp | 2.08 | 0.00 | 3.85 | 1.72 | 0.06 | 1.24 | 1.98 | 0.00 | 4.00 |
| | REACTOME_THE_CITRIC_ACID_TCA_CYCLE_AND_RESPIRATORY_ELECTRON_TRANSPORT.v2022.1. | 1.96 | 0.00 | 3.45 | 1.72 | 0.05 | 1.26 | 1.87 | 0.00 | 4.04 |
| | GOBP_ELECTRON_TRANSPORT_CHAIN.v2022.1.Hs.grp | 1.84 | 0.00 | 2.70 | 1.71 | 0.05 | 1.26 | 1.88 | 0.00 | 4.19 |
| | REACTOME_RESPIRATORY_ELECTRON_TRANSPORT.v2022.1.Hs.grp | 1.95 | 0.00 | 3.41 | 1.70 | 0.05 | 1.30 | 1.91 | 0.00 | 4.04 |
| | KEGG_OXIDATIVE_PHOSPHORYLATION.v2022.1.Hs.grp | 2.07 | 0.00 | 3.87 | 1.65 | 0.06 | 1.21 | 2.05 | 0.00 | 4.00 |
| | REACTOME_COMPLEX_I_BIOGENESIS.v2022.1.Hs.grp | 1.85 | 0.00 | 2.77 | 1.58 | 0.08 | 1.09 | 1.80 | 0.00 | 3.38 |
| Myeloid Migration | GOBP_NEUTROPHIL_MIGRATION.v2022.1.Hs.grp | 1.81 | 0.00 | 2.59 | 1.43 | 0.14 | 0.84 | 1.83 | 0.00 | 3.72 |
| | GOBP_MYELOID_LEUKOCYTE_MIGRATION.v2022.1.Hs.grp | 1.88 | 0.00 | 2.95 | 1.38 | 0.17 | 0.78 | 1.84 | 0.00 | 3.72 |
| | GOBP_NEUTROPHIL_CHEMOTAXIS.v2022.1.Hs.grp | 1.84 | 0.00 | 2.70 | 1.26 | 0.21 | 0.68 | 1.75 | 0.00 | 3.07 |
| | GOBP_GRANULOCYTE_MIGRATION.v2022.1.Hs.grp | 1.89 | 0.00 | 2.99 | 1.54 | 0.09 | 1.03 | 1.75 | 0.00 | 3.07 |
| | GOBP_LEUKOCYTE_CHEMOTAXIS.v2022.1.Hs.grp | 1.91 | 0.00 | 3.14 | 1.35 | 0.17 | 0.76 | 1.51 | 0.02 | 1.72 |
| | GOBP_GRANULOCYTE_CHEMOTAXIS.v2022.1.Hs.grp | 1.90 | 0.00 | 3.05 | 1.39 | 0.16 | 0.79 | 1.71 | 0.00 | 2.84 |
| | GOBP_CELL_CHEMOTAXIS.v2022.1.Hs.grp | 1.98 | 0.00 | 3.60 | 1.58 | 0.08 | 1.11 | 1.64 | 0.00 | 2.41 |
| | GOBP_LEUKOCYTE_MIGRATION.v2022.1.Hs.grp | 1.80 | 0.00 | 2.54 | 1.43 | 0.14 | 0.84 | 1.68 | 0.00 | 2.68 |
| | GOBP_TAXIS.v2022.1.Hs.grp | 1.76 | 0.00 | 2.38 | 1.50 | 0.12 | 0.91 | 1.62 | 0.01 | 2.29 |
| Myeloid Activation | GOBP_GRANULOCYTE_ACTIVATION.v2022.1.Hs.grp | 1.57 | 0.02 | 1.66 | 1.55 | 0.09 | 1.03 | 1.73 | 0.00 | 3.01 |
| | GOBP_POSITIVE_REGULATION_OF_MYELOID_CELL_DIFFERENTIATION.v2022.1.Hs.grp | 1.29 | 0.15 | 0.83 | 1.08 | 0.39 | 0.41 | 1.76 | 0.00 | 3.16 |
| | GOBP_REGULATION_OF_MYELOID_LEUKOCYTE_MEDIATED_IMMUNITY.v2022.1.Hs.grp | 1.39 | 0.08 | 1.10 | 1.19 | 0.28 | 0.55 | 1.79 | 0.00 | 3.35 |
| | GOBP_LEUKOCYTE_MEDIATED_CYTOTOXICITY.v2022.1.Hs.grp | 1.35 | 0.10 | 1.00 | 0.91 | 0.65 | 0.19 | 1.72 | 0.00 | 2.95 |
| | GOBP_FC_RECEPTOR_SIGNALING_PATHWAY.v2022.1.Hs.grp | 1.39 | 0.08 | 1.10 | 0.94 | 0.60 | 0.22 | 1.77 | 0.00 | 3.19 |
| | GOBP_REGULATION_OF_LEUKOCYTE_MEDIATED_CYTOTOXICITY.v2022.1.Hs.grp | 1.12 | 0.31 | 0.50 | -0.69 | 0.99 | 0.00 | 1.79 | 0.00 | 3.39 |
| Inflammation | GOBP_RESPONSE_TO_CHEMOKINE.v2022.1.Hs.grp | 1.61 | 0.02 | 1.82 | -0.90 | 0.79 | 0.10 | 1.73 | 0.00 | 3.00 |
| | KEGG_CYTOKINE_CYTOKINE_RECEPTOR_INTERACTION.v2022.1.Hs.grp | 1.62 | 0.01 | 1.83 | 0.84 | 0.78 | 0.11 | 1.64 | 0.00 | 2.37 |
| | HALLMARK_IL6_JAK_STAT3_SIGNALING.v2022.1.Hs.grp | 1.67 | 0.01 | 2.02 | 0.66 | 0.96 | 0.02 | 1.50 | 0.02 | 1.66 |
| | GOBP_POSITIVE_REGULATION_OF_TUMOR_NECROSIS_FACTOR_SUPERFAMILY_CYTOKINE_PRODUCTION.v2022.1.Hs.grp | 1.24 | 0.18 | 0.75 | -1.12 | 0.48 | 0.32 | 1.89 | 0.00 | 4.15 |
| | GOBP_NEGATIVE_REGULATION_OF_VIRAL_GENOME_REPLICATION.v2022.1.Hs.grp | 1.52 | 0.03 | 1.51 | 1.66 | 0.06 | 1.26 | 1.83 | 0.00 | 3.70 |
| | GOBP_TUMOR_NECROSIS_FACTOR_SUPERFAMILY_CYTOKINE_PRODUCTION.v2022.1.Hs.grp | 1.38 | 0.08 | 1.09 | -0.87 | 0.83 | 0.08 | 1.77 | 0.00 | 3.17 |
| | HALLMARK_TNFA_SIGNALING_VIA_NFKB.v2022.1.Hs.grp | 1.65 | 0.01 | 1.96 | -1.33 | 0.32 | 0.49 | 1.38 | 0.06 | 1.20 |
| | REACTOME_INTERFERON_ALPHA_BETA_SIGNALING.v2022.1.Hs.grp | 1.67 | 0.01 | 2.01 | 1.12 | 0.35 | 0.46 | 1.73 | 0.00 | 3.01 |
| | HALLMARK_INTERFERON_ALPHA_RESPONSE.v2022.1.Hs.grp | 1.83 | 0.00 | 2.69 | 1.43 | 0.15 | 0.84 | 1.84 | 0.00 | 3.74 |
| | HALLMARK_INTERFERON_GAMMA_RESPONSE.v2022.1.Hs.grp | 1.98 | 0.00 | 3.57 | 1.30 | 0.20 | 0.70 | 1.78 | 0.00 | 3.28 |
| | GOBP_ACUTE_INFLAMMATORY_RESPONSE.v2022.1.Hs.grp | 1.85 | 0.00 | 2.74 | 1.68 | 0.06 | 1.24 | 1.71 | 0.00 | 2.83 |
| | HALLMARK_ALLOGRAFT_REJECTION.v2022.1.Hs.grp | 1.94 | 0.00 | 3.33 | 0.96 | 0.57 | 0.24 | 1.53 | 0.02 | 1.78 |
| | HALLMARK_INFLAMMATORY_RESPONSE.v2022.1.Hs.grp | 1.82 | 0.00 | 2.66 | 1.07 | 0.40 | 0.39 | 1.71 | 0.00 | 2.84 |

**Supplemental Table 5: GSEA on scRNAseq Populations, related to Figure 6**
**NES=Normalized Enrichment Score, FDR=False Discovery Rate**

| Group | MSigDB Signature Name | Combo vs Placebo | | | | | | | | |
|---|---|---|---|---|---|---|---|---|---|---|
| | | Macs/Dend | | | Neus | | | Tumor | | |
| | | NES | FDR c | -Log(q) | NES | FDR q | -Log(q) | NES | FDR q | -Log(q) |
| DNA Replication and Damage | REACTOME_G1_S_DNA_DAMAGE_CHECKPOINTS.v2022.1.Hs.grp | 2.19 | 0.00 | 4.00 | 1.86 | 0.00 | 2.76 | 1.50 | 0.03 | 1.60 |
| | REACTOME_ORC1_REMOVAL_FROM_CHROMATIN.v2022.1.Hs.grp | 2.27 | 0.00 | 4.00 | 1.95 | 0.00 | 3.16 | 1.44 | 0.04 | 1.39 |
| | REACTOME_SWITCHING_OF_ORIGINS_TO_A_POST_REPLICATIVE_STATE.v2022.1.Hs.grp | 2.18 | 0.00 | 4.00 | 1.95 | 0.00 | 3.16 | 1.42 | 0.05 | 1.33 |
| | REACTOME_CDT1_ASSOCIATION_WITH_THE_CDC6_ORC_ORIGIN_COMPLEX.v2022.1.Hs.grp | 2.22 | 0.00 | 4.00 | 1.88 | 0.00 | 2.86 | 1.47 | 0.03 | 1.52 |
| | REACTOME_SYNTHESIS_OF_DNA.v2022.1.Hs.grp | 2.06 | 0.00 | 4.01 | 2.01 | 0.00 | 3.50 | 1.39 | 0.06 | 1.23 |
| | REACTOME_APC_C_MEDIATED_DEGRADATION_OF_CELL_CYCLE_PROTEINS.v2022.1.Hs.grp | 2.18 | 0.00 | 4.00 | 1.96 | 0.00 | 3.18 | 1.47 | 0.03 | 1.52 |
| Protein and RNA Processing | KEGG_RIBOSOME.v2022.1.Hs.grp | 3.06 | 0.00 | 4.00 | 3.34 | 0.00 | 4.00 | 1.61 | 0.01 | 2.04 |
| | REACTOME_EUKARYOTIC_TRANSLATION_ELONGATION.v2022.1.Hs.grp | 3.06 | 0.00 | 4.00 | 3.33 | 0.00 | 4.00 | 1.57 | 0.01 | 1.90 |
| | REACTOME_SRP_DEPENDENT_COTRANSLATIONAL_PROTEIN_TARGETING_TO_MEMBRANE.v2022.1.Hs.grp | 3.03 | 0.00 | 4.00 | 3.34 | 0.00 | 4.00 | 1.58 | 0.01 | 1.92 |
| | REACTOME_RESPONSE_OF_EIF2AK4_GCN2_TO_AMINO_ACID_DEFICIENCY.v2022.1.Hs.grp | 3.06 | 0.00 | 4.00 | 3.32 | 0.00 | 4.00 | 1.52 | 0.02 | 1.68 |
| | REACTOME_EUKARYOTIC_TRANSLATION_INITIATION.v2022.1.Hs.grp | 2.94 | 0.00 | 4.00 | 3.36 | 0.00 | 4.00 | 1.58 | 0.01 | 1.92 |
| | REACTOME_INFLUENZA_INFECTION.v2022.1.Hs.grp | 2.89 | 0.00 | 4.00 | 3.23 | 0.00 | 4.00 | 1.33 | 0.09 | 1.02 |
| | GOBP_CYTOPLASMIC_TRANSLATION.v2022.1.Hs.grp | 2.81 | 0.00 | 4.00 | 3.26 | 0.00 | 4.00 | 1.48 | 0.03 | 1.53 |
| | REACTOME_TRANSLATION.v2022.1.Hs.grp | 2.81 | 0.00 | 4.00 | 3.21 | 0.00 | 4.00 | 1.35 | 0.08 | 1.11 |
| | REACTOME_SELENOAMINO_ACID_METABOLISM.v2022.1.Hs.grp | 3.04 | 0.00 | 4.00 | 3.36 | 0.00 | 4.00 | 1.50 | 0.02 | 1.61 |
| | REACTOME_CELLULAR_RESPONSE_TO_STARVATION.v2022.1.Hs.grp | 2.86 | 0.00 | 4.00 | 3.22 | 0.00 | 4.00 | 1.54 | 0.02 | 1.72 |
| | REACTOME_METABOLISM_OF_AMINO_ACIDS_AND_DERIVATIVES.v2022.1.Hs.grp | 2.72 | 0.00 | 4.00 | 3.06 | 0.00 | 4.00 | 1.41 | 0.05 | 1.32 |
| | REACTOME_NONSENSE_MEDIATED_DECAY_NMD.v2022.1.Hs.grp | 2.93 | 0.00 | 4.00 | 3.33 | 0.00 | 4.00 | 1.50 | 0.02 | 1.61 |
| | REACTOME_RRNA_PROCESSING.v2022.1.Hs.grp | 2.93 | 0.00 | 4.00 | 3.28 | 0.00 | 4.00 | 1.48 | 0.03 | 1.55 |
| Oxidative Phosphorylation | GOBP_AEROBIC_RESPIRATION.v2022.1.Hs.grp | 1.70 | 0.01 | 2.12 | 2.59 | 0.00 | 4.00 | 1.86 | 0.00 | 3.69 |
| | GOBP_OXIDATIVE_PHOSPHORYLATION.v2022.1.Hs.grp | 1.88 | 0.00 | 2.91 | 2.66 | 0.00 | 4.00 | 1.96 | 0.00 | 4.13 |
| | GOBP_ATP_SYNTHESIS_COUPLED_ELECTRON_TRANSPORT.v2022.1.Hs.grp | 1.83 | 0.00 | 2.73 | 2.54 | 0.00 | 4.00 | 1.92 | 0.00 | 3.99 |
| | GOBP_RESPIRATORY_ELECTRON_TRANSPORT_CHAIN.v2022.1.Hs.grp | 1.71 | 0.01 | 2.15 | 2.47 | 0.00 | 4.00 | 1.91 | 0.00 | 3.86 |
| | REACTOME_RESPIRATORY_ELECTRON_TRANSPORT_ATP_SYNTHESIS_BY_CHEMIOSMOTIC_COUPLING_AND_HEAT_PRODUCTION_BY_UNCOUPLING_PROTEINS.v2022.1.Hs.grp | 1.75 | 0.00 | 2.32 | 2.69 | 0.00 | 4.00 | 2.10 | 0.00 | 4.00 |
| | GOBP_ATP_BIOSYNTHETIC_PROCESS.v2022.1.Hs.grp | 1.84 | 0.00 | 2.73 | 2.45 | 0.00 | 4.00 | 2.06 | 0.00 | 4.00 |
| | REACTOME_THE_CITRIC_ACID_TCA_CYCLE_AND_RESPIRATORY_ELECTRON_TRANSPORT.v2022.1.Hs.grp | 1.48 | 0.04 | 1.43 | 2.64 | 0.00 | 4.00 | 1.92 | 0.00 | 3.97 |
| | GOBP_ELECTRON_TRANSPORT_CHAIN.v2022.1.Hs.grp | 1.46 | 0.04 | 1.36 | 2.27 | 0.00 | 4.00 | 1.90 | 0.00 | 3.77 |
| | REACTOME_RESPIRATORY_ELECTRON_TRANSPORT.v2022.1.Hs.grp | 1.70 | 0.01 | 2.13 | 2.53 | 0.00 | 4.00 | 2.01 | 0.00 | 3.79 |
| | KEGG_OXIDATIVE_PHOSPHORYLATION.v2022.1.Hs.grp | 1.69 | 0.01 | 2.12 | 2.63 | 0.00 | 4.00 | 2.13 | 0.00 | 4.00 |
| | REACTOME_COMPLEX_I_BIOGENESIS.v2022.1.Hs.grp | 1.82 | 0.00 | 2.63 | 2.20 | 0.00 | 4.00 | 1.90 | 0.00 | 3.75 |
| Myeloid Migration | GOBP_NEUTROPHIL_MIGRATION.v2022.1.Hs.grp | -1.43 | 0.06 | 1.25 | -1.49 | 0.08 | 1.11 | 1.16 | 0.27 | 0.57 |
| | GOBP_MYELOID_LEUKOCYTE_MIGRATION.v2022.1.Hs.grp | -1.51 | 0.04 | 1.43 | -1.43 | 0.10 | 1.00 | 1.30 | 0.11 | 0.94 |
| | GOBP_NEUTROPHIL_CHEMOTAXIS.v2022.1.Hs.grp | -1.41 | 0.07 | 1.18 | -1.50 | 0.07 | 1.13 | 1.20 | 0.22 | 0.66 |
| | GOBP_GRANULOCYTE_MIGRATION.v2022.1.Hs.grp | -1.49 | 0.04 | 1.40 | -1.43 | 0.10 | 0.99 | 1.23 | 0.18 | 0.74 |
| | GOBP_LEUKOCYTE_CHEMOTAXIS.v2022.1.Hs.grp | -1.50 | 0.04 | 1.43 | -1.76 | 0.01 | 1.84 | 1.09 | 0.37 | 0.43 |
| | GOBP_GRANULOCYTE_CHEMOTAXIS.v2022.1.Hs.grp | -1.50 | 0.04 | 1.43 | -1.46 | 0.09 | 1.06 | 1.25 | 0.17 | 0.78 |
| | GOBP_CELL_CHEMOTAXIS.v2022.1.Hs.grp | -1.50 | 0.04 | 1.43 | -1.84 | 0.01 | 2.03 | 1.19 | 0.23 | 0.64 |
| | GOBP_LEUKOCYTE_MIGRATION.v2022.1.Hs.grp | -1.60 | 0.02 | 1.64 | -1.78 | 0.01 | 1.89 | 1.13 | 0.31 | 0.51 |
| | GOBP_TAXIS.v2022.1.Hs.grp | -1.65 | 0.02 | 1.81 | -1.72 | 0.02 | 1.77 | 1.14 | 0.31 | 0.51 |
| Myeloid Activation | GOBP_GRANULOCYTE_ACTIVATION.v2022.1.Hs.grp | -1.19 | 0.22 | 0.65 | -1.21 | 0.24 | 0.62 | 1.04 | 0.47 | 0.33 |
| | GOBP_POSITIVE_REGULATION_OF_MYELOID_CELL_DIFFERENTIATION.v2022.1.Hs.grp | -1.44 | 0.05 | 1.28 | -1.80 | 0.01 | 1.91 | 0.75 | 0.88 | 0.06 |
| | GOBP_REGULATION_OF_MYELOID_LEUKOCYTE_MEDIATED_IMMUNITY.v2022.1.Hs.grp | -0.84 | 0.78 | 0.11 | -1.05 | 0.45 | 0.35 | -1.00 | 0.47 | 0.33 |
| | GOBP_LEUKOCYTE_MEDIATED_CYTOTOXICITY.v2022.1.Hs.grp | 1.23 | 0.16 | 0.79 | -1.33 | 0.16 | 0.81 | 1.42 | 0.05 | 1.33 |
| | GOBP_FC_RECEPTOR_SIGNALING_PATHWAY.v2022.1.Hs.grp | -0.91 | 0.66 | 0.18 | -1.51 | 0.07 | 1.14 | 0.67 | 0.93 | 0.03 |
| | GOBP_REGULATION_OF_LEUKOCYTE_MEDIATED_CYTOTOXICITY.v2022.1.Hs.grp | 1.22 | 0.17 | 0.77 | -1.58 | 0.05 | 1.32 | 0.97 | 0.58 | 0.23 |
| Inflammation | GOBP_RESPONSE_TO_CHEMOKINE.v2022.1.Hs.grp | -1.90 | 0.00 | 2.76 | -2.13 | 0.00 | 4.00 | -1.25 | 0.16 | 0.81 |
| | KEGG_CYTOKINE_CYTOKINE_RECEPTOR_INTERACTION.v2022.1.Hs.grp | -1.56 | 0.03 | 1.56 | -1.79 | 0.01 | 1.93 | -1.97 | 0.00 | 2.45 |
| | HALLMARK_IL6_JAK_STAT3_SIGNALING.v2022.1.Hs.grp | -1.55 | 0.03 | 1.53 | -1.97 | 0.00 | 2.37 | -1.15 | 0.24 | 0.61 |
| | p | 0.94 | 0.58 | 0.24 | -0.99 | 0.52 | 0.28 | 1.38 | 0.06 | 1.19 |
| | GOBP_NEGATIVE_REGULATION_OF_VIRAL_GENOME_REPLICATION.v2022.1.Hs.grp | 1.30 | 0.11 | 0.97 | 1.12 | 0.29 | 0.54 | -2.23 | 0.00 | 2.47 |
| | GOBP_TUMOR_NECROSIS_FACTOR_SUPERFAMILY_CYTOKINE_PRODUCTION.v2022.1.Hs.grp | -1.04 | 0.43 | 0.37 | -0.99 | 0.53 | 0.28 | 1.08 | 0.39 | 0.41 |
| | HALLMARK_TNFA_SIGNALING_VIA_NFKB.v2022.1.Hs.grp | -2.36 | 0.00 | 4.00 | -3.21 | 0.00 | 4.00 | -1.41 | 0.07 | 1.18 |
| | REACTOME_INTERFERON_ALPHA_BETA_SIGNALING.v2022.1.Hs.grp | 1.52 | 0.03 | 1.52 | 1.22 | 0.18 | 0.74 | -2.17 | 0.00 | 2.77 |
| | HALLMARK_INTERFERON_ALPHA_RESPONSE.v2022.1.Hs.grp | 1.44 | 0.05 | 1.34 | 1.39 | 0.08 | 1.11 | -2.14 | 0.00 | 2.93 |
| | HALLMARK_INTERFERON_GAMMA_RESPONSE.v2022.1.Hs.grp | 1.32 | 0.10 | 1.00 | -0.98 | 0.54 | 0.27 | -1.44 | 0.06 | 1.21 |
| | GOBP_ACUTE_INFLAMMATORY_RESPONSE.v2022.1.Hs.grp | 1.63 | 0.01 | 1.87 | -1.54 | 0.06 | 1.21 | 1.36 | 0.08 | 1.12 |
| | HALLMARK_ALLOGRAFT_REJECTION.v2022.1.Hs.grp | 1.36 | 0.08 | 1.11 | -1.03 | 0.47 | 0.33 | 1.06 | 0.43 | 0.36 |
| | HALLMARK_INFLAMMATORY_RESPONSE.v2022.1.Hs.grp | -1.55 | 0.03 | 1.52 | -1.90 | 0.01 | 2.28 | -1.47 | 0.06 | 1.22 |

### Supplemental Table 5 Continued: GSEA on scRNAseq Populations, related to Figure 6
### NES=Normalized Enrichment Score, FDR=False Discovery Rate

| Group | MSigDB Signature Name | GSK vs Placebo | | | | | | | | |
|---|---|---|---|---|---|---|---|---|---|---|
| | | Macs/Dend | | | Neus | | | Tumor | | |
| | | NES | FDR q | -Log(q) | NES | FDR q | -Log(q) | NES | FDR q | -Log(q) |
| DNA Replication and Damage | REACTOME_G1_S_DNA_DAMAGE_CHECKPOINTS.v2022.1.Hs.grp | 1.87 | 0.00 | 2.68 | 1.83 | 0.00 | 2.66 | 1.14 | 0.35 | 0.46 |
| | REACTOME_ORC1_REMOVAL_FROM_CHROMATIN.v2022.1.Hs.grp | 2.09 | 0.00 | 4.15 | 1.89 | 0.00 | 2.83 | 1.19 | 0.32 | 0.49 |
| | REACTOME_SWITCHING_OF_ORIGINS_TO_A_POST_REPLICATIVE_STATE.v2022.1.Hs.grp | 2.02 | 0.00 | 3.39 | 1.97 | 0.00 | 3.13 | 1.13 | 0.36 | 0.45 |
| | REACTOME_CDT1_ASSOCIATION_WITH_THE_CDC6_ORC_ORIGIN_COMPLEX.v2022.1.Hs.grp | 2.22 | 0.00 | 4.00 | 1.86 | 0.00 | 2.74 | 1.21 | 0.30 | 0.52 |
| | REACTOME_SYNTHESIS_OF_DNA.v2022.1.Hs.grp | 1.87 | 0.00 | 2.69 | 1.95 | 0.00 | 3.06 | 1.14 | 0.34 | 0.46 |
| | REACTOME_APC_C_MEDIATED_DEGRADATION_OF_CELL_CYCLE_PROTEINS.v2022.1.Hs.grp | 2.02 | 0.00 | 3.41 | 1.95 | 0.00 | 3.06 | 1.09 | 0.41 | 0.39 |
| Protein and RNA Processing | KEGG_RIBOSOME.v2022.1.Hs.grp | 2.45 | 0.00 | 4.00 | 3.46 | 0.00 | 4.00 | 2.46 | 0.00 | 4.00 |
| | REACTOME_EUKARYOTIC_TRANSLATION_ELONGATION.v2022.1.Hs.grp | 2.53 | 0.00 | 4.00 | 3.51 | 0.00 | 4.00 | 2.47 | 0.00 | 4.00 |
| | REACTOME_SRP_DEPENDENT_COTRANSLATIONAL_PROTEIN_TARGETING_TO_MEMBRANE.v2022.1.Hs.grp | 2.50 | 0.00 | 4.00 | 3.46 | 0.00 | 4.00 | 2.37 | 0.00 | 4.00 |
| | REACTOME_RESPONSE_OF_EIF2AK4_GCN2_TO_AMINO_ACID_DEFICIENCY.v2022.1.Hs.grp | 2.32 | 0.00 | 4.00 | 3.45 | 0.00 | 4.00 | 2.44 | 0.00 | 4.00 |
| | REACTOME_EUKARYOTIC_TRANSLATION_INITIATION.v2022.1.Hs.grp | 2.27 | 0.00 | 4.00 | 3.46 | 0.00 | 4.00 | 2.44 | 0.00 | 4.00 |
| | REACTOME_INFLUENZA_INFECTION.v2022.1.Hs.grp | 2.00 | 0.00 | 3.24 | 3.42 | 0.00 | 4.00 | 2.20 | 0.00 | 4.00 |
| | GOBP_CYTOPLASMIC_TRANSLATION.v2022.1.Hs.grp | 2.16 | 0.00 | 4.00 | 3.36 | 0.00 | 4.00 | 2.43 | 0.00 | 4.00 |
| | REACTOME_TRANSLATION.v2022.1.Hs.grp | 2.06 | 0.00 | 4.19 | 3.47 | 0.00 | 4.00 | 2.29 | 0.00 | 4.00 |
| | REACTOME_SELENOAMINO_ACID_METABOLISM.v2022.1.Hs.grp | 2.37 | 0.00 | 4.00 | 3.48 | 0.00 | 4.00 | 2.38 | 0.00 | 4.00 |
| | REACTOME_CELLULAR_RESPONSE_TO_STARVATION.v2022.1.Hs.grp | 2.31 | 0.00 | 4.00 | 3.38 | 0.00 | 4.00 | 2.38 | 0.00 | 4.00 |
| | REACTOME_METABOLISM_OF_AMINO_ACIDS_AND_DERIVATIVES.v2022.1.Hs.grp | 2.13 | 0.00 | 4.00 | 3.29 | 0.00 | 4.00 | 1.88 | 0.00 | 3.25 |
| | REACTOME_NONSENSE_MEDIATED_DECAY_NMD.v2022.1.Hs.grp | 2.21 | 0.00 | 4.00 | 3.47 | 0.00 | 4.00 | 2.35 | 0.00 | 4.00 |
| | REACTOME_RRNA_PROCESSING.v2022.1.Hs.grp | 2.18 | 0.00 | 4.00 | 3.41 | 0.00 | 4.00 | 2.37 | 0.00 | 4.00 |
| Oxidative Phosphorylation | GOBP_AEROBIC_RESPIRATION.v2022.1.Hs.grp | -1.23 | 0.21 | 0.67 | 1.94 | 0.00 | 3.03 | 1.14 | 0.35 | 0.46 |
| | GOBP_OXIDATIVE_PHOSPHORYLATION.v2022.1.Hs.grp | -1.13 | 0.32 | 0.49 | 2.05 | 0.00 | 3.80 | 1.25 | 0.27 | 0.58 |
| | GOBP_ATP_SYNTHESIS_COUPLED_ELECTRON_TRANSPORT.v2022.1.Hs.grp | -1.20 | 0.24 | 0.61 | 2.04 | 0.00 | 3.71 | 1.08 | 0.42 | 0.37 |
| | GOBP_RESPIRATORY_ELECTRON_TRANSPORT_CHAIN.v2022.1.Hs.grp | -1.09 | 0.36 | 0.44 | 2.00 | 0.00 | 3.42 | 1.18 | 0.33 | 0.48 |
| | REACTOME_RESPIRATORY_ELECTRON_TRANSPORT_ATP_SYNTHESIS_BY_CHEMIOSMOTIC_COUPLING_AND_HEAT_PRODUCTION_BY_UNCOUPLING_PROTEINS.v2022.1.Hs.grp | -1.34 | 0.13 | 0.90 | 2.14 | 0.00 | 4.10 | 1.31 | 0.20 | 0.70 |
| | GOBP_ATP_BIOSYNTHETIC_PROCESS.v2022.1.Hs.grp | -1.22 | 0.22 | 0.66 | 1.87 | 0.00 | 2.74 | 1.04 | 0.48 | 0.32 |
| | REACTOME_THE_CITRIC_ACID_TCA_CYCLE_AND_RESPIRATORY_ELECTRON_TRANSPORT.v2022.1.Hs.grp | -1.36 | 0.12 | 0.93 | 2.04 | 0.00 | 3.81 | 1.16 | 0.34 | 0.47 |
| | GOBP_ELECTRON_TRANSPORT_CHAIN.v2022.1.Hs.grp | -1.10 | 0.36 | 0.44 | 1.54 | 0.03 | 1.54 | 1.17 | 0.34 | 0.46 |
| | REACTOME_RESPIRATORY_ELECTRON_TRANSPORT.v2022.1.Hs.grp | -1.22 | 0.22 | 0.66 | 1.99 | 0.00 | 3.34 | 1.16 | 0.33 | 0.48 |
| | KEGG_OXIDATIVE_PHOSPHORYLATION.v2022.1.Hs.grp | -1.40 | 0.09 | 1.05 | 2.18 | 0.00 | 4.00 | 1.18 | 0.33 | 0.48 |
| | REACTOME_COMPLEX_I_BIOGENESIS.v2022.1.Hs.grp | -0.99 | 0.51 | 0.29 | 1.62 | 0.02 | 1.82 | 0.96 | 0.59 | 0.23 |
| Myeloid Migration | GOBP_NEUTROPHIL_MIGRATION.v2022.1.Hs.grp | -2.04 | 0.00 | 3.50 | -1.70 | 0.02 | 1.73 | -2.20 | 0.00 | 4.00 |
| | GOBP_MYELOID_LEUKOCYTE_MIGRATION.v2022.1.Hs.grp | -2.02 | 0.00 | 3.64 | -1.75 | 0.01 | 1.93 | -2.08 | 0.00 | 3.48 |
| | GOBP_NEUTROPHIL_CHEMOTAXIS.v2022.1.Hs.grp | -2.02 | 0.00 | 3.75 | -1.59 | 0.04 | 1.46 | -2.10 | 0.00 | 3.43 |
| | GOBP_GRANULOCYTE_MIGRATION.v2022.1.Hs.grp | -2.11 | 0.00 | 3.59 | -1.86 | 0.01 | 2.10 | -2.07 | 0.00 | 3.52 |
| | GOBP_LEUKOCYTE_CHEMOTAXIS.v2022.1.Hs.grp | -2.02 | 0.00 | 3.70 | -1.90 | 0.01 | 2.26 | -1.87 | 0.00 | 2.61 |
| | GOBP_GRANULOCYTE_CHEMOTAXIS.v2022.1.Hs.grp | -2.09 | 0.00 | 3.72 | -1.66 | 0.02 | 1.62 | -2.02 | 0.00 | 3.44 |
| | GOBP_CELL_CHEMOTAXIS.v2022.1.Hs.grp | -2.02 | 0.00 | 3.80 | -2.09 | 0.00 | 3.02 | -1.94 | 0.00 | 3.05 |
| | GOBP_LEUKOCYTE_MIGRATION.v2022.1.Hs.grp | -2.01 | 0.00 | 3.84 | -2.04 | 0.00 | 2.77 | -1.98 | 0.00 | 3.34 |
| | GOBP_TAXIS.v2022.1.Hs.grp | -1.98 | 0.00 | 3.76 | -2.04 | 0.00 | 2.74 | -1.77 | 0.01 | 2.24 |
| Myeloid Activation | GOBP_GRANULOCYTE_ACTIVATION.v2022.1.Hs.grp | -1.60 | 0.02 | 1.71 | -1.64 | 0.03 | 1.60 | -1.99 | 0.00 | 3.33 |
| | GOBP_POSITIVE_REGULATION_OF_MYELOID_CELL_DIFFERENTIATION.v2022.1.Hs.grp | -1.65 | 0.02 | 1.80 | -2.07 | 0.00 | 2.88 | -2.16 | 0.00 | 3.73 |
| | GOBP_REGULATION_OF_MYELOID_LEUKOCYTE_MEDIATED_IMMUNITY.v2022.1.Hs.grp | -1.26 | 0.19 | 0.71 | -1.34 | 0.13 | 0.90 | -2.04 | 0.00 | 3.54 |
| | GOBP_LEUKOCYTE_MEDIATED_CYTOTOXICITY.v2022.1.Hs.grp | -1.09 | 0.36 | 0.44 | -1.25 | 0.19 | 0.73 | -1.57 | 0.03 | 1.55 |
| | GOBP_FC_RECEPTOR_SIGNALING_PATHWAY.v2022.1.Hs.grp | -1.43 | 0.08 | 1.12 | -1.34 | 0.13 | 0.90 | | | |
| | GOBP_REGULATION_OF_LEUKOCYTE_MEDIATED_CYTOTOXICITY.v2022.1.Hs.grp | -0.93 | 0.62 | 0.21 | -1.18 | 0.24 | 0.61 | -1.84 | 0.00 | 2.54 |
| Inflammation | GOBP_RESPONSE_TO_CHEMOKINE.v2022.1.Hs.grp | -2.00 | 0.00 | 3.73 | -1.86 | 0.01 | 2.11 | -1.85 | 0.00 | 2.58 |
| | KEGG_CYTOKINE_CYTOKINE_RECEPTOR_INTERACTION.v2022.1.Hs.grp | -1.96 | 0.00 | 3.85 | -1.85 | 0.01 | 2.13 | -2.34 | 0.00 | 4.00 |
| | HALLMARK_IL6_JAK_STAT3_SIGNALING.v2022.1.Hs.grp | -2.01 | 0.00 | 3.88 | -1.98 | 0.00 | 2.58 | -2.16 | 0.00 | 3.76 |
| | p | -1.10 | 0.36 | 0.44 | -1.00 | 0.49 | 0.31 | -1.74 | 0.01 | 2.14 |
| | GOBP_NEGATIVE_REGULATION_OF_VIRAL_GENOME_REPLICATION.v2022.1.Hs.grp | -1.22 | 0.23 | 0.65 | -1.23 | 0.20 | 0.69 | -2.37 | 0.00 | 4.00 |
| | GOBP_TUMOR_NECROSIS_FACTOR_SUPERFAMILY_CYTOKINE_PRODUCTION.v2022.1.Hs.grp | -1.35 | 0.12 | 0.92 | -1.20 | 0.23 | 0.64 | -2.08 | 0.00 | 3.50 |
| | HALLMARK_TNFA_SIGNALING_VIA_NFKB.v2022.1.Hs.grp | -2.43 | 0.00 | 4.00 | -3.00 | 0.00 | 4.00 | -1.93 | 0.00 | 3.05 |
| | REACTOME_INTERFERON_ALPHA_BETA_SIGNALING.v2022.1.Hs.grp | -1.00 | 0.50 | 0.30 | 0.99 | 0.50 | 0.30 | -2.38 | 0.00 | 4.00 |
| | HALLMARK_INTERFERON_ALPHA_RESPONSE.v2022.1.Hs.grp | -1.17 | 0.27 | 0.57 | 1.05 | 0.40 | 0.40 | -2.66 | 0.00 | 4.00 |
| | HALLMARK_INTERFERON_GAMMA_RESPONSE.v2022.1.Hs.grp | -1.44 | 0.07 | 1.14 | -1.52 | 0.05 | 1.30 | -2.39 | 0.00 | 4.00 |
| | GOBP_ACUTE_INFLAMMATORY_RESPONSE.v2022.1.Hs.grp | -1.41 | 0.09 | 1.07 | -1.91 | 0.01 | 2.24 | -2.10 | 0.00 | 3.49 |
| | HALLMARK_ALLOGRAFT_REJECTION.v2022.1.Hs.grp | -1.61 | 0.02 | 1.70 | -1.39 | 0.10 | 1.00 | -1.43 | 0.06 | 1.20 |
| | HALLMARK_INFLAMMATORY_RESPONSE.v2022.1.Hs.grp | -2.12 | 0.00 | 3.42 | -2.32 | 0.00 | 4.00 | -2.23 | 0.00 | 4.00 |

**Supplemental Table 5 Continued: GSEA on scRNAseq Populations, related to Figure 6**
**NES=Normalized Enrichment Score, FDR=False Discovery Rate**

| Group | MSigDB Signature Name | aPD1 vs Placebo | | | | | | | | |
|---|---|---|---|---|---|---|---|---|---|---|
| | | Macs/Dend | | | Neus | | | Tumor | | |
| | | NES | FDR c | -Log(q) | NES | FDR q | -Log(q) | NES | FDR q | -Log(q) |
| DNA Replication and Damage | REACTOME_G1_S_DNA_DAMAGE_CHECKPOINTS.v2022.1.Hs.grp | 2.15 | 0.00 | 4.00 | 1.81 | 0.00 | 2.45 | 0.83 | 0.81 | 0.09 |
| | REACTOME_ORC1_REMOVAL_FROM_CHROMATIN.v2022.1.Hs.grp | 2.32 | 0.00 | 4.00 | 1.73 | 0.01 | 2.06 | 0.80 | 0.84 | 0.07 |
| | REACTOME_SWITCHING_OF_ORIGINS_TO_A_POST_REPLICATIVE_STATE.v2022.1.Hs.grp | 2.24 | 0.00 | 4.00 | 1.65 | 0.02 | 1.81 | 0.75 | 0.88 | 0.05 |
| | REACTOME_CDT1_ASSOCIATION_WITH_THE_CDC6_ORC_ORIGIN_COMPLEX.v2022.1.Hs.grp | 2.39 | 0.00 | 4.00 | 1.70 | 0.01 | 1.96 | 0.94 | 0.65 | 0.18 |
| | REACTOME_SYNTHESIS_OF_DNA.v2022.1.Hs.grp | 2.16 | 0.00 | 4.00 | 1.63 | 0.02 | 1.74 | 0.79 | 0.86 | 0.07 |
| | REACTOME_APC_C_MEDIATED_DEGRADATION_OF_CELL_CYCLE_PROTEINS.v2022.1.Hs.grp | 2.33 | 0.00 | 4.00 | 1.66 | 0.01 | 1.83 | 0.84 | 0.80 | 0.10 |
| Protein and RNA Processing | KEGG_RIBOSOME.v2022.1.Hs.grp | 3.25 | 0.00 | 4.00 | 3.40 | 0.00 | 4.00 | 2.48 | 0.00 | 4.00 |
| | REACTOME_EUKARYOTIC_TRANSLATION_ELONGATION.v2022.1.Hs.grp | 3.17 | 0.00 | 4.00 | 3.35 | 0.00 | 4.00 | 2.50 | 0.00 | 4.00 |
| | REACTOME_SRP_DEPENDENT_COTRANSLATIONAL_PROTEIN_TARGETING_TO_MEMBRANE.v2022.1.Hs.grp | 3.04 | 0.00 | 4.00 | 3.33 | 0.00 | 4.00 | 2.46 | 0.00 | 4.00 |
| | REACTOME_RESPONSE_OF_EIF2AK4_GCN2_TO_AMINO_ACID_DEFICIENCY.v2022.1.Hs.grp | 3.00 | 0.00 | 4.00 | 3.28 | 0.00 | 4.00 | 2.22 | 0.00 | 4.00 |
| | REACTOME_EUKARYOTIC_TRANSLATION_INITIATION.v2022.1.Hs.grp | 2.91 | 0.00 | 4.00 | 3.37 | 0.00 | 4.00 | 2.41 | 0.00 | 4.00 |
| | REACTOME_INFLUENZA_INFECTION.v2022.1.Hs.grp | 2.63 | 0.00 | 4.00 | 3.27 | 0.00 | 4.00 | 2.34 | 0.00 | 4.00 |
| | GOBP_CYTOPLASMIC_TRANSLATION.v2022.1.Hs.grp | 2.67 | 0.00 | 4.00 | 3.32 | 0.00 | 4.00 | 2.39 | 0.00 | 4.00 |
| | REACTOME_TRANSLATION.v2022.1.Hs.grp | 2.54 | 0.00 | 4.00 | 3.23 | 0.00 | 4.00 | 2.29 | 0.00 | 4.00 |
| | REACTOME_SELENOAMINO_ACID_METABOLISM.v2022.1.Hs.grp | 3.05 | 0.00 | 4.00 | 3.35 | 0.00 | 4.00 | 2.41 | 0.00 | 4.00 |
| | REACTOME_CELLULAR_RESPONSE_TO_STARVATION.v2022.1.Hs.grp | 2.85 | 0.00 | 4.00 | 3.25 | 0.00 | 4.00 | 2.17 | 0.00 | 4.00 |
| | REACTOME_METABOLISM_OF_AMINO_ACIDS_AND_DERIVATIVES.v2022.1.Hs.grp | 2.58 | 0.00 | 4.00 | 3.16 | 0.00 | 4.00 | 2.00 | 0.00 | 4.00 |
| | REACTOME_NONSENSE_MEDIATED_DECAY_NMD.v2022.1.Hs.grp | 2.90 | 0.00 | 4.00 | 3.38 | 0.00 | 4.00 | 2.42 | 0.00 | 4.00 |
| | REACTOME_RRNA_PROCESSING.v2022.1.Hs.grp | 2.62 | 0.00 | 4.00 | 3.29 | 0.00 | 4.00 | 2.38 | 0.00 | 4.00 |
| Oxidative Phosphorylation | GOBP_AEROBIC_RESPIRATION.v2022.1.Hs.grp | -1.04 | 0.47 | 0.33 | 0.98 | 0.58 | 0.24 | 1.64 | 0.01 | 1.96 |
| | GOBP_OXIDATIVE_PHOSPHORYLATION.v2022.1.Hs.grp | -1.05 | 0.45 | 0.34 | 0.96 | 0.61 | 0.21 | 1.71 | 0.00 | 2.32 |
| | GOBP_ATP_SYNTHESIS_COUPLED_ELECTRON_TRANSPORT.v2022.1.Hs.grp | -1.14 | 0.32 | 0.49 | 0.87 | 0.75 | 0.12 | 1.62 | 0.01 | 1.89 |
| | GOBP_RESPIRATORY_ELECTRON_TRANSPORT_CHAIN.v2022.1.Hs.grp | -1.15 | 0.32 | 0.50 | 1.03 | 0.50 | 0.30 | 1.61 | 0.01 | 1.85 |
| | REACTOME_RESPIRATORY_ELECTRON_TRANSPORT_ATP_SYNTHESIS_BY_CHEMIOSMOTIC_COUPLING_AND_HEAT_PRODUCTION_BY_UNCOUPLING_PROTEINS.v2022.1.Hs.grp | -1.11 | 0.36 | 0.44 | 0.93 | 0.66 | 0.18 | 1.77 | 0.00 | 2.64 |
| | GOBP_ATP_BIOSYNTHETIC_PROCESS.v2022.1.Hs.grp | -0.98 | 0.55 | 0.26 | 1.12 | 0.38 | 0.42 | 1.73 | 0.00 | 2.44 |
| | REACTOME_THE_CITRIC_ACID_TCA_CYCLE_AND_RESPIRATORY_ELECTRON_TRANSPORT.v2022.1.Hs.grp | -1.19 | 0.27 | 0.57 | 1.08 | 0.43 | 0.36 | 1.69 | 0.01 | 2.18 |
| | GOBP_ELECTRON_TRANSPORT_CHAIN.v2022.1.Hs.grp | -1.17 | 0.29 | 0.54 | 0.74 | 0.93 | 0.03 | 1.57 | 0.02 | 1.64 |
| | REACTOME_RESPIRATORY_ELECTRON_TRANSPORT.v2022.1.Hs.grp | -1.14 | 0.32 | 0.50 | 0.75 | 0.92 | 0.04 | 1.65 | 0.01 | 1.97 |
| | KEGG_OXIDATIVE_PHOSPHORYLATION.v2022.1.Hs.grp | -1.26 | 0.19 | 0.72 | 1.39 | 0.09 | 1.03 | 1.75 | 0.00 | 2.56 |
| | REACTOME_COMPLEX_I_BIOGENESIS.v2022.1.Hs.grp | -1.04 | 0.46 | 0.34 | 0.64 | 0.98 | 0.01 | 1.38 | 0.10 | 1.00 |
| Myeloid Migration | GOBP_NEUTROPHIL_MIGRATION.v2022.1.Hs.grp | -2.07 | 0.00 | 4.00 | -1.38 | 0.13 | 0.89 | -2.21 | 0.00 | 4.00 |
| | GOBP_MYELOID_LEUKOCYTE_MIGRATION.v2022.1.Hs.grp | -2.11 | 0.00 | 4.00 | -1.61 | 0.05 | 1.28 | -1.91 | 0.00 | 2.81 |
| | GOBP_NEUTROPHIL_CHEMOTAXIS.v2022.1.Hs.grp | -2.06 | 0.00 | 4.00 | -1.45 | 0.10 | 1.00 | -2.14 | 0.00 | 4.00 |
| | GOBP_GRANULOCYTE_MIGRATION.v2022.1.Hs.grp | -2.16 | 0.00 | 4.00 | -1.62 | 0.05 | 1.27 | -2.08 | 0.00 | 3.99 |
| | GOBP_LEUKOCYTE_CHEMOTAXIS.v2022.1.Hs.grp | -1.95 | 0.00 | 3.67 | -1.86 | 0.01 | 1.85 | -1.98 | 0.00 | 3.28 |
| | GOBP_GRANULOCYTE_CHEMOTAXIS.v2022.1.Hs.grp | -2.15 | 0.00 | 4.00 | -1.73 | 0.03 | 1.55 | -2.01 | 0.00 | 3.44 |
| | GOBP_CELL_CHEMOTAXIS.v2022.1.Hs.grp | -1.91 | 0.00 | 3.35 | -2.09 | 0.00 | 2.62 | -1.83 | 0.00 | 2.51 |
| | GOBP_LEUKOCYTE_MIGRATION.v2022.1.Hs.grp | -1.97 | 0.00 | 3.86 | -1.91 | 0.01 | 1.99 | -1.81 | 0.00 | 2.38 |
| | GOBP_TAXIS.v2022.1.Hs.grp | -1.87 | 0.00 | 3.06 | -1.96 | 0.01 | 2.07 | -1.75 | 0.01 | 2.18 |
| Myeloid Activation | GOBP_GRANULOCYTE_ACTIVATION.v2022.1.Hs.grp | -1.56 | 0.03 | 1.55 | -1.50 | 0.09 | 1.06 | -1.51 | 0.04 | 1.39 |
| | GOBP_POSITIVE_REGULATION_OF_MYELOID_CELL_DIFFERENTIATION.v2022.1.Hs.grp | -1.67 | 0.01 | 2.02 | -2.28 | 0.00 | 3.23 | -1.78 | 0.00 | 2.31 |
| | GOBP_REGULATION_OF_MYELOID_LEUKOCYTE_MEDIATED_IMMUNITY.v2022.1.Hs.grp | -0.87 | 0.72 | 0.15 | 0.69 | 0.96 | 0.02 | -1.87 | 0.00 | 2.65 |
| | GOBP_LEUKOCYTE_MEDIATED_CYTOTOXICITY.v2022.1.Hs.grp | -0.84 | 0.75 | 0.13 | -1.13 | 0.32 | 0.49 | -1.68 | 0.01 | 1.92 |
| | GOBP_FC_RECEPTOR_SIGNALING_PATHWAY.v2022.1.Hs.grp | -0.95 | 0.60 | 0.22 | | | | -1.27 | 0.15 | 0.83 |
| | GOBP_REGULATION_OF_LEUKOCYTE_MEDIATED_CYTOTOXICITY.v2022.1.Hs.grp | 1.01 | 0.42 | 0.38 | -1.17 | 0.30 | 0.52 | -1.87 | 0.00 | 2.66 |
| Inflammation | GOBP_RESPONSE_TO_CHEMOKINE.v2022.1.Hs.grp | -2.07 | 0.00 | 4.00 | -1.68 | 0.03 | 1.46 | -1.80 | 0.00 | 2.31 |
| | KEGG_CYTOKINE_CYTOKINE_RECEPTOR_INTERACTION.v2022.1.Hs.grp | -1.80 | 0.00 | 2.60 | -1.69 | 0.03 | 1.48 | -2.23 | 0.00 | 4.00 |
| | HALLMARK_IL6_JAK_STAT3_SIGNALING.v2022.1.Hs.grp | -1.72 | 0.01 | 2.23 | -1.56 | 0.06 | 1.19 | -1.91 | 0.00 | 2.80 |
| | p | -1.15 | 0.30 | 0.52 | 0.73 | 0.93 | 0.03 | -1.34 | 0.10 | 1.00 |
| | GOBP_NEGATIVE_REGULATION_OF_VIRAL_GENOME_REPLICATION.v2022.1.Hs.grp | -1.48 | 0.06 | 1.24 | -1.60 | 0.05 | 1.28 | -2.10 | 0.00 | 4.00 |
| | GOBP_TUMOR_NECROSIS_FACTOR_SUPERFAMILY_CYTOKINE_PRODUCTION.v2022.1.Hs.grp | -1.41 | 0.08 | 1.07 | -1.08 | 0.38 | 0.42 | -1.48 | 0.05 | 1.32 |
| | HALLMARK_TNFA_SIGNALING_VIA_NFKB.v2022.1.Hs.grp | -2.32 | 0.00 | 4.00 | -2.33 | 0.00 | 2.93 | -1.87 | 0.00 | 2.63 |
| | REACTOME_INTERFERON_ALPHA_BETA_SIGNALING.v2022.1.Hs.grp | -1.29 | 0.16 | 0.79 | -1.79 | 0.02 | 1.70 | -2.28 | 0.00 | 4.00 |
| | HALLMARK_INTERFERON_ALPHA_RESPONSE.v2022.1.Hs.grp | -1.39 | 0.09 | 1.04 | -1.15 | 0.32 | 0.49 | -2.46 | 0.00 | 4.00 |
| | HALLMARK_INTERFERON_GAMMA_RESPONSE.v2022.1.Hs.grp | -1.43 | 0.08 | 1.12 | -1.04 | 0.41 | 0.38 | -2.18 | 0.00 | 4.00 |
| | GOBP_ACUTE_INFLAMMATORY_RESPONSE.v2022.1.Hs.grp | -1.37 | 0.11 | 0.98 | -1.90 | 0.01 | 2.01 | -1.77 | 0.01 | 2.24 |
| | HALLMARK_ALLOGRAFT_REJECTION.v2022.1.Hs.grp | -1.40 | 0.09 | 1.06 | -1.13 | 0.32 | 0.49 | -1.62 | 0.02 | 1.75 |
| | HALLMARK_INFLAMMATORY_RESPONSE.v2022.1.Hs.grp | -2.14 | 0.00 | 4.00 | -1.61 | 0.05 | 1.29 | -1.88 | 0.00 | 2.65 |

**Supplemental Table 6: Genes Highly Expressed in Neutrophil Clusters, Related to Figure 6**
**Log2FC=Log2-fold change between cluster and all others, pct=percentage of cells expressing**

| Cluster | Gene Symbol | Log2FC | pct.1 | pct.2 | P value | Adj P value | Cluster | Gene Symbol | Log2FC | pct.1 | pct.2 | P value | Adj P value |
|---|---|---|---|---|---|---|---|---|---|---|---|---|---|
| Neu-1 | Cst3 | 2.050 | 0.929 | 0.784 | 0.0E+00 | 0.0E+00 | Neu-3 | Gadd45b | 2.421 | 0.877 | 0.664 | 0.0E+00 | 0.0E+00 |
| | Gngt2 | 1.920 | 0.812 | 0.557 | 0.0E+00 | 0.0E+00 | | Nceh1 | 2.367 | 0.856 | 0.760 | 0.0E+00 | 0.0E+00 |
| | Hexb | 1.393 | 0.794 | 0.754 | 0.0E+00 | 0.0E+00 | | Gpnmb | 2.274 | 0.781 | 0.565 | 0.0E+00 | 0.0E+00 |
| | Gpx1 | 1.277 | 0.869 | 0.823 | 0.0E+00 | 0.0E+00 | | Psap | 2.244 | 0.852 | 0.718 | 0.0E+00 | 0.0E+00 |
| | Ptgs1 | 1.126 | 0.772 | 0.702 | 0.0E+00 | 0.0E+00 | | Ifrd1 | 2.039 | 0.819 | 0.707 | 0.0E+00 | 0.0E+00 |
| | Ms4a6d | 0.881 | 0.805 | 0.780 | 0.0E+00 | 0.0E+00 | | Hilpda | 2.038 | 0.836 | 0.799 | 0.0E+00 | 0.0E+00 |
| | Gm19951 | 0.635 | 0.840 | 0.837 | 0.0E+00 | 0.0E+00 | | Ccl3 | 1.982 | 0.920 | 0.759 | 0.0E+00 | 0.0E+00 |
| | Cenpx | 0.430 | 0.745 | 0.593 | 0.0E+00 | 0.0E+00 | | Atp6v1c1 | 1.914 | 0.857 | 0.766 | 0.0E+00 | 0.0E+00 |
| | Hist1h4d | 0.351 | 0.793 | 0.747 | 0.0E+00 | 0.0E+00 | | Lamp1 | 1.832 | 0.885 | 0.784 | 0.0E+00 | 0.0E+00 |
| | Ranbp1 | 0.333 | 0.704 | 0.546 | 0.0E+00 | 0.0E+00 | | Hcar2 | 1.829 | 0.884 | 0.824 | 0.0E+00 | 0.0E+00 |
| | Card11 | 0.298 | 0.831 | 0.810 | 0.0E+00 | 0.0E+00 | | Ctsb | 1.789 | 0.977 | 0.916 | 0.0E+00 | 0.0E+00 |
| | Hist1h1d | 0.297 | 0.697 | 0.506 | 0.0E+00 | 0.0E+00 | | Cd63 | 1.786 | 0.936 | 0.864 | 0.0E+00 | 0.0E+00 |
| | Dmkn | 0.263 | 0.794 | 0.646 | 0.0E+00 | 0.0E+00 | | Zeb2 | 1.742 | 0.829 | 0.553 | 0.0E+00 | 0.0E+00 |
| | Cpt1a | 0.256 | 0.724 | 0.482 | 0.0E+00 | 0.0E+00 | | Ctsz | 1.678 | 0.936 | 0.850 | 0.0E+00 | 0.0E+00 |
| | H1f0 | 0.809 | 0.700 | 0.554 | 4.0E-279 | 5.9E-276 | | Ftl1 | 1.590 | 1.000 | 0.999 | 0.0E+00 | 0.0E+00 |
| | Laptm5 | 0.807 | 0.829 | 0.784 | 8.0E-255 | 1.2E-251 | | F10 | 1.513 | 0.863 | 0.864 | 0.0E+00 | 0.0E+00 |
| | Asah1 | 0.427 | 0.666 | 0.444 | 2.5E-208 | 3.7E-205 | | Gas2l3 | 1.194 | 0.818 | 0.737 | 0.0E+00 | 0.0E+00 |
| | Atp1a1 | 1.115 | 0.779 | 0.714 | 3.2E-204 | 4.8E-201 | | Dock10 | 1.193 | 0.783 | 0.789 | 0.0E+00 | 0.0E+00 |
| | Pmaip1 | 0.631 | 0.787 | 0.697 | 2.3E-202 | 3.5E-199 | | Lhfpl2 | 1.059 | 0.784 | 0.606 | 0.0E+00 | 0.0E+00 |
| | Hist1h1e | 0.525 | 0.794 | 0.723 | 2.0E-201 | 2.9E-198 | | P2rx7 | 0.743 | 0.760 | 0.628 | 0.0E+00 | 0.0E+00 |
| | Atp5g1 | 0.535 | 0.667 | 0.503 | 3.9E-197 | 5.8E-194 | | Pdxk | 0.707 | 0.726 | 0.490 | 0.0E+00 | 0.0E+00 |
| | Ltc4s | 0.855 | 0.644 | 0.562 | 2.4E-193 | 3.5E-190 | | Gstm1 | 0.948 | 0.697 | 0.537 | 5.0E-285 | 7.5E-282 |
| | Ptma | 1.199 | 0.760 | 0.667 | 4.4E-184 | 6.5E-181 | | Tst | 0.523 | 0.688 | 0.501 | 9.7E-282 | 1.5E-278 |
| | Reep5 | 0.947 | 0.749 | 0.636 | 1.4E-175 | 2.1E-172 | | Hexa | 1.286 | 0.839 | 0.735 | 4.4E-268 | 6.5E-265 |
| | Fam96a | 0.381 | 0.589 | 0.440 | 2.4E-164 | 3.7E-161 | | Atp6v0d2 | 0.978 | 0.752 | 0.684 | 3.5E-267 | 5.3E-264 |
| | Cd81 | 0.475 | 0.278 | 0.442 | 2.2E-155 | 3.3E-152 | | Dhfr | 0.859 | 0.789 | 0.859 | 3.2E-265 | 4.7E-262 |
| | Rps2 | 0.817 | 0.873 | 0.819 | 8.0E-152 | 1.2E-148 | | Chka | 0.492 | 0.738 | 0.529 | 2.2E-261 | 3.2E-258 |
| | Chil3 | 1.278 | 0.842 | 0.760 | 4.3E-130 | 6.4E-127 | | Plcxd2 | 0.704 | 0.728 | 0.788 | 1.7E-250 | 2.5E-247 |
| | Agap1 | 0.711 | 0.586 | 0.439 | 4.9E-117 | 7.3E-114 | | Fcgr2b | 0.662 | 0.838 | 0.714 | 5.9E-241 | 8.8E-238 |
| | Cd302 | 0.256 | 0.296 | 0.474 | 2.5E-116 | 3.8E-113 | | Gns | 1.455 | 0.782 | 0.705 | 7.0E-240 | 1.1E-236 |
| | Cd300c2 | 0.778 | 0.716 | 0.616 | 2.5E-111 | 3.8E-108 | | Ctsd | 0.702 | 0.987 | 0.976 | 1.2E-220 | 1.8E-217 |
| | Lrp1 | 0.630 | 0.529 | 0.374 | 2.7E-108 | 4.0E-105 | | Atf3 | 1.518 | 0.679 | 0.486 | 1.4E-214 | 2.2E-211 |
| | Krt19 | 0.266 | 0.659 | 0.814 | 1.1E-103 | 1.7E-100 | | Npc1 | 1.184 | 0.749 | 0.753 | 6.1E-211 | 9.1E-208 |
| | Hebp1 | 0.441 | 0.646 | 0.524 | 5.1E-103 | 7.6E-100 | | Aprt | 1.030 | 0.820 | 0.695 | 1.5E-210 | 2.3E-207 |
| | Ccnd2 | 0.297 | 0.487 | 0.674 | 3.5E-88 | 5.2E-85 | | Cd68 | 1.480 | 0.714 | 0.531 | 1.7E-208 | 2.6E-205 |
| | Ssbp4 | 0.336 | 0.596 | 0.543 | 6.5E-87 | 9.8E-84 | | Canx | 0.895 | 0.769 | 0.640 | 1.5E-207 | 2.3E-204 |
| | Tcf4 | 0.262 | 0.442 | 0.288 | 1.4E-86 | 2.1E-83 | | Cd274 | 1.443 | 0.725 | 0.671 | 2.9E-205 | 4.3E-202 |
| | Dpep2 | 0.452 | 0.652 | 0.764 | 1.9E-86 | 2.8E-83 | | Plekhm2 | 1.059 | 0.785 | 0.766 | 4.3E-197 | 6.4E-194 |
| | Grn | 0.508 | 0.869 | 0.830 | 2.0E-84 | 2.9E-81 | | Prdx1 | 1.174 | 0.673 | 0.462 | 6.6E-191 | 9.9E-188 |
| | Calr | 0.613 | 0.710 | 0.613 | 1.3E-82 | 1.9E-79 | | Tpp1 | 1.030 | 0.716 | 0.620 | 1.2E-185 | 1.8E-182 |
| | B930036N10Ri | 0.429 | 0.642 | 0.682 | 1.8E-82 | 2.7E-79 | | Naglu | 0.956 | 0.616 | 0.469 | 3.1E-165 | 4.6E-162 |
| | Naaa | 0.466 | 0.655 | 0.600 | 1.6E-81 | 2.4E-78 | | H2-Eb1 | 0.929 | 0.683 | 0.729 | 7.5E-150 | 1.1E-146 |
| | Pgls | 0.322 | 0.607 | 0.512 | 2.7E-81 | 4.0E-78 | | Dhrs3 | 0.764 | 0.674 | 0.525 | 6.6E-145 | 9.9E-142 |
| | Id2 | 0.400 | 0.925 | 0.882 | 1.7E-75 | 2.6E-72 | | Dnmt1 | 0.341 | 0.626 | 0.526 | 3.8E-137 | 5.7E-134 |
| | Bcl2a1b | 0.796 | 0.640 | 0.563 | 8.2E-75 | 1.2E-71 | | Npc2 | 1.027 | 0.875 | 0.886 | 1.8E-132 | 2.7E-129 |
| | Ctsa | 0.531 | 0.744 | 0.686 | 2.7E-68 | 4.0E-65 | | Tmem86a | 0.556 | 0.724 | 0.748 | 2.4E-128 | 3.5E-125 |
| | Manf | 0.354 | 0.814 | 0.794 | 6.8E-68 | 1.0E-64 | | Gadd45g | 1.393 | 0.634 | 0.509 | 5.6E-122 | 8.4E-119 |
| | Cybb | 0.303 | 0.519 | 0.409 | 4.6E-65 | 6.8E-62 | | Id2 | 0.969 | 0.874 | 0.904 | 4.8E-121 | 7.2E-118 |
| | Gm26917 | 0.282 | 0.585 | 0.474 | 1.5E-61 | 2.2E-58 | | Hpgds | 0.522 | 0.680 | 0.724 | 9.2E-119 | 1.4E-115 |
| | Ndufa4 | 0.667 | 0.623 | 0.564 | 1.9E-60 | 2.9E-57 | | Inhba | 0.874 | 0.622 | 0.511 | 6.6E-114 | 9.9E-111 |
| | Hist1h1c | 0.752 | 0.664 | 0.572 | 3.1E-58 | 4.7E-55 | | Tcirg1 | 1.016 | 0.670 | 0.594 | 1.6E-112 | 2.5E-109 |
| | Bcl2a1a | 0.768 | 0.592 | 0.516 | 3.8E-58 | 5.7E-55 | | Lgmn | 0.951 | 0.627 | 0.518 | 1.4E-111 | 2.1E-108 |
| | Gm2a | 0.892 | 0.637 | 0.616 | 1.3E-56 | 2.0E-53 | | Ccnf | 0.540 | 0.662 | 0.781 | 5.7E-109 | 8.6E-106 |
| | Phlda1 | 0.268 | 0.391 | 0.530 | 1.3E-55 | 1.9E-52 | | Creg1 | 1.377 | 0.702 | 0.577 | 1.2E-106 | 1.9E-103 |
| | Mt1 | 0.286 | 0.750 | 0.734 | 4.5E-55 | 6.8E-52 | | Myo5a | 0.533 | 0.480 | 0.319 | 2.3E-99 | 3.5E-96 |
| | Hist1h4i | 0.880 | 0.646 | 0.592 | 5.1E-54 | 7.6E-51 | | Mpeg1 | 0.803 | 0.701 | 0.585 | 5.7E-99 | 8.5E-96 |

| | | | | | | | | | | | | | |
|---|---|---|---|---|---|---|---|---|---|---|---|---|---|
| | Nap1l1 | 0.311 | 0.514 | 0.434 | 7.4E-52 | 1.1E-48 | | Ccl4 | 1.163 | 0.868 | 0.935 | 3.7E-95 | 5.6E-92 |
| | Ccng1 | 0.519 | 0.520 | 0.461 | 1.5E-51 | 2.2E-48 | | Syngr1 | 0.793 | 0.721 | 0.791 | 7.6E-92 | 1.1E-88 |
| | Unc93b1 | 0.566 | 0.570 | 0.493 | 1.8E-51 | 2.6E-48 | | Aplp2 | 0.334 | 0.718 | 0.588 | 7.3E-91 | 1.1E-87 |
| | Pycard | 0.298 | 0.870 | 0.846 | 2.1E-51 | 3.1E-48 | | Hmox1 | 1.205 | 0.790 | 0.778 | 9.1E-85 | 1.4E-81 |
| | P2ry6 | 0.379 | 0.710 | 0.721 | 1.9E-50 | 2.9E-47 | | Hspa9 | 0.589 | 0.651 | 0.663 | 2.2E-81 | 3.4E-78 |
| | Tubb5 | 0.364 | 0.356 | 0.505 | 2.0E-50 | 3.0E-47 | | Hspa5 | 0.742 | 0.800 | 0.740 | 2.0E-80 | 2.9E-77 |
| | Tubb4b | 0.403 | 0.662 | 0.575 | 5.7E-48 | 8.6E-45 | | Dpp7 | 0.502 | 0.624 | 0.692 | 1.0E-72 | 1.5E-69 |
| | Pdia6 | 0.555 | 0.775 | 0.771 | 7.2E-48 | 1.1E-44 | | Hsp90b1 | 0.441 | 0.736 | 0.690 | 3.7E-72 | 5.5E-69 |
| | Gm20186 | 0.390 | 0.404 | 0.535 | 4.5E-47 | 6.7E-44 | | Rgs1 | 0.942 | 0.774 | 0.774 | 7.5E-70 | 1.1E-66 |
| | Ucp2 | 0.715 | 0.743 | 0.739 | 8.8E-42 | 1.3E-38 | | Hk2 | 0.381 | 0.466 | 0.660 | 4.3E-69 | 6.4E-66 |
| | Clec12a | 0.455 | 0.481 | 0.424 | 1.0E-39 | 1.6E-36 | | Sqstm1 | 0.921 | 0.608 | 0.494 | 1.2E-67 | 1.8E-64 |
| | Colgalt1 | 0.261 | 0.486 | 0.423 | 4.1E-39 | 6.1E-36 | | Emp1 | 0.827 | 0.723 | 0.773 | 6.3E-65 | 9.4E-62 |
| | Cks2 | 0.955 | 0.534 | 0.469 | 1.2E-36 | 1.8E-33 | | Rps2 | 0.628 | 0.867 | 0.830 | 3.3E-63 | 4.9E-60 |
| | Rpn1 | 0.256 | 0.347 | 0.457 | 3.6E-31 | 5.3E-28 | | Hspa1b | 0.866 | 0.659 | 0.604 | 4.8E-63 | 7.2E-60 |
| | Id1 | 0.318 | 0.832 | 0.819 | 1.9E-28 | 2.8E-25 | | Fam20c | 0.379 | 0.847 | 0.869 | 2.1E-60 | 3.1E-57 |
| | Fam20c | 0.332 | 0.792 | 0.904 | 6.0E-27 | 9.0E-24 | | Slc37a2 | 0.582 | 0.609 | 0.576 | 3.5E-55 | 5.3E-52 |
| | Ctsc | 0.283 | 0.661 | 0.629 | 8.2E-26 | 1.2E-22 | | Ctsl | 0.651 | 0.807 | 0.742 | 5.6E-54 | 8.4E-51 |
| | Lamtor4 | 0.498 | 0.641 | 0.610 | 1.3E-25 | 2.0E-22 | | Fabp5 | 0.488 | 0.626 | 0.784 | 3.1E-49 | 4.6E-46 |
| | Csf1r | 0.363 | 0.280 | 0.375 | 2.1E-25 | 3.2E-22 | | Ctsa | 0.516 | 0.748 | 0.695 | 6.2E-49 | 9.3E-46 |
| | Rhoc | 0.353 | 0.529 | 0.648 | 1.6E-24 | 2.4E-21 | | Hsp90aa1 | 1.357 | 0.762 | 0.791 | 1.2E-48 | 1.8E-45 |
| | Cdc42ep3 | 0.331 | 0.512 | 0.462 | 5.0E-23 | 7.5E-20 | | Timp2 | 1.159 | 0.724 | 0.658 | 9.5E-47 | 1.4E-43 |
| | Krt18 | 0.259 | 0.611 | 0.616 | 1.1E-21 | 1.7E-18 | | Hal | 0.601 | 0.514 | 0.776 | 7.5E-29 | 1.1E-25 |
| | Rgs1 | 0.444 | 0.650 | 0.843 | 6.1E-21 | 9.2E-18 | | Cldn1 | 0.594 | 0.582 | 0.669 | 4.4E-28 | 6.6E-25 |
| | H2afz | 0.536 | 0.744 | 0.728 | 1.6E-19 | 2.5E-16 | | Jdp2 | 0.437 | 0.670 | 0.653 | 2.7E-25 | 4.0E-22 |
| | Rpl3 | 0.306 | 0.538 | 0.517 | 1.6E-16 | 2.4E-13 | | Acod1 | 0.845 | 0.548 | 0.530 | 8.1E-24 | 1.2E-20 |
| | Hist1h2ap | 0.291 | 0.508 | 0.492 | 1.7E-16 | 2.5E-13 | | Slc7a11 | 0.925 | 0.565 | 0.555 | 1.8E-23 | 2.7E-20 |
| | Mpeg1 | 0.501 | 0.595 | 0.621 | 2.6E-16 | 3.8E-13 | | Grn | 0.382 | 0.873 | 0.835 | 1.8E-21 | 2.8E-18 |
| | Cfp | 0.260 | 0.543 | 0.504 | 1.5E-15 | 2.2E-12 | | Vegfa | 0.831 | 0.495 | 0.452 | 1.0E-20 | 1.6E-17 |
| | Tubb6 | 0.260 | 0.394 | 0.341 | 4.6E-14 | 7.0E-11 | | Gm26870 | 0.478 | 0.567 | 0.719 | 2.0E-19 | 3.0E-16 |
| | Erp29 | 0.740 | 0.600 | 0.627 | 1.3E-13 | 2.0E-10 | | Gusb | 0.335 | 0.618 | 0.634 | 2.3E-19 | 3.5E-16 |
| | Fyb | 0.264 | 0.751 | 0.780 | 2.7E-10 | 4.0E-07 | | Cd300c2 | 0.375 | 0.675 | 0.645 | 4.3E-19 | 6.5E-16 |
| | Spp1 | 0.341 | 0.831 | 0.843 | 3.7E-09 | 5.5E-06 | | Hspe1 | 0.432 | 0.639 | 0.663 | 1.4E-15 | 2.1E-12 |
| | Atad2 | 0.270 | 0.495 | 0.481 | 4.1E-09 | 6.2E-06 | | Slc43a3 | 0.361 | 0.577 | 0.707 | 1.5E-15 | 2.3E-12 |
| | Bcl2a1d | 0.651 | 0.380 | 0.413 | 7.1E-08 | 1.1E-04 | | Abcg1 | 0.435 | 0.571 | 0.559 | 2.1E-15 | 3.2E-12 |
| | S100a10 | 0.397 | 0.513 | 0.610 | 3.7E-07 | 5.6E-04 | | Syne1 | 0.692 | 0.511 | 0.667 | 2.2E-14 | 3.3E-11 |
| | Ctss | 0.529 | 0.679 | 0.716 | 6.8E-07 | 1.0E-03 | | Rps6ka2 | 0.368 | 0.521 | 0.558 | 4.6E-14 | 6.8E-11 |
| | Apoe | 0.328 | 0.748 | 0.737 | 8.0E-06 | 1.2E-02 | | Ftl1-ps1 | 0.479 | 0.338 | 0.347 | 1.7E-11 | 2.5E-08 |
| | Tnf | 0.345 | 0.327 | 0.311 | 8.5E-06 | 1.3E-02 | | C3 | 0.346 | 0.632 | 0.634 | 4.3E-11 | 6.5E-08 |
| | Gm12840 | 1.062 | 0.321 | 0.302 | 6.5E-05 | 9.7E-02 | | Thbs1 | 1.191 | 0.451 | 0.593 | 1.1E-09 | 1.6E-06 |
| | Pou2f2 | 0.584 | 0.387 | 0.411 | 7.3E-04 | 1.0E+00 | | Hbb-bs | 0.414 | 0.804 | 0.812 | 3.3E-09 | 5.0E-06 |
| | Aprt | 0.360 | 0.639 | 0.770 | 1.4E-03 | 1.0E+00 | | Rpl3 | 0.505 | 0.532 | 0.522 | 5.9E-09 | 8.8E-06 |
| | Abca1 | 0.524 | 0.768 | 0.829 | 4.0E-03 | 1.0E+00 | | Acp5 | 0.601 | 0.473 | 0.461 | 1.5E-03 | 1.0E+00 |
| | Tgfbi | 0.309 | 0.586 | 0.654 | 5.9E-03 | 1.0E+00 | | Hspa1a | 0.817 | 0.527 | 0.564 | 1.7E-03 | 1.0E+00 |
| | Acp5 | 0.644 | 0.451 | 0.471 | 7.0E-03 | 1.0E+00 | | Spp1 | 1.001 | 0.762 | 0.862 | 5.5E-03 | 1.0E+00 |
| **Neu-2** | BC100530 | 1.757 | 0.815 | 0.739 | 0.0E+00 | 0.0E+00 | **Neu-5-IFN** | Isg15 | 3.959 | 0.914 | 0.574 | 8.6E-166 | 1.3E-162 |
| | Gm5483 | 1.712 | 0.938 | 0.789 | 0.0E+00 | 0.0E+00 | | Rsad2 | 3.922 | 0.893 | 0.799 | 5.5E-153 | 8.3E-150 |
| | Wfdc17 | 1.379 | 0.993 | 0.891 | 0.0E+00 | 0.0E+00 | | Ifi47 | 2.686 | 0.858 | 0.765 | 3.1E-121 | 4.7E-118 |
| | Cxcl2 | 1.173 | 0.951 | 0.703 | 0.0E+00 | 0.0E+00 | | Ifitm3 | 2.227 | 0.937 | 0.742 | 1.4E-114 | 2.1E-111 |
| | Ifitm1 | 1.111 | 0.963 | 0.776 | 0.0E+00 | 0.0E+00 | | Gbp2 | 3.371 | 0.883 | 0.701 | 2.0E-112 | 3.0E-109 |
| | Lrg1 | 0.856 | 0.929 | 0.751 | 0.0E+00 | 0.0E+00 | | Ifit2 | 1.589 | 0.827 | 0.776 | 2.5E-109 | 3.8E-106 |
| | Egr1 | 0.840 | 0.912 | 0.633 | 0.0E+00 | 0.0E+00 | | Rtp4 | 2.640 | 0.820 | 0.711 | 1.3E-94 | 1.9E-91 |
| | Retnlg | 0.817 | 0.960 | 0.812 | 0.0E+00 | 0.0E+00 | | Slfn4 | 2.454 | 0.848 | 0.736 | 8.9E-90 | 1.3E-86 |
| | Wfdc21 | 0.764 | 0.945 | 0.788 | 0.0E+00 | 0.0E+00 | | Slfn5 | 2.495 | 0.787 | 0.619 | 1.1E-87 | 1.7E-84 |
| | Ccl6 | 0.733 | 0.910 | 0.693 | 0.0E+00 | 0.0E+00 | | Ifit1bl2 | 0.982 | 0.794 | 0.613 | 2.2E-77 | 3.3E-74 |
| | Lcn2 | 0.710 | 0.911 | 0.736 | 0.0E+00 | 0.0E+00 | | Isg20 | 2.415 | 0.731 | 0.518 | 3.4E-64 | 5.1E-61 |
| | Cxcl3 | 0.666 | 0.840 | 0.353 | 0.0E+00 | 0.0E+00 | | Ifit1 | 3.118 | 0.693 | 0.425 | 1.5E-61 | 2.2E-58 |
| | Tceal9 | 0.526 | 0.888 | 0.729 | 0.0E+00 | 0.0E+00 | | Zbp1 | 1.428 | 0.789 | 0.729 | 2.5E-60 | 3.7E-57 |
| | Slfn4 | 0.469 | 0.942 | 0.649 | 0.0E+00 | 0.0E+00 | | Ifi202b | 0.654 | 0.739 | 0.617 | 3.0E-57 | 4.5E-54 |
| | Id1 | 0.277 | 0.915 | 0.784 | 4.8E-284 | 7.2E-281 | | Ifit3b | 2.094 | 0.698 | 0.535 | 7.2E-57 | 1.1E-53 |
| | Slpi | 0.579 | 0.904 | 0.662 | 4.9E-284 | 7.3E-281 | | Oasl1 | 2.280 | 0.668 | 0.414 | 6.9E-56 | 1.0E-52 |
| | Dgat2 | 0.449 | 0.795 | 0.605 | 5.1E-282 | 7.6E-279 | | Ifit3 | 3.099 | 0.612 | 0.263 | 4.7E-39 | 7.0E-36 |
| | Adam8 | 1.014 | 0.729 | 0.463 | 8.4E-262 | 1.3E-258 | | Cldn1 | 0.258 | 0.777 | 0.646 | 1.1E-38 | 1.6E-35 |
| | Prok2 | 0.277 | 0.790 | 0.700 | 4.3E-239 | 6.4E-236 | | Bst2 | 1.698 | 0.657 | 0.525 | 6.1E-38 | 9.1E-35 |
| | Vim | 0.636 | 0.831 | 0.556 | 1.3E-219 | 2.0E-216 | | Ctss | 1.414 | 0.764 | 0.701 | 7.1E-37 | 1.1E-33 |
| | G0s2 | 0.703 | 0.819 | 0.568 | 3.6E-170 | 5.3E-167 | | Fcgr4 | 0.876 | 0.812 | 0.699 | 1.1E-32 | 1.6E-29 |
| | Glrx | 0.709 | 0.678 | 0.390 | 5.3E-170 | 7.9E-167 | | Cmpk2 | 1.332 | 0.647 | 0.692 | 9.6E-30 | 1.4E-26 |

| | | | | | | |
|---|---|---|---|---|---|---|
| | F630028O10Ri | 0.557 | 0.798 | 0.542 | 6.6E-167 | 9.8E-164 |
| | Ier3 | 0.594 | 0.972 | 0.826 | 5.5E-157 | 8.2E-154 |
| | Steap4 | 0.560 | 0.732 | 0.674 | 8.3E-127 | 1.2E-123 |
| | Gadd45a | 0.551 | 0.731 | 0.616 | 2.4E-122 | 3.6E-119 |
| | Stfa2l1 | 1.367 | 0.682 | 0.553 | 2.0E-118 | 3.0E-115 |
| | Csf2rb | 0.259 | 0.921 | 0.714 | 1.3E-114 | 1.9E-111 |
| | Stfa2 | 0.989 | 0.680 | 0.704 | 2.8E-87 | 4.2E-84 |
| | Il1f9 | 0.250 | 0.718 | 0.540 | 2.8E-75 | 4.3E-72 |
| | Asprv1 | 0.555 | 0.707 | 0.722 | 3.9E-33 | 5.8E-30 |
| | Saa3 | 0.387 | 0.748 | 0.691 | 3.2E-27 | 4.7E-24 |
| | Tacstd2 | 0.305 | 0.584 | 0.468 | 3.7E-24 | 5.5E-21 |
| | Osm | 0.506 | 0.460 | 0.331 | 1.0E-09 | 1.6E-06 |
| | Hacd4 | 0.294 | 0.496 | 0.406 | 4.6E-03 | 1.0E+00 |
| **Neu-4-classic** | Retnlg | 3.329 | 0.993 | 0.845 | 0.0E+00 | 0.0E+00 |
| | Ifitm6 | 3.306 | 0.992 | 0.478 | 0.0E+00 | 0.0E+00 |
| | Lcn2 | 2.827 | 0.999 | 0.771 | 0.0E+00 | 0.0E+00 |
| | Wfdc21 | 2.632 | 0.999 | 0.822 | 0.0E+00 | 0.0E+00 |
| | Mmp8 | 2.622 | 0.971 | 0.633 | 0.0E+00 | 0.0E+00 |
| | Ly6g | 2.187 | 0.944 | 0.567 | 0.0E+00 | 0.0E+00 |
| | Anxa1 | 2.167 | 0.997 | 0.716 | 0.0E+00 | 0.0E+00 |
| | Prok2 | 2.127 | 0.960 | 0.707 | 0.0E+00 | 0.0E+00 |
| | Lrg1 | 1.793 | 0.992 | 0.789 | 0.0E+00 | 0.0E+00 |
| | Ifitm3 | 1.588 | 0.989 | 0.726 | 0.0E+00 | 0.0E+00 |
| | Cd177 | 1.349 | 0.907 | 0.656 | 0.0E+00 | 0.0E+00 |
| | Ggt1 | 1.045 | 0.897 | 0.603 | 0.0E+00 | 0.0E+00 |
| | Glul | 0.436 | 0.853 | 0.387 | 0.0E+00 | 0.0E+00 |
| | Gyg | 0.996 | 0.863 | 0.596 | 8.3E-285 | 1.2E-281 |
| | Wfdc17 | 1.832 | 0.997 | 0.916 | 2.1E-279 | 3.1E-276 |
| | Chil1 | 0.973 | 0.942 | 0.592 | 8.9E-269 | 1.3E-265 |
| | Steap4 | 0.716 | 0.940 | 0.670 | 1.5E-254 | 2.2E-251 |
| | Stfa2 | 1.185 | 0.909 | 0.678 | 2.8E-254 | 4.2E-251 |
| | Tgm1 | 0.493 | 0.865 | 0.635 | 2.0E-252 | 3.0E-249 |
| | Mgst1 | 0.981 | 0.870 | 0.580 | 2.4E-243 | 3.6E-240 |
| | Flna | 0.955 | 0.875 | 0.433 | 4.9E-242 | 7.4E-239 |
| | Ngp | 1.645 | 0.786 | 0.355 | 7.6E-234 | 1.1E-230 |
| | Slpi | 1.162 | 0.978 | 0.715 | 2.2E-232 | 3.3E-229 |
| | Hacd4 | 1.004 | 0.822 | 0.400 | 5.4E-230 | 8.1E-227 |
| | Ifitm1 | 1.769 | 0.992 | 0.819 | 1.3E-203 | 2.0E-200 |
| | Ccl6 | 1.106 | 0.959 | 0.742 | 2.3E-188 | 3.5E-185 |
| | Smpdl3a | 0.852 | 0.826 | 0.558 | 3.0E-187 | 4.5E-184 |
| | Tuba1a | 0.578 | 0.780 | 0.474 | 1.3E-175 | 1.9E-172 |
| | Vim | 0.919 | 0.966 | 0.611 | 2.8E-175 | 4.1E-172 |
| | Syne1 | 0.469 | 0.810 | 0.615 | 7.0E-163 | 1.1E-159 |
| | Tgfbi | 0.673 | 0.922 | 0.604 | 6.7E-159 | 1.0E-155 |
| | Glrx | 0.596 | 0.853 | 0.445 | 1.3E-157 | 2.0E-154 |
| | Tacstd2 | 0.565 | 0.812 | 0.476 | 1.1E-141 | 1.7E-138 |
| | Pi16 | 0.611 | 0.785 | 0.543 | 5.7E-120 | 8.6E-117 |
| | Lyz2 | 0.426 | 0.978 | 0.755 | 2.3E-116 | 3.4E-113 |
| | Lmo4 | 0.573 | 0.715 | 0.315 | 5.2E-112 | 7.9E-109 |
| | Ly6c2 | 0.617 | 0.708 | 0.439 | 1.4E-109 | 2.1E-106 |
| | BC100530 | 1.364 | 0.877 | 0.752 | 3.0E-94 | 4.5E-91 |
| | Aldh2 | 0.324 | 0.698 | 0.433 | 2.9E-86 | 4.3E-83 |
| | Stfa2l1 | 0.440 | 0.835 | 0.571 | 5.3E-68 | 7.9E-65 |
| | Ceacam1 | 0.277 | 0.709 | 0.530 | 2.0E-64 | 3.0E-61 |
| | Plac8 | 0.327 | 0.668 | 0.476 | 5.6E-64 | 8.4E-61 |
| | Acvrl1 | 0.598 | 0.673 | 0.666 | 1.5E-62 | 2.3E-59 |
| | Camp | 1.496 | 0.597 | 0.402 | 1.9E-49 | 2.9E-46 |
| | Serpinb1a | 0.774 | 0.616 | 0.519 | 1.8E-45 | 2.7E-42 |
| | Abcd2 | 0.289 | 0.621 | 0.535 | 1.2E-40 | 1.8E-37 |
| | Stfa3 | 0.395 | 0.382 | 0.646 | 1.5E-34 | 2.3E-31 |
| | Sept9 | 0.260 | 0.546 | 0.375 | 1.8E-28 | 2.7E-25 |
| | Olfm4 | 0.922 | 0.683 | 0.609 | 5.7E-22 | 8.6E-19 |
| | Asprv1 | 0.255 | 0.706 | 0.718 | 2.2E-13 | 3.3E-10 |
| | C130026I21Rik | 0.253 | 0.399 | 0.417 | 2.3E-11 | 3.5E-08 |

| | | | | | | |
|---|---|---|---|---|---|---|
| | Clec4a3 | 0.377 | 0.614 | 0.450 | 1.1E-28 | 1.6E-25 |
| | Acod1 | 0.910 | 0.764 | 0.528 | 9.0E-26 | 1.3E-22 |
| | Il18 | 0.463 | 0.556 | 0.366 | 4.8E-23 | 7.2E-20 |
| | Fxyd3 | 0.255 | 0.571 | 0.443 | 7.7E-23 | 1.2E-19 |
| | Irf7 | 1.505 | 0.642 | 0.564 | 5.3E-21 | 7.9E-18 |
| | Il18bp | 0.433 | 0.508 | 0.318 | 7.1E-20 | 1.1E-16 |
| | Cxcl10 | 1.827 | 0.551 | 0.388 | 8.9E-20 | 1.3E-16 |
| | Lair1 | 0.305 | 0.609 | 0.516 | 8.0E-19 | 1.2E-15 |
| | Gm4316 | 0.443 | 0.279 | 0.343 | 1.0E-17 | 1.5E-14 |
| | Npc2 | 0.571 | 0.904 | 0.883 | 1.2E-17 | 1.9E-14 |
| | Gbp5 | 1.547 | 0.660 | 0.801 | 1.3E-17 | 1.9E-14 |
| | AW112010 | 0.469 | 0.520 | 0.370 | 1.3E-17 | 1.9E-14 |
| | Cd274 | 0.977 | 0.736 | 0.682 | 3.9E-16 | 5.9E-13 |
| | Hes1 | 0.375 | 0.358 | 0.549 | 4.7E-16 | 7.0E-13 |
| | Ifi207 | 0.258 | 0.655 | 0.691 | 6.5E-15 | 9.7E-12 |
| | Usp18 | 1.418 | 0.604 | 0.700 | 7.7E-15 | 1.2E-11 |
| | Abcg1 | 0.446 | 0.701 | 0.558 | 1.4E-13 | 2.2E-10 |
| | Fyb | 0.553 | 0.835 | 0.768 | 1.2E-12 | 1.8E-09 |
| | Bcl2a1b | 0.420 | 0.751 | 0.586 | 9.3E-11 | 1.4E-07 |
| | Cxcl9 | 0.426 | 0.736 | 0.737 | 1.9E-08 | 2.8E-05 |
| | Ifi27l2a | 2.036 | 0.553 | 0.506 | 1.4E-07 | 2.1E-04 |
| | Cybb | 0.284 | 0.546 | 0.446 | 3.7E-07 | 5.5E-04 |
| | Ly6i | 0.647 | 0.586 | 0.648 | 9.8E-07 | 1.5E-03 |
| | Plac8 | 1.572 | 0.533 | 0.491 | 1.0E-06 | 1.6E-03 |
| | Mpeg1 | 0.339 | 0.726 | 0.608 | 2.2E-06 | 3.4E-03 |
| | Emp3 | 0.292 | 0.447 | 0.314 | 2.4E-06 | 3.5E-03 |
| | Gm20234 | 0.310 | 0.609 | 0.679 | 2.7E-06 | 4.1E-03 |
| | Cst3 | 0.308 | 0.858 | 0.835 | 9.4E-06 | 1.4E-02 |
| | Ddx60 | 1.381 | 0.528 | 0.576 | 2.6E-05 | 3.9E-02 |
| | Ifi209 | 1.094 | 0.563 | 0.675 | 6.0E-05 | 9.0E-02 |
| | Unc93b1 | 0.390 | 0.569 | 0.519 | 6.2E-05 | 9.3E-02 |
| | Hba-a1 | 0.963 | 0.429 | 0.553 | 1.2E-04 | 1.7E-01 |
| | Lyz2 | 0.310 | 0.838 | 0.771 | 1.9E-03 | 1.0E+00 |
| | Ccl4 | 0.628 | 0.921 | 0.920 | 2.1E-03 | 1.0E+00 |
| | Ly6c2 | 1.048 | 0.482 | 0.460 | 3.0E-03 | 1.0E+00 |

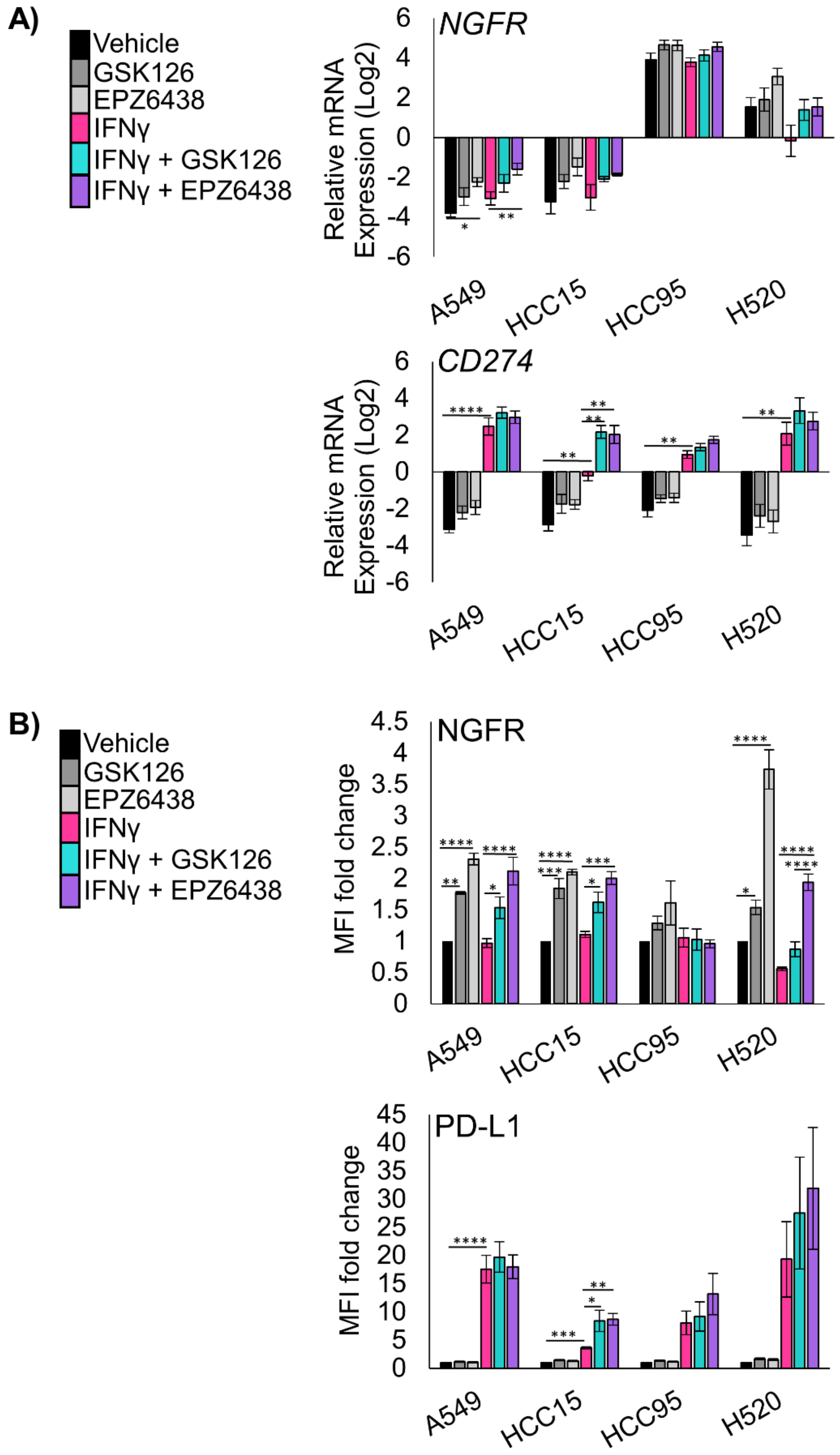


**Supplementary Figure 1: Related to Figure 1**

**A)** RT-qPCR in the indicated four human lung cancer cell lines treated for 7 days with vehicle or EZH2 inhibition with IFNγ added on day 5 for the genes *NGFR* and *CD274*, mean +/- SEM is graphed, n = 4 individual experiments, * indicated p=0.0481 **p<0.0098, ****p<0.0001 by one-way ANOVA with pairwise comparisons and Holm-Šídák's *post hoc* test. **B)** Flow cytometry analysis of indicated four human lung cancer cell lines treated for 6 days with vehicle or EZH2 inhibition with IFNγ added on day 5 for the cell surface proteins NGFR and PD-L1, mean +/- SEM is graphed, n = 4 individual experiments, * indicated $p = 0.04$, **$p=0.0033$, ***$p<0.0003$, ****$p<0.0001$ by one-way ANOVA with pairwise comparisons and Holm-Šídák's *post hoc* test.

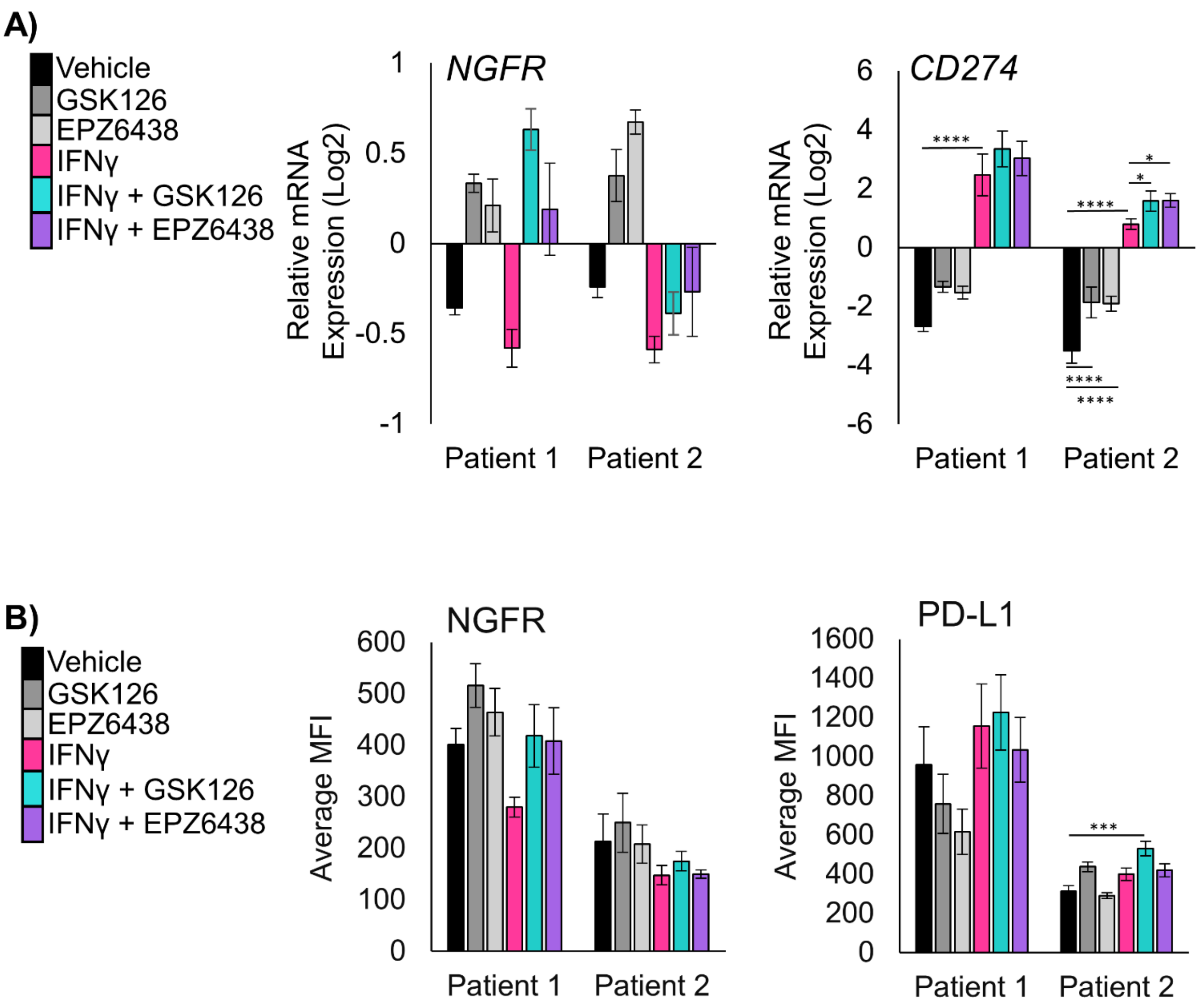


**Supplementary Figure 2: Related to Figure 2**

**A)** RT-qPCR in the indicated two unique patient-derived tumoroid cultures treated for 11 days and IFNγ added on day 9 for the genes *NGFR* and *CD274*, mean +/- SEM is graphed, n = 4 individual experiments, * indicates p<0.03, ****p<0.0001 by one-way ANOVA with pairwise comparisons and Holm-Šídák's *post hoc* test. **B)** Flow cytometry analysis of indicated two unique patient derived tumoroid cultures treated for 11 days and IFNγ added on day 9 for the cell surface proteins NGFR and PD-L1, mean +/- SEM is graphed, n = 4 individual experiments, *** indicates p=0.0002 by one-way ANOVA with pairwise comparisons and Holm-Šídák's *post hoc* test.

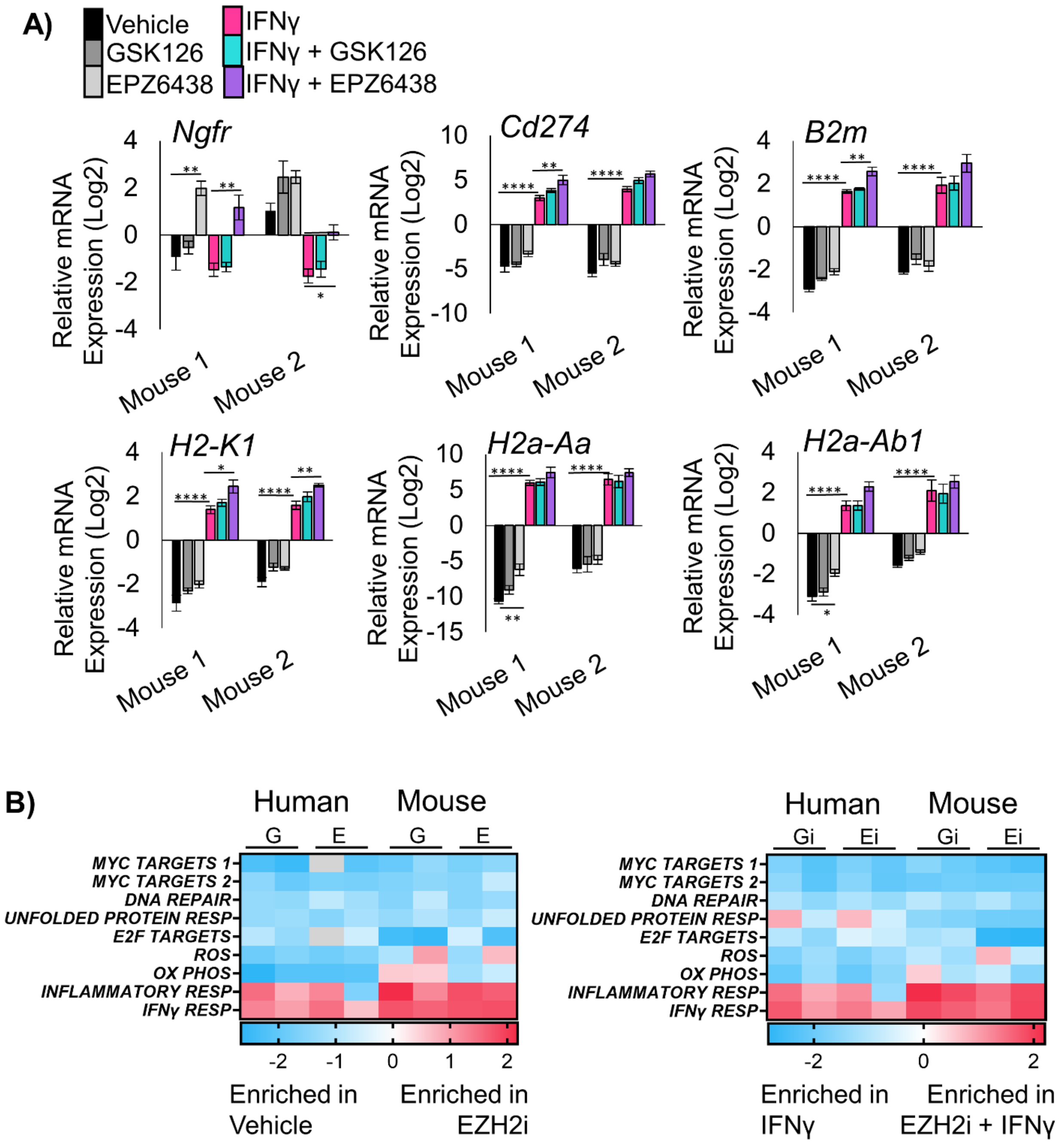


**Supplementary Figure 3: Related to Figure 3**

**A)** RT-qPCR in the indicated two unique murine tumoroid cultures treated for 11 days and IFNγ added on day 9 for the genes *Ngfr, B2m, H2-K1, Cd274,* and *H2a-Aa*, mean +/- SEM is graphed, n = 5 individual experiments for mouse 1, n=4 individual experiments for mouse 2, * indicates p<0.05, **p<0.009, ****p<0.0001 by one-way ANOVA with pairwise comparisons and Holm-Šídák's *post hoc* test. **B)** Heat map of Enrichment Scores using Gene Set Enrichment Analysis on human or murine tumoroids treated with the EZH2 inhibitors GSK126 (G) or EPZ6438 (E) contrasted to vehicle control, or treated with EZH2 inhibitor and IFNγ contrasted to IFNγ alone. See also Supp. Table 2.

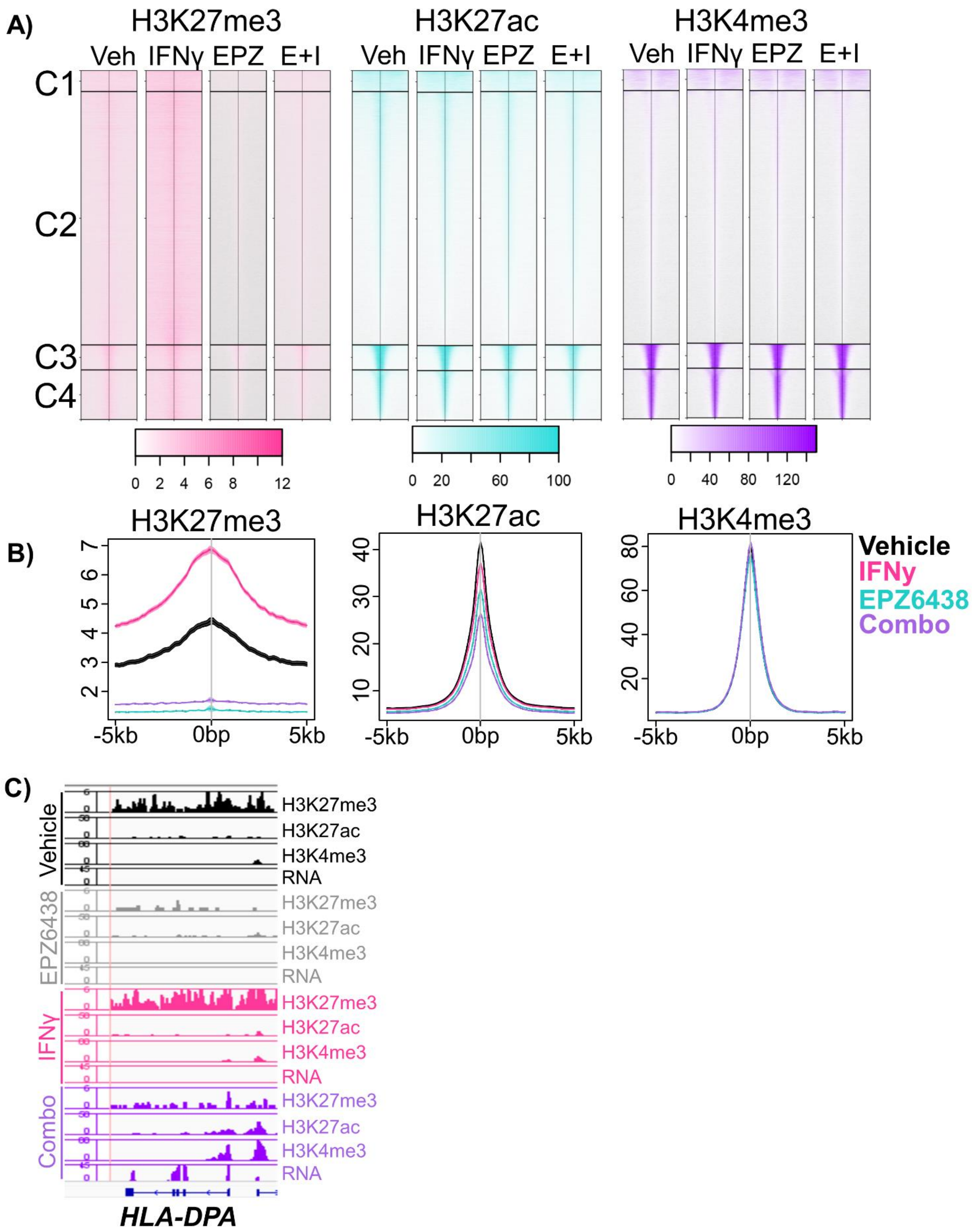


**Supplementary Figure 4: Related to Figure 4**

**A)** Heatmap representation of H3K27me3-, H3K27ac- and H3K4me3-bound chromatin peaks centered across a ±5kb window that shows occupancy in tumoroids cultures of the indicated treatments. **B)** Histogram of merged

ChIP-seq peaks in the indicated 3D tumoroid samples. **C)** Wiggle plots for H3K27me3, H3K27ac, and H3K4me3 histone mark enrichments, and matched RNAseq tracks in patient-derived tumoroids from the indicated treatment groups for the gene *HLA-DPA*.

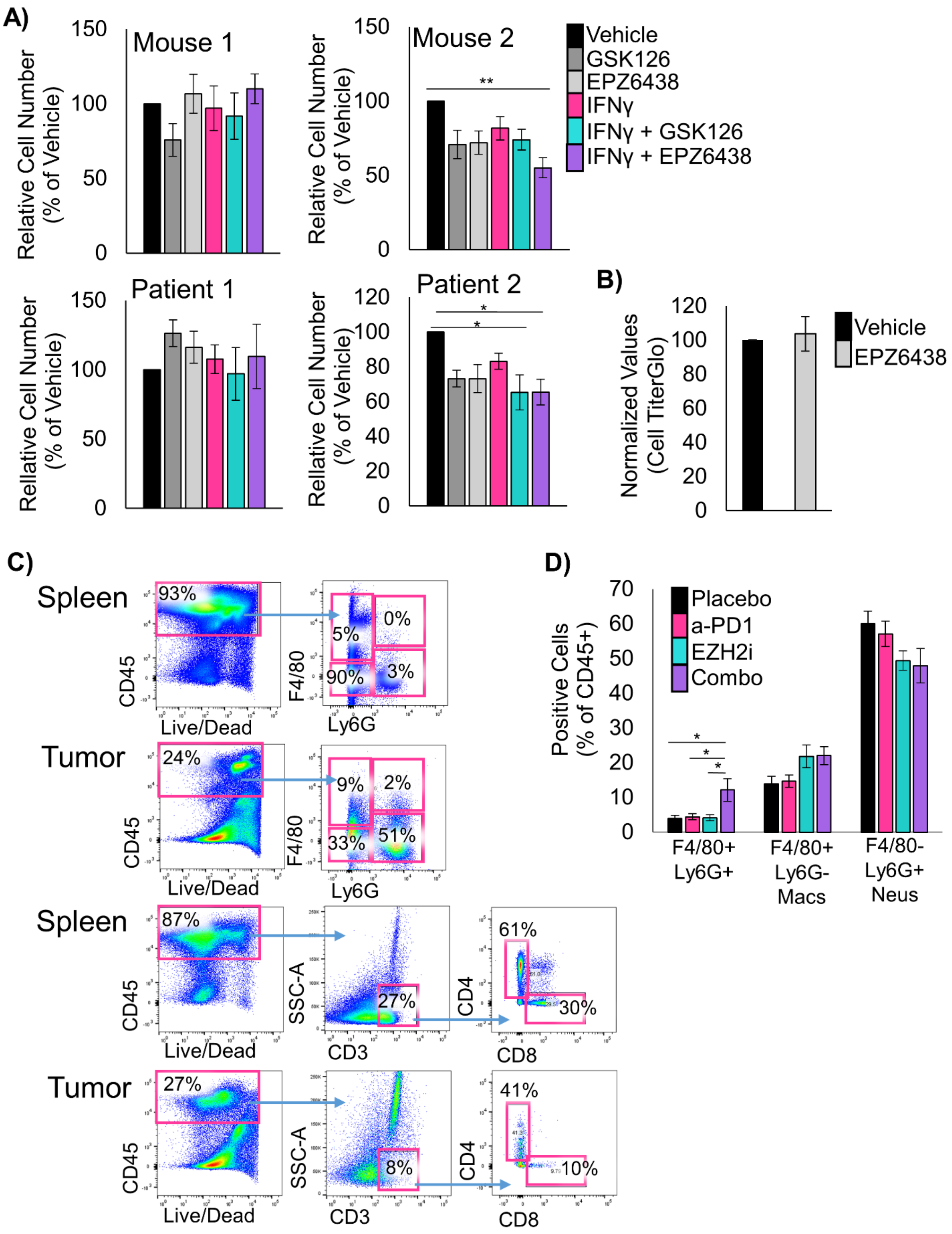


**Supplementary Figure 5: Related to Figure 5**

**A)** Relative counts of cells after dissociation of tumoroids treated with the indicated treatments, mean +/- s.e.m. is plotted, n=4 individual experiments for the human tumoroids, n=7 experiments for mouse 1 and n=5 experiments for mouse 2, * indicates p<0.0250, **p=0.0048 by one-way ANOVA with multiple comparisons and

Holm-Šídák's *post-hoc* test. **B)** Relative luminescence from CellTiter Glo assayed tumoroid cultures treated with vehicle or EPZ6438, mean +/- s.e.m. is plotted, n=4 individual experiments. **C)** Representative flow cytometry plots with the indicated samples and markers. The left plots are gated on FSC/SSC and Live/Dead negative cells. Spleen samples are shown as staining controls. **D)** Flow cytometry analysis of dissociated tumors from the syngeneic grafts from the indicated treatment arms at day 14. Percentage of CD45+ cells expressing F4/80 or Ly6G are graphed, mean +/- s.e.m. is plotted, placebo n=8, EZH2 inhibitor n=8, anti-PD1 n=9, combo n=7, * indicates p<0.012, by one-way ANOVA with multiple comparisons and Holm-Šídák's *post-hoc* test.

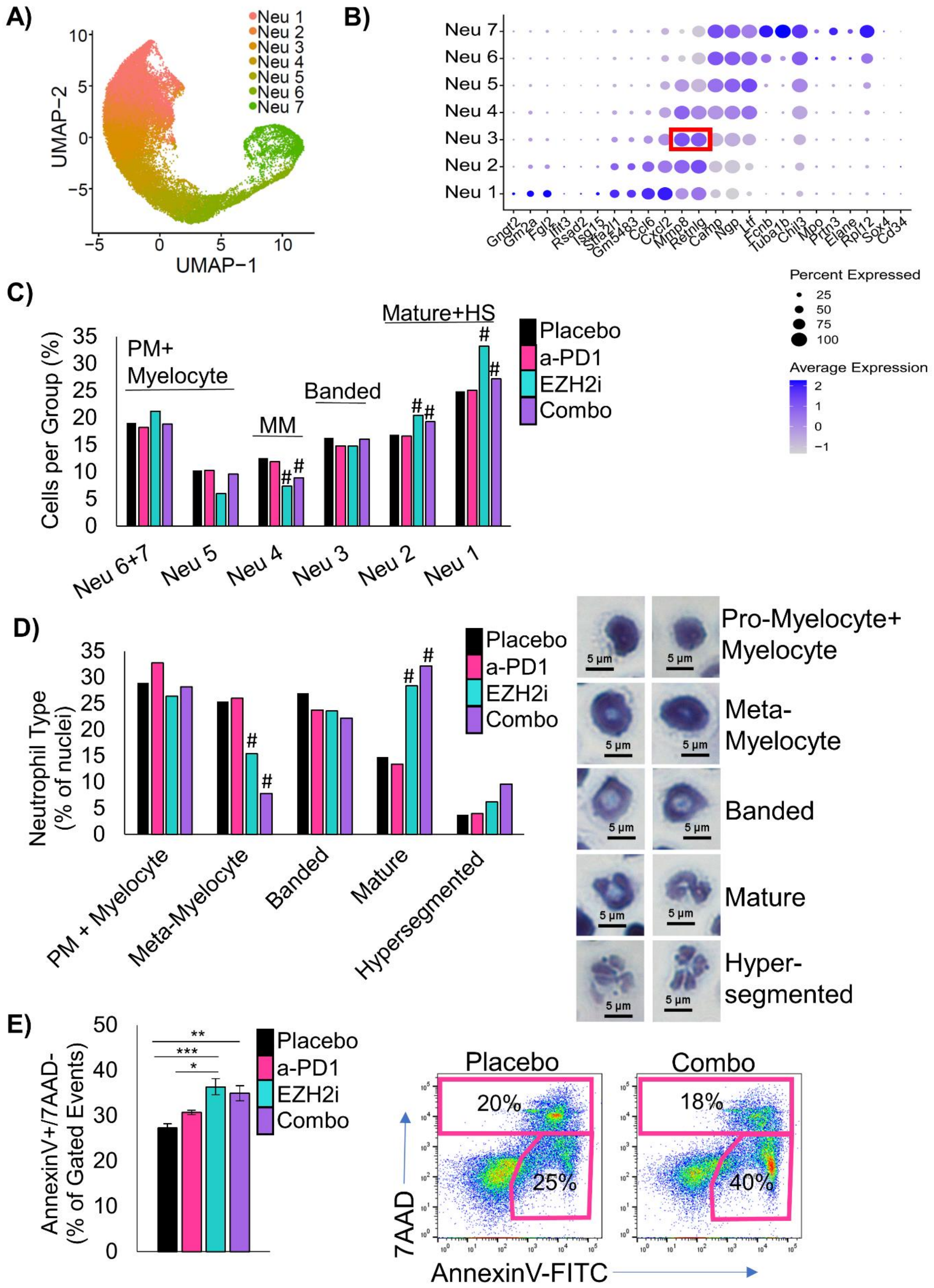


**Supplementary Figure 6: Related to Figure 6**

**A)** Annotated Uniform Manifold Approximation and Projection (UMAP) plot showing the 8 different populations of neutrophils within the bone marrow of tumor-bearing mice treated with placebo, GSK126, anti-PD1, or combined GSK126 with anti-PD1. **B)** Dot plot showing the relative expression of marker genes (x-axis) in each neutrophil cluster (y-axis). Expression of *Mmp8* and *Rentlg* were shown to be enriched in banded neutrophils. **C)** Percentage of cells per treatment group graphed for the neutrophil populations, # indicates adjusted $p<0.008$ by proportion z-test. **D)** Proportions of different nuclear morphologies in bone marrow cytospins, average of n=2 samples for Placebo, n=1 sample for others, 500 nuclei were counted, # indicates $p<0.0004$ with Fisher's Exact test between Vehicle and EZH2 inhibitor or Vehicle and Placebo for meta-myelocyte vs mature neutrophils. Representative images of nuclei types shown, scale bar = 5µm. **E)** Percentage of AnnexinV+/7AAD- cells in bone marrow cultures 48 hours post isolation from mice treated with the indicated therapies, n=3 biological replicates each with 2 experimental replicates for placebo, EPZ6438 and combo, n=4 biological replicates each with 2 experimental replicates for anti-PD1, * indicates p=0.012, **p=0.0017, ***p= 0.0003 by one-way ANOVA with pairwise comparisons and Holm-Šídák's *post-hoc* test. Representative flow plots of placebo and combination treated cultures shown.
